# cis-γ-Amino-L-proline peptides as chemical probes of amyloidogenic processing in neurons and APP/PS1 mice

**DOI:** 10.64898/2026.04.17.719160

**Authors:** Dayaneth Jácome, Marina Pérez-Palau, Inés Martínez-Soria, Laia Lidón, Cristina Vergara, Daniel Carbajo, Ximena Pulido, Macarena Sánchez-Navarro, Ernest Giralt, Fernando Albericio, Miriam Royo, Rosalina Gavín, José Antonio Del Río

**Author notes:** Shared correspondence: Dr. R. Gavín, MCN lab, Institute for Bioengineering of Catalonia (IBEC). Baldiri i Reixac 15-21, 08028 Barcelona, Spain, Prof. J. A. del Río, MCN lab, Institute for Bioengineering of Catalonia (IBEC) Baldiri i Reixac 15-21, 08028 Barcelona, Spain, Dr. M. Royo, Institute for Advanced Chemistry of Catalonia (IQAC-CSIC) Jordi Girona 18-26, 08034 Barcelona, Spain. Memory Unit, Neurology Department, Hospital de Sant Pau, 08041 Barcelona, Spain (Laia Lidón). R&D Manager at Minoryx Therapeutics, 6041 Gosselies, Walloon Reguin, Belgium (Cristina Vergara). Institute for Advanced Chemistry of Catalonia (IQAC-CSIC), 08034 Barcelona, Spain. Departamento de Química, Universidad del Tolima, 730006299 Ibagué, Colombia (Ximena Pulido). Instituto de Parasitología y Biomedicina López-Neyra (IPB-CSIC), Granada, Spain.

## Abstract

Alzheimer’s disease (AD) is characterized by the accumulation of amyloid-β (Aβ) peptides, which are a key factor in its pathogenesis. In this study, we present the design and evaluation of γ-amino-L-proline peptides as metabolically stable, cell-penetrating molecules that can modulate amyloidogenic processing. We screened a library of γ-peptides in primary neuronal cultures to determine their effects on endogenous Aβ1-42 production, cytotoxicity, and β-secretase (BACE1) activity. Comparative analysis of structurally related analogues enabled the identification of molecular features associated with Aβ-lowering activity, establishing a qualitative structure–activity relationship. Peptide 33 (**P33**) emerged as a lead candidate, selectively reducing BACE1 activity without significantly inhibiting the homologous enzyme, BACE2. *In vitro* blood-brain barrier (BBB) assays revealed that **P33** exhibits favorable transendothelial permeability. Intraperitoneal administration of **P33** in APP/PS1 mice decreased Aβ levels, reduced amyloid plaque burden, and improved performance in a behavioral recognition task without inducing cytotoxicity or systemic toxicity. These results define cis-γ-amino-L-proline peptides as a bioorganically distinct and modular scaffold for the development of intracellular modulators of Aβ production.

**Highlights:**

- γLJAminoLJLLJproline peptides as metabolically stable modulators of Aβ production.
- P33 showed BBB permeability and BACE1 inhibition in primary cortical neurons.
- In APP/PS1 mice, P33 lowers amyloid burden and improves cognition.
- P33 shows good biocompatibility, supporting its therapeutic potential in AD

## 1. Introduction

### 1.1 Aβ peptide as a therapeutic target in Alzheimer’s disease

Alzheimer’s disease (AD) is a progressive neurodegenerative disorder characterized by a gradual decline in cognitive function that ultimately leads to dementia and neuronal loss. With a few exceptions associated with rapidly progressing forms of the disease [1–3], AD typically develops over decades. Neuropathologically, the disease is defined by the presence of extracellular senile plaques and intracellular neurofibrillary tangles (NFTs) in the brain. Senile plaques are composed of aggregated amyloid-β (Aβ) peptides, while NFTs contain hyperphosphorylated forms of the microtubule-associated protein tau [4]. These lesions appear in a stereotyped spatiotemporal pattern that correlates with clinical progression, ranging from preclinical and asymptomatic stages (Braak I–II) to prodromal phases or mild cognitive impairment (Braak III–IV) to symptomatic AD dementia (Braak V–VI) [5,6].

Genetic and experimental evidence strongly suggest that Aβ plays a central role in the pathogenesis of Alzheimer’s disease (AD). Mutations in the amyloid precursor protein (APP) or in enzymes involved in its proteolytic processing, such as β-site APP-cleaving enzyme 1 (BACE1) and components of the γ-secretase complex, can induce amyloid pathology and cognitive impairment in transgenic mouse models, including APP/PS1 and J20 mice. These observations support the amyloid cascade hypothesis, which proposes that excessive Aβ production or impaired clearance triggers a cascade of events that lead to synaptic dysfunction, neurodegeneration, and dementia. According to this framework, therapeutic benefit is expected only if Aβ levels are reduced below a critical threshold early in the disease process, before extensive neurodegeneration occurs (typically beyond Braak stage III [7]).

Consistent with this view, most disease-modifying clinical trials conducted over the past decade have focused on components of the amyloid pathway [8–10]. Recent advances have led to the clinical approval of monoclonal antibodies that target aggregated forms of Aβ. These include Aducanumab, Lecanemab, and Donanemab, which differ in their selectivity for fibrillar and protofibrillar Aβ species, or N-terminally pyroglutamated (pGlu Aβ), which is present in mature plaques [11–13]. Although these agents have demonstrated the ability to reduce amyloid burden and modestly slow cognitive decline in early AD, their clinical efficacy is limited, and safety concerns persist, including amyloid-related imaging abnormalities such as vasogenic edema and cerebral microhemorrhages [14,15]. Moreover, amyloid-targeting immunotherapies have shown little benefit in advanced stages of the disease (Braak V-VI; ICD-10; NINCDS-ADRDA score) [8], highlighting the need for alternative or complementary therapeutic strategies that can modulate amyloidogenic processes with improved safety and broader applicability.

In parallel with immunotherapeutic approaches, extensive efforts have been dedicated to developing small-molecule modulators of amyloidogenic processing, including β-secretase inhibitors [16–19]. Despite promising preclinical results, many of these compounds have failed in clinical trials due to limited brain exposure, off-target effects, or mechanism-based toxicity [20–22]. These failures highlight the need for alternative chemical modalities that can modulate intracellular processes with improved selectivity and tolerability.

### 1.2. Cell-penetrating **γ**-peptides as chemical modulators of amyloidogenic processing

Small molecules and peptides that interfere with amyloidogenic processing of APP represent an attractive alternative to antibody-based therapies. In this context, cell-penetrating peptides (CPPs) have emerged as versatile tools for intracellular delivery and modulation of cellular pathways. CPPs can efficiently cross biological membranes and exhibit favorable biocompatibility profiles [23–26]; however, their widespread application is often limited by poor metabolic stability and rapid proteolytic degradation [27]. Non-natural peptide scaffolds that combine metabolic stability with cell-penetrating properties are an appealing strategy to modulate disease-relevant intracellular pathways.

To address these limitations, a class of γ-peptides based on cis-γ-amino-L-proline oligomers has been developed. These γ-peptides are functionalized at the α-amine of each residue with a natural amino acid side-chain mimetic moiety, resulting in sequences with defined charge distribution and amphiphilicity [28]. These γ-peptides are highly resistant to enzymatic degradation and, in several cases, exhibit cell-penetrating properties comparable to those of the HIV-1 TAT peptide, one of the most extensively studied CPPs [28–33]. The modularity of this scaffold allows for systematic variation in charge, side-chain composition, and spatial patterns, making it well-suited for structure-activity relationship (SAR) studies in complex biological systems.

Here, we present a phenotypic screening approach to identify γ-peptides capable of modulating Aβ production in neuronal systems. We evaluated a library of cis-γ-amino-L-proline-derived peptides in primary neuronal cultures to determine their effects on endogenous Aβ release, cytotoxicity, and selected amyloidogenic readouts. Comparative analysis of structurally related analogues revealed key molecular features associated with Aβ-lowering activity, establishing a qualitative structure-activity relationship. Peptide 33 (P33) emerged as a lead candidate and was evaluated further in exploratory *in vivo* studies using an APP/PS1 mouse model of AD. Together, these results establish γ-peptides as a distinct and modular chemical platform for developing metabolically stable, cell-penetrating agents that can modulate amyloidogenic processing.

## 2. Experimental section

### 2.1. General chemistry methods

The library of γ-peptides with potential CPP abilities was provided by the IQAC-CSIC group. Its synthesis was carried out in solid-phase and will be reported elsewhere. All reagents and solvents were used as purchased from commercial suppliers without further purification. Reactions were monitored by HPLC-MS with a Waters 2795 HPLC coupled to a Waters 2996 UV/Vis detector and a Micromass ZQ mass spectrometer. Intermediate products were purified by reverse-phase automated flash column chromatography with a CombiFlash Rf+ from Teledyne ISCO and using C18 columns (HCOOH or TFA were used as additives when indicated). The final product was purified by semipreparative HPLC with a Waters 1525 HPLC coupled to a Waters 2489 UV/Vis detector using a XBridge® Prep C18 column (5 µM OBDTM, 19 × 100 mm). NMR spectra were obtained with a Bruker Avance Neo 400 MHz spectrometer using the indicated deuterated solvent. Chemical shifts (δ) are given in parts per million (ppm) and referenced to the residual solvent as the internal standard (δH 7.26, δC 77.2 for CDCl3 and δH 3.31, δC 49.0 for MeOD). Signals are reported as follows: s, singlet; d, doublet; t, triplet; m, multiplet; and br, broad. Coupling constants (J) are measured in hertz (Hz). Two-dimensional techniques (HSQC and HMBC) were used to assist with structure elucidation. HPLC chromatograms were obtained with a Waters ARC HPLC coupled to a Waters 2998 PDA detector, using a XBridge® C18 column (3.5 µM, 4.6 × 100 mm). MALDI-TOF MS spectra were obtained with a Bruker Daltonics Autoflex III smartbeam mass spectrometer, using α-cyano-4-hydroxycinnamic acid as the coating matrix.

### 2.2. Synthesis of P33 (1)

#### Methyl (2S,4S)-4-((tert-butoxycarbonyl)amino)-1-isopentylpyrrolidine-2-carboxylate (Boc-NH-g-Amp(ipen)-COOMe, 2)

Methyl (2*S*,4*S*)-4-((tert-butoxycarbonyl)amino)pyrrolidine-2-carboxylate (1.00 g, 3.6 mmol) was dissolved in DCE (20 mL) and the DIPEA was added until pH _ 5. Then isovaleraldehyde (590 µL, 5.3 mmol) was added and the mixture was stirred for 10 min at rt. Then, NaBH_3_CN (335 mg, 5.3 mmol) was added and the mixture was stirred for 16 h at rt. Further isovaleraldehyde (290 µL, 2.7 mmol) and NaBH_3_CN (168 mg, 2.7 mmol) were added, and the mixture was stirred for 4 h at rt. The reaction mixture was quenched with MeOH (2 mL) and partitioned between brine and DCM (40 ml each). The aqueous phase was further extracted with DCM (40 mL). The combined organic layers combined were dried with anhydrous MgSO_4_ and evaporated. The residue was purified by automatic reverse flash column chromatography (from 40% to 60% ACN in H_2_O). The title product (**2**) (601 mg, 1.9 mmol, 54% yield) was obtained as a white solid. **^1^H NMR (400 MHz, CDCl_3_)** δ 5.42 (d, *J* = 8.1 Hz, 1H), 4.28 – 4.18 (m, 1H), 3.73 (s, 3H), 3.18 (dd, *J* = 9.7, 4.8 Hz, 1H), 2.96 (d, *J* = 9.6 Hz, 1H), 2.67 (td, *J* = 10.8, 6.4 Hz, 1H), 2.60 (dd, *J* = 9.7, 5.6 Hz, 1H), 2.42 (ddt, *J* = 16.8, 12.1, 5.3 Hz, 2H), 1.83 – 1.74 (m, 1H), 1.58 (dp, *J* = 13.3, 6.7 Hz, 1H), 1.42 (s, 9H), 1.42 – 1.24 (m, 2H), 0.88 (d, *J* = 6.6 Hz, 3H), 0.87 (d, *J* = 6.6 Hz, 3H). **^13^C NMR (101 MHz, CDCl_3_)** δ 175.0, 155.6, 79.3, 64.8, 60.3, 53.1, 52.2, 49.6, 37.3, 37.0, 28.6, 26.4, 22.9, 22.5. **LC-MS (ESI)** *m/z* [M+H]^+^ 315.5.

#### (2S,4S)-4-((tert-Butoxycarbonyl)amino)-1-isopentylpyrrolidine-2-carboxylic acid (Boc-NH-g-Amp(ipen)-COOH, 3)

An aqueous solution of LiOH 0.1 M (19 mL, 1.9 mmol) was added to a solution of **2** (400 mg, 1.27 mmol) in THF (65 mL). The mixture was stirred at rt for 2 h and quenched with an aqueous solution of HCl 1 M (1.9 mL). The volatiles were evaporated and the residue was purified by reverse flash column chromatography (from 0 to 100% ACN in H_2_O) to obtain **3** (355 mg, 1.18 mmol, 93% yield) as a white solid. **^1^H NMR (400 MHz, MeOD)** δ 4.32 – 4.23 (m, 1H), 3.96 (t, *J* = 8.6 Hz, 1H), 3.63 (dd, *J* = 12.0, 4.0 Hz, 1H), 3.40 (dd, *J* = 12.0, 6.9 Hz, 1H), 3.37 – 3.28 (m, 1H), 3.18 – 3.09 (m, 1H), 2.75 (ddd, *J* = 13.8, 9.0, 7.2 Hz, 1H), 2.08 (ddd, *J* = 13.8, 8.0, 5.9 Hz, 1H), 1.74 – 1.57 (m, 3H), 1.44 (s, 9H), 0.97 (d, *J* = 6.4 Hz, 6H). **^13^C NMR (101 MHz, MeOD)** δ 172.5, 157.7, 80.9, 69.6, 60.2, 55.7, 50.3, 36.2, 35.3, 28.6, 27.2, 22.7, 22.4. **LC-MS (ESI)** *m/z* [M+H]^+^ 301.4.

#### Methyl (2S,4S)-4-((tert-butoxycarbonyl)amino)-1-phenethylpyrrolidine-2-carboxylate (Boc-NH-g-Amp(phet)-COOMe, 4)

Methyl (2*S*,4*S*)-4-((tert-butoxycarbonyl)amino)pyrrolidine-2-carboxylate (1.00 g, 3.6 mmol) was dissolved in DCE (20 mL) and the DIPEA was added until pH _ 5. Then phenylacetaldehyde (840 µL, 7.2 mmol) was added and the mixture was stirred for 10 min at rt. Then, NaBH_3_CN (452 mg, 7.2 mmol) was added and the mixture was stirred for 16 h at rt. The reaction mixture was quenched with MeOH (2 mL) and partitioned between brine and DCM (40 ml each). The aqueous phase was further extracted with DCM (40 mL). The combined organic layers combined were dried with anhydrous MgSO_4_ and evaporated. The residue was purified by automatic reverse flash column chromatography (from 40% to 65% ACN in H_2_O). The title compound (**4**) (783 mg, 2.2 mmol, 63% yield) was obtained as a white solid. **^1^H NMR (400 MHz, CDCl_3_)** δ 7.33 – 7.23 (m, 2H), 7.24 – 7.15 (m, 3H), 5.44 (d, *J* = 7.9 Hz, 1H), 4.35 – 4.19 (m, 1H), 3.71 (s, 3H), 3.29 (dd, *J* = 9.6, 4.5 Hz, 1H), 3.04 (d, *J* = 10.3 Hz, 1H), 2.98 – 2.88 (m, 1H), 2.82 – 2.65 (m, 4H), 2.47 (dt, *J* = 13.8, 8.8 Hz, 1H), 1.88 – 1.77 (m, 1H), 1.44 (s, 9H). **^13^C NMR (101 MHz, CDCl_3_)** δ 174.8, 155.6, 139.9, 128.8, 128.5, 126.3, 79.4, 64.5, 60.2, 56.3, 52.2, 49.7, 36.9, 35.0, 28.6. **LC-MS (ESI)** *m/z* [M+H]^+^ 349.4.

#### Methyl (2*S*,4*S*)-4-amino-1-phenethylpyrrolidine-2-carboxylate, hydrochloride salt (HCl.NH_2_-γ-Amp(phet)-COOMe, 5)

A 4M HCl solution in dioxane (5 mL) was added to a solution of **4** (450 mg, 1.29 mmol) in dioxane (5 mL). The mixture was stirred for 1.5 h and the volatiles evaporated. The residue was dissolved in H_2_O/ACN 1:1 and lyophilized to obtain **5** (440 mg, 1.29 mmol, quantitative yield) as a white solid. The product was used in the next step without further purification. **^1^H NMR (400 MHz, MeOD)** δ 7.40 – 7.31 (m, 4H), 7.31 – 7.24 (m, 1H), 4.72 (dd, *J* = 11.1, 8.2 Hz, 1H), 4.38 – 4.26 (m, 1H), 4.01 (dd, *J* = 13.0, 4.9 Hz, 1H), 3.90 (s, 3H), 3.90 – 3.81 (m, 2H), 3.57 – 3.49 (m, 1H), 3.21 – 3.14 (m, 2H), 3.08 (dt, *J* = 14.0, 8.2 Hz, 1H), 2.42 (ddd, *J* = 14.0, 11.1, 7.5 Hz, 1H). **^13^C NMR (101 MHz, MeOD)** δ 168.1, 137.2, 130.0, 129.9, 128.5, 67.4, 58.2, 57.4, 54.4, 48.3, 33.5, 32.8. **LC-MS (ESI)** *m/z* [M+H]^+^ 249.3.

#### Methyl (2*S*,4*S*)-4-[(2*S*,4*S*)-4-((tert-butoxycarbonyl)amino)-1-isopentylpyrrolidine-2-carboxamido]-1-phenethylpyrrolidine-2-carboxylate (Boc-NH-γ-Amp(ipen)-Amp(phet)-COOMe, 6)

DIPEA (180 µL, 1.04 mmol) was added to a mixture of **3** (240 mg, 0.80 mmol), **5** (296 mg, 1.04 mmol), HOBt·H_2_O (147 mg, 0.96 mmol) and EDC·HCl (230 mg, 1.2 mmol) in DMF (10 mL). The mixture was stirred at rt for 16 h and evaporated. The residue was dissolved in DCM (20 mL) and washed with a saturated solution of NaHCO_3_ (20 mL), a 0.5% citric acid solution (20 mL) and brine (20 mL). The organic layer was dried with anhydrous MgSO_4_ and evaporated. The residue was purified by reverse flash column chromatography (from 5% to 70% ACN in H_2_O) to obtain **6** (350 mg, 0.66 mmol, 82% yield) as a sticky solid**. ^1^H NMR (400 MHz, CDCl_3_)** δ 8.43 (d, *J* = 9.7 Hz, 1H), 7.28 (t, *J* = 7.3 Hz, 2H), 7.23 – 7.16 (m, 3H), 5.64 (d, *J* = 7.5 Hz, 1H), 4.66 – 4.58 (m, 1H), 4.17 – 4.07 (m, 1H), 3.73 (s, 3H), 3.39 (dd, *J* = 10.0, 3.0 Hz, 1H), 3.12 (d, *J* = 9.8 Hz, 1H), 3.01 (dd, *J* = 10.8, 4.4 Hz, 1H), 2.97 – 2.89 (m, 2H), 2.86 – 2.71 (m, 4H), 2.64 (ddd, *J* = 11.9, 9.3, 7.0 Hz, 1H), 2.58 – 2.46 (m, 2H), 2.47 – 2.32 (m, 2H), 1.89 – 1.78 (m, 3H), 1.66 (dp, *J* = 13.0, 6.4 Hz, 1H), 1.44 – 1.35 (s + m, 10H), 0.89 (d, *J* = 6.6 Hz, 3H) 0.88 (d, *J* = 6.6 Hz, 3H). **^13^C NMR (101 MHz, CDCl_3_)** δ 176.1, 173.7, 155.5, 139.9, 128.7, 128.6, 126.3, 79.2, 66.8, 63.9, 59.8, 59.7, 56.3, 53.9, 52.5, 50.1, 47.9, 38.1, 37.9, 36.7, 35.3, 28.6, 26.2, 23.1, 22.7. **LC-MS (ESI)** *m/z* [M+H]^+^ 531.5.

#### (2S,4S)-4-[(2S,4S)-4-((tert-butoxycarbonyl)amino)-1-isopentylpyrrolidine-2-carboxamido]-1-phenethylpyrrolidine-2-carboxylic acid (Boc-NH-g-Amp(ipen)-Amp(phet)-COOH, 7)

An aqueous 0.1 M solution of LiOH (7.1 mL, 0.71 mmol) was added to a solution of **6** (252 mg, 0.47 mmol) in THF (20 mL). The mixture was stirred at rt for 3 h and a further 0.1 M LiOH was added (1.0 mL, 0.1 mmol). Then, the mixture was stirred at rt for 1.5 h and quenched with an aqueous HCl 1 M solution (800 µL). The mixture was concentrated under reduced pressure, dissolved in H_2_O (15 mL), and extracted with DCM:MeOH 9:1 (3 × 15 mL). The aqueous layer was basified with saturated NaHCO_3_ to pH = 9 and extracted with further DCM:MeOH 9:1 (3 × 15 mL). The combined organic layers combined were dried with anhydrous MgSO_4_ and concentrated under reduced pressure and lyophilized to obtain **7** (233 mg, 0.45 mmol, 95% yield) as a white solid. The product was used in the next step without further purification. **LC-MS (ESI)** *m/z* [M+H]^+^ 517.5.

#### (2S,4S)-4-[(2S,4S)-4-((tert-butoxycarbonyl)amino)-1-isopentylpyrrolidine-2-carboxamido]-1-phenethylpyrrolidine-2-carboxamide (Boc-NH-g-Amp(ipen)-Amp(phet)-CONH2, 8)

Et_3_N (100 µL, 0.96 mmol) was added to a mixture of **7** (164 mg, 0.32 mmol), NH_4_Cl (26 mg, 0.48 mmol), and HBTU (121 mg, 0.32 mmol) in ACN (5 mL). The mixture was stirred at rt for 1.5 h and evaporated. The residue was partitioned between DCM and a 5% NaHCO_3_ aqueous solution. The organic layer was dried with anhydrous MgSO_4_ and concentrated under reduced pressure. The residue was purified by reverse flash column chromatography (from 30% to 70% ACN in H_2_O) to obtain **8** (134 mg, 0.26 mmol, 82% yield) as a white solid. **^1^H NMR (400 MHz, CDCl_3_)** δ 7.50 (br s, 1H), 7.31 (t, *J* = 7.3 Hz, 2H), 7.25 – 7.20 (m, 3H), 6.52 (br s, 2H), 5.48 (br s, 1H), 4.44 (br s, 1H), 4.09 (br s, 1H), 3.18 (dd, *J* = 10.4, 3.7 Hz, 1H), 3.14 (d, *J* = 10.5 Hz, 1H), 3.08 (d, *J* = 9.7 Hz, 1H), 3.02 – 2.92 (m, 2H), 2.91 – 2.82 (m, 2H), 2.82 – 2.70 (m, 2H), 2.68 – 2.29 (m, 5H), 2.04 – 1.94 (m, 1H), 1.65 – 1.56 (m, 2H), 1.43 (s, 9H), 1.41 – 1.35 (m, 2H), 0.93 (d, *J* = 6.6 Hz, 3H), 0.92 (d, *J* = 6.6 Hz, 3H). **^13^C NMR (101 MHz, CDCl_3_)** δ 177.5, 173.4, 155.2, 139.9, 128.8, 128.7, 126.7, 80.0, 67.1, 66.3, 60.1, 58.9, 57.7, 54.0, 49.6, 49.0, 37.9, 37.2, 37.0, 35.4, 28.6, 26.4, 23.1, 22.7. **LC-MS (ESI)** *m/z* [M+H]^+^ 516.5.

#### (2*S*,4*S*)-4-[(2*S*,4*S*)-4-amino-1-isopentylpyrrolidine-2-carboxamido]-1-phenethylpyrrolidine-2-carboxamide, hydrochloride salt (HCl.NH_2_-γ-Amp(ipen)-Amp(phet)-CONH_2_, 9)

A solution of 4 M HCl in dioxane (4 mL, 16 mmol) was added to a solution of **8** (130 mg, 0.25 mmol) in dioxane (4 mL). The mixture was stirred at rt for 1 h and the volatiles were evaporated under reduced pressure. The title compound (**9)** was obtained quantitatively as a white solid and used in the next step without further purification. **LC-MS (ESI)** *m/z* [M+H]^+^ 416.5.

#### Boc-NH-γ-Amp(ipen)-Amp(phet)-Amp(ipen)-Amp(phet)-CONH_2_ (10)

DIPEA (48 µL, 0.28 mmol) was added to a mixture of **7** (142 mg, 0.28 mol), **9** (153 mg, 0.25 mol), HOBt·H_2_O (51 mg, 0.33 mmol) and EDC·HCl (73 mg, 0.38 mmol) in DMF (5 mL). The mixture was stirred at rt for 3 h and concentrated under reduced pressure. The residue was partitioned between EtOAc (20 mL) and saturated NaHCO_3_ (20 mL). The organic layer was washed with saturated NaCl (20 mL), dried with anhydrous MgSO_4_, and evaporated under reduced pressure. The title compound (**10**) was obtained as an orange solid (214 mg, 0.23 mmol 93% yield) and was used in the next step without further purification. **^1^H NMR (400 MHz, MeOD)** δ 7.30 – 7.24 (m, 4H), 7.22 – 7.14 (m, 6H), 4.48 – 4.41 (m, 1H), 4.41 – 4.35 (m, 1H), 4.31 – 4.24 (m, 1H), 4.00 (s, 1H), 3.17 (dd, *J* = 9.3, 5.3 Hz, 1H), 3.13 – 3.09 (m, 1H), 3.08 – 2.96 (m, 6H), 2.92 – 2.37 (m, 19H), 2.33 – 2.23 (m, 1H), 1.85 – 1.75 (m, 2H), 1.73 – 1.54 (m, 4H), 1.45 – 1.36 (s + m, 13H), 0.92 – 0.87 (m, 12H). **^13^C NMR (101 MHz, MeOD)** δ 179.3, 176.3, 175.8, 175.7, 157.7, 141.3, 129.7, 129.7, 129.5, 129.5, 127.3, 127.2, 80.1, 67.9, 67.6, 67.5, 67.3, 60.3, 60.1, 60.0, 58.0, 57.9, 55.0, 54.8, 51.4, 38.9, 38.8, 38.0, 37.9, 36.5, 36.2, 28.9, 27.5, 27.4, 23.4, 23.4, 23.1, 23.0. **LC-MS (ESI)** *m/z* [M+H]^+^ 914.5.

#### HCl.NH_2_-γ-Amp(ipen)-Amp(phet)-Amp(ipen)-Amp(phet)-CONH_2_(11)

A solution of 4 M HCl in dioxane (4 mL, 16 mmol) was added to a mixture of **10** (214 mg, 0.23 mmol) in dioxane (4 mL). The reaction mixture was stirred at rt for 1 h and the volatiles were evaporated under reduced pressure. The title compound (**11**) was obtained quantitatively as a white-off solid and used in the next step without further purification. **LC-MS (ESI)** *m/z* [M+H]^+^ 814.6.

#### Boc-γ-Amp(ipen)-Amp(phet)-Amp(ipen)-Amp(phet)-Amp(ipen)-Amp(phet)-CONH_2_ (12)

DIPEA (44 µL, 0.25 mmol) was added to a mixture of **7** (101 mg, 0.20 mol), **11** (187 mg, 0.23 mol), HOBt·H_2_O (46 mg, 0.30 mmol), and EDC·HCl (67 mg, 0.35 mmol) in DMF (5 mL). The mixture was stirred at rt for 3 h and concentrated under reduced pressure. The residue was partitioned between EtOAc (20 mL) and saturated NaHCO_3_ (20 mL). The organic layer was washed with saturated NaCl (20 mL), dried with anhydrous MgSO_4_, and evaporated under reduced pressure. The residue was purified by reverse flash column chromatography (from 5% to 50% ACN in H_2_O, with 0.1% HCOOH) to obtain **12** as a pale orange solid (202 mg, 0.15 mmol 79% yield). **^1^H NMR (400 MHz, MeOD)** δ 7.35 – 7.24 (m, 6H), 7.24 – 7.13 (m, 9H), 4.47 – 4.31 (m, 5H), 4.15 – 4.08 (m, 1H), 3.58 – 3.46 (m, 1H), 3.39 – 3.31 (m, 1H), 3.29 – 3.21 (m, 2H), 3.20 – 3.03 (m, 8H), 2.96 – 2.38 (m, 30H), 1.93 – 1.57 (m, 9H), 1.49 – 1.35 (s + m, 15H), 0.94 – 0.86 (m, 18H). **^13^C NMR (101 MHz, MeOD)** δ 178.9, 175.8, 175.3, 157.7, 141.1, 141.1, 141.0, 129.7, 129.7, 129.6, 129.5, 127.3, 127.3, 80.5, 68.2, 68.0, 67.7, 67.5, 60.3, 60.0, 58.0, 57.8, 57.8, 55.0, 54.7, 50.9, 38.4, 38.3, 38.0, 37.7, 36.1, 36.0, 36.0, 28.8, 27.5, 27.5, 27.3, 23.3, 23.3, 22.9, 22.9, 22.8. **LC-MS (ESI)** *m/z* [M+H]^+^ 1312.7. **HRMS (ESI)**: calculated exact mass for C_74_H_115_N_13_O_8_ [M+2H]^2+^ 656.9491, found 656.9496.

#### Boc-γ-Amp(ipen, me)-Amp(phet, me)-Amp(ipen, me)-Amp(phet, me)-Amp(ipen, me)-Amp(phet, me)-CONH_2_ (13)

MeI (360 µL, 5.7 mmol) was added to a mixture of **12** (25 mg, mmol) and NaHCO_3_ (19 mg, 0.23 mmol) in ACN (400 µL). The mixture was stirred at 35 °C for 7 days. Further MeI (360 µL, 5.7 mmol) was added and the mixture was stirred at 35°C for 7 days. The mixture was evaporated and it was suspended in MeI (720 µL, 11.4 mmol) and NaHCO_3_ (19 mg, 0.23 mmol), and then it was stirred at 35°C for 3 days. The final mixture was evaporated and purified by reverse flash column chromatography (from 5% to 40% ACN in H_2_O, with 0.1% HCOOH) to yield **13** (24.6 mg, 11 µmol, 58% yield) as an orange solid. **MALDI-TOF MS** 1398.5.

#### HCl.NH_2_-γ-Amp(ipen, me)-Amp(phet, me)-Amp(ipen, me-Amp(phet, me)-Amp(ipen, me)-Amp(phet, me)-CONH_2_(14)

A solution of 4 M HCl in dioxane (2 mL, 8 mmol) was added to a mixture of **13** (24.6 mg, 11 µmol) in dioxane (2 mL) and ACN (1 mL). The reaction mixture was stirred at rt for 3.5 h, and the volatiles were evaporated under reduced pressure. The product was used in the next step without further purification.

#### Ac-NH-γ-Amp(ipen, me)-Amp(phet, me)-Amp(ipen, me-Amp(phet, me)-Amp(ipen, me)-Amp(phet, me)-CONH_2_ (P33, 1)

DIPEA (13 µL, 75 µmol) was added to a solution of **14** (24 mg, 15 µmol) in Ac_2_O (0.5 mL) and ACN (0.5 mL). The mixture was stirred for 1 h at rt, quenched with water (1 mL), evaporated, and lyophilized. The residue was purified by semipreparative HPLC (from 5% to 60% ACN in H_2_O, with 0.1% TFA) to obtain **1** (13.4 mg, 6.6 µmol, 45% yield, 97% purity by HPLC) as a white solid. **HPLC** (from 5% to 100% ACN in H_2_O, with 0.05% TFA) rt 4.75 min. **LC-MS (ESI)** *m/z* [M]^2+^ 670.5, [M]^3+^ 447.6, [M]^4+^ 336.0. **MALDI-TOF MS** 1340.2.

### 2.3. Mouse strains and genotyping

In this study, several mouse strains were used: CD1 mice were purchased from Charles River, (Lyon, France), while APP/PS1 mice were purchased from Jackson Laboratory (Bar Harbor, ME, USA). APP/PS1 transgenic line (B6.Cg-Tg(APPswe, PSEN1dE9) expresses human APP carrying the *Swedish* mutation and also Presenilin 1 deleted in exon 9. Mice were genotyped with the method described previously by Jankowsky et al. [34] using three primers: PrP^C^ sense primer (5’-cctctttgtgactatgtggactgatgtcgg-3’), PrP^C^ antisense primer (5’-gtggataccccctcccccagcctagacc-3’), and one primer specific to the transgene (PS1: 5’-caggtggtggagcaagatg-3’, APP: 5’-ccgagatctctgaagtgaagatggatg-3’). All studies were performed under the guidelines and protocols of the Ethical Committee for Animal Experimentation (CEEA) at the University of Barcelona, and the protocol for the use of animals in this study was reviewed and approved by the CEEA at the University of Barcelona (CEEA approval# 207/24).

### 2.4. Primary cortical cultures

Primary cultures from E16.5 CD1 embryos were prepared in ice-cold PBS pH 7.3 supplemented with 6.5 mg/mL glucose, as previously described [35]. Briefly, mouse embryo brains were dissected and washed in ice-cold PBS containing 6.5 mg/mL glucose. Meninges were removed, and the cortical lobes isolated. Tissue was trypsinized, and cells were dissociated by trituration in PBS containing 0.025% DNase with a polished pipette. Dissociated cells were plated at ≈3,000 cells/mm^2^ on plates coated with poly-D-lysine. The culture medium used was Neurobasal^TM^ supplemented with 2 mM glutamine, 6.5 mg/mL glucose, antibiotics (Pen./Strept.), 5% horse serum, and 1x B27 (Invitrogen-Life Technologies). After 24 h in culture, the serum was removed to reduce glial proliferation, and the media was changed every 3 days.

### 2.5. *In vitro* γ-peptides treatment

After 6 DIV, neural primary cultures were treated with γ-peptides at 25 µM for 48 or 120 h (8 and 11 DIV, respectively). During this period, the media remained unchanged, and it was collected with 1x protease inhibitor and analysed by ELISA. In addition, cells were collected for western blot analysis with lysis buffer: 50 mM Hepes, 150 mM NaCl, 1.5 mM MgCl_2_, 1 mM EGTA, 10% glycerol, 1% Triton X-100 with supplemental 1x protease inhibitor cocktail (Roche Diagnostic) and 0.1 M sodium fluoride, 10 mM sodium pyrophosphate, and 200 μM sodium orthovanadate as phosphatase inhibitors. After this, samples were centrifuged at 13,000 x *g* for 20 min at 4°C. The resulting supernatant was normalized for protein content using the bicinchoninic acid (BCA) kit (Pierce). Alternatively, samples were collected according to the protocol described below to quantify the BACE1 and BACE2 activities of the cell extracts.

### 2.6. Human BBB cellular model assay

*In vitro* model of BBB was made using human endothelial cells derived from pluripotent stem cells co-cultured with bovine pericytes [36]. Briefly, the monolayer of human endothelial cells was co-cultured with bovine pericytes for 8 days. Lucifer Yellow is used as an internal standard (of monolayer quality). Assay is performed at 50 µM for 2h at 37°C. Both donor and acceptor are analyzed by HPLC or UPLC. Apparent permeability (Papp*10^6^) was calculated as follows: Papp=(dQ/dt)*(1/A)*(1/C_0_) (cms^-1^) (see [37] and [38] for technical details).

### 2.7. P33 intraperitoneal injection of APP/PS1 mice

We developed two treatment strategies: acute and chronic. For the acute assay, 6-month-old APP/PS1 mice (13 mice: 6 females and 7 males) were treated intraperitoneally with PBS (6 mice: 3 females and 3 males) or 200 µg/kg/day of P33 (7 mice: 3 females and 4 males) for 3 consecutive days. Three days after the last administration, animals were killed and their brains were removed. The CTX and HIP of each animal were dissected, divided into two hemispheres, and immediately frozen at -80°C. Subsequently, one half of each brain was processed for biochemical analysis and the other half for RNA extraction.

For the chronic assay, 12-month-old APP/PS1 mice (19 animals: 11 females and 8 males) were treated intraperitoneally with PBS (11 animals: 7 females and 4 males) or 200 µg/kg/day of P33 (8 animals: 4 females and 4 males) 3 times per week for 8 consecutive weeks. Three days after the last administration, the animals were subjected to behavioural tests and later killed to collect one half of the CTX and HIP for further biochemical analysis (immediately frozen at -80°C) while the other half was fixed in 4% PFA for immunohistochemical assay. In addition, several tissues from each animal were removed for histopathological analysis. Specifically, liver, lung, kidney, and spleen were immersion-fixed in 4% PFA, embedded in parafilm, and sectioned for haematoxylin and eosin staining.

In both cases, samples collected for biochemical assays were homogenized in lysis buffer: 100 mM Tris pH 7.0, 100 mM NaCl, 10 mM EDTA, 0.5% NP-40, and 0.5% sodium deoxycolate with supplemental 1x protease inhibitor cocktail and with supplemental 1x protease inhibitor cocktail (Roche Diagnostic) and 0.1 M sodium fluoride, 10 mM sodium pyrophosphate, and 200 μM sodium orthovanadate as phosphatase inhibitors. After this, samples were centrifuged at 13,000 x g for 20 min at 4°C. The resulting supernatant was normalized for protein content using the BCA kit (Pierce).

### 2.8. ELISA A**β**1-42 quantification

Brain homogenates from APP/PS1 mice and culture media collected from treated neural cultures were analysed by ELISA to reveal the levels of Aβ. Brain homogenates obtained as previously described were diluted 1:2 in standard diluent buffer and 1x protease inhibitor. Media from the previously described cultures were also analysed without diluting the sample. Aβ levels were measured using the Aβ1-42 Human ELISA kit (Invitrogen Life Technologies) following the manufacturer’s protocol. Microwell plate absorbance at 450 nm was read with an Opsys MR microplate reader (Dynex Technologies).

### 2.9. BACE1 and BACE2 activity

β-secretase-1 activity was quantified using the fluorometric kit BACE1 Activity detection kit (CS0010). Briefly, neuronal primary culture cells treated with the γ-peptides were scraped and collected after 48 or 120 h of treatment with CelLytic M (Sigma). Samples were centrifuged at 13,000 x *g* for 20 min at 4°C, and the resulting supernatant was used to assay enzyme activity following the manufacturer’s protocol with excitation and emission wavelengths of 320 and 450 nm, respectively. For β-secretase-2 activity quantification the fluorometric kits used was Sensolyte^®^ 520 BACE2 Activity assay kit (AS-72225). In this case, neuronal primary culture cells were also scraped and collected after 48 or 120 h of treatment using the assay buffer supplied with the kit. Samples were centrifuged at 2,500 x *g* for 10 min at 4°C, and the resulting supernatant was used to assay enzyme activity following the manufacturer’s protocol with excitation and emission wavelengths of 490 and 520 nm, respectively. In both cases, fluorescence was measured after the substrate was added to determine the “time zero” point and after 2 h of incubation, using an Infinite M200 Pro Microplate Reader (Tecan Trading AG).

### 2.10. Western blot analysis

Soluble homogenate from the cortical brains of APP/PS1 mice and from cultured cells was processed for western blotting. Samples obtained as previously indicated were boiled in Laemmli sample buffer at 96°C for 5 min, followed by 10% SDS-PAGE. They were then electrotransferred into nitrocellulose membranes for 1 hour at 4°C, 100V. Membranes were blocked with 5% non-fat milk in 0.1 M Tris-buffered saline (pH 7.4) for 1 hour and incubated overnight in 0.5% blocking solution containing primary antibodies. Polyclonal antibody against caspase-3 (cleaved Asp175) was used to determine cleaved caspase-3 levels, and monoclonal anti-α-tubulin was used for standardization. After incubation with peroxidase-tagged secondary antibodies (1:2000 diluted, VectorLabs), membranes were revealed with the ECL-plus chemiluminescence Western blot kit (Amersham-Pharmacia Biotech). For western blot quantification, developed films were scanned at 2400 × 2400 dpi (i800 MICROTEK high quality film scanner (Microtek International), and the densitometric analysis was performed using Fiji™ software.

### 2.11. Real-time quantitative PCR

Total RNA from cortex samples obtained from treated mice was extracted with mirVana isolation kit (Ambion). Purified RNAs were used to generate the corresponding cDNAs required as templates for the RT-qPCR amplification. The primers used in this study were: 5’-agcaaaccaccaagtggagga-3’ and 5’-gctggcaccactagttggttgt-3’ for mouse TNFα; 5’-ttgtggctgtggagaagctgt-3’ and 5’-aacgtcacacaccagcaggtt-3’ for mouse IL1β [39]. PCR reaction was performed with Roche LightCycler 480 detector, using 2x SYBR Green master Mix (Roche) as reagent, following the manufacturer’s protocol. Reaction consisted of a denaturation-activation cycle (95°C for 10 min) followed by 40 cycles of denaturation-annealing-extension (95°C for 10 seconds, 55°C for 15 seconds and 72°C for 20 seconds). mRNA levels were quantified using LightCycler 480 software. Data were analysed with SDS 1.9.1 Software (Applied Biosystem method of Applied [40]. Results were normalized by the expression levels of the housekeeping gene, *GAPDH* (5’-aggtcggtgtgaacggatttg-3’) and (5’-tgtagaccatgtagttgaggtca-3’), which were quantified simultaneously with the target gene [41].

### 2.12. Cytokine profiler array

Cytokine antibody array was performed with the mouse cytokine array panel A (ARY006, RD Systems). Briefly, brain homogenates from 2 APP/PS1 mice treated with P33 and 2 treated with PBS were analysed. 200 μg of tissue lysate obtained as previously indicated was incubated with each array membrane according to the manufacturer’s protocol. Signals were analysed using the streptavidin-horseradish peroxidase supplied by the kit, followed by chemiluminescent detection with Image Quant LAS 4000. The densitometric analysis in the captured images was performed using Fiji™ software.

### 2.13. Histological studies

Brains fixed in 4% buffered PFA were sectioned (30 μm) on a freezing microtome (Leica). They were then pretreated with 70% formic acid for 30 min before being processed for the immunohistochemical detection of Aβ, following an immunoperoxidase protocol. Briefly, sections were rinsed in PBS, and endogenous peroxidase activity was blocked by incubation in 3% hydrogen peroxide (H_2_O_2_) and 10% methanol dissolved in PBS. After extensive rinsing, sections were blocked in PBS containing 0.2% gelatin, 10% normal goat serum, 0.2% glycine, and 0.2% Triton X-100, for 1 h at room temperature. Afterwards, sections were incubated for 16 h at 4°C with the anti-Aβ antibody 4G8 (1:500; Covance, Princeton, NJ, USA). After that, sections were incubated with anti-mouse biotinylated secondary antibody (2 h, 1:200 diluted) and streptavidin-horseradish peroxidase complex (2 h, 1:400 diluted). Peroxidase activity was revealed with 0.025 % diaminobenzidine (DAB) and 0.003 % H_2_O_2_. After rinsing, sections were mounted onto slides and dehydrated, and coverslipped with Eukitt^TM^ (Bio-Optica Milano S.p.A, Milan, Italy). Liver, lung, kidney, and spleen fixed in 4% PFA were processed for embedding in paraffin. Lastly, they were sectioned in a microtome (Leica) at 4 µm thickness and adhered to a gelatinized glass slide. Sections were stored at 37°C until hematoxylin-eosin (HE) staining, and then deparaffinized in xylene (2 changes, 15 min each), rehydrated in a graded ethanol series (100%, 95%, and 70%, 10 min each), and briefly washed in distilled water. After HE staining, sections were dehydrated in ethanol, cleared in xylene, and mounted in Eukitt^TM^. Images were obtained using an Olympus BX61 microscope equipped with a cooled DP72L digital camera.

### 2.14. Novel object recognition test

The novel object recognition (NOR) test was conducted using a square white open-field arena (50 × 50 × 40 cm), with 10 mice (5 treated with PBS and 5 treated with P33). The experiment consisted of two phases, each lasting 20 min. In the first phase (object recognition (OR)), two identical objects were placed diagonally in the box. In the second phase (NOR), one object was replaced with another different one in shape and color. The arena and objects were cleaned and disinfected with 70% ethanol between trials. Mouse behavior was recorded and analyzed using MouseBeat software [42], with exploration defined as head approaches or touches to the objects.

### 2.15. Statistical analysis

Data analysis was performed using Prism 8.0 (GraphPad Software). The differences between groups were analysed using Student’s *t*-test throughout the study. For the novel object recognition test, statistical analysis was performed using a two-way ANOVA. All data are expressed as mean ± standard error of the mean (SEM). Significant differences were noted as *p* < 0.05 (*), *p* < 0.01 (**), *p* < 0.001 (***), and *p* < 0.0001 (****).

## 3. Results

### 3.1. Identification of γ-peptides that reduce amyloid-β release in primary neuronal cultures

An initial cytotoxicity screening was performed in cultured cerebellar granule cells (CGCs) [43] using a focused library of 50 cis-γ-amino-L-proline-derived peptides (library preparation and cell penetrating screening to be reported elsewhere) at concentrations of 12.5, 25, and 50 µM. Eighteen γ-peptides displaying low cytotoxicity were selected from this screening for further evaluation (for the chemical structures of these γ-peptides see Supplementary Figure 1). The safety profile of these peptides was subsequently confirmed in primary cortical neuronal cultures, in which no significant toxicity was observed at 25 µM (see Supplementary Figure 2).

To evaluate the capacity of the selected γ-peptides to modulate endogenous amyloidogenic processing [44], primary cortical neurons derived from CD1 mouse embryos were cultured at three different plating densities. The temporal profile of Aβ release into the culture medium was monitored at 8, 11, and 15 days *in vitro* (DIV). Quantification of secreted Aβ1–42 by ELISA revealed reproducible peaks of Aβ release at 8 and 11 DIV. These time points were used for peptide treatment experiments (data not shown).

Neuronal cultures were treated at 6 DIV with each γ-peptide (25 µM), and conditioned media were collected after 48 or 120 h to quantify Aβ1–42 levels. Compared to vehicle-treated cultures, most γ-peptides reduced Aβ release at 11 DIV (Figure 1A). Among them, peptides P24, P33, and P39 produced the most pronounced decreases, reducing Aβ levels to 0.34 ± 0.12, 0.38 ± 0.09, and 0.45 ± 0.02 relative units, respectively (mean ± SEM). Notably, P33 and P39 also significantly reduced Aβ release at 8 DIV, while P24 showed a stronger effect at the later time point.

**Figure 1.**
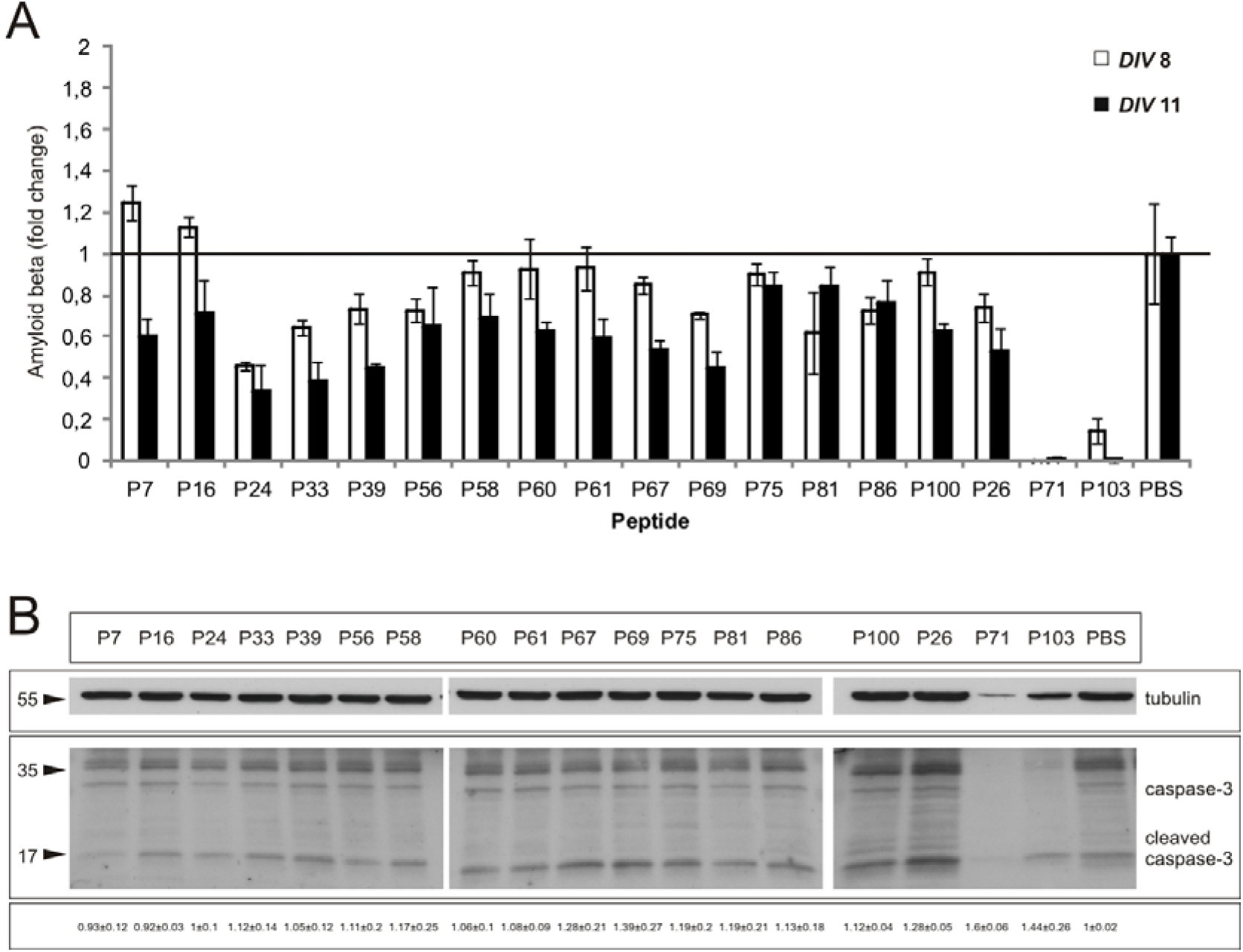
Phenotypic screening of γ-peptides reveals differential modulation of Aβ production and neuronal viability. Primary neuronal cultures derived from CD1 mouse embryonic cortices were pleated and treated with the previously selected γ-peptides (25 µM) or vehicle (PBS). Treatments were initiated at 6 days *in vitro* (DIV), and conditioned media were collected after 48 h (8 DIV) or 120 h (11 DIV). (A) Levels of secreted Aβ1–42 in the culture medium were quantified by ELISA and compared to vehicle-treated controls. Data are presented as mean ± SEM. (B) Western blot analysis of total caspase-3 and cleaved caspase-3 levels in cell lysates was performed to assess potential peptide-induced cytotoxicity. Tubulin was used as a loading control and normalization.

To exclude the possibility that the reduction in Aβ levels was caused by peptide-induced neuronal loss, cell lysates were collected at 8 DIV and analysed for cleaved caspase-3 by Western blot. Several γ-peptides, including P58, P67, and P69, reduced Aβ release by 30%–50%. However, they also induced a marked increase in cleaved caspase-3 levels (Figure 1B), indicating peptide-associated cytotoxicity. These peptides were excluded from further analysis. In contrast, peptides such as P24, P33, and P39 did not significantly activate caspase-3, supporting a noncytotoxic mechanism underlying their effects on Aβ release. Other γ-peptides, such as P7 and P16, showed toxicity levels 10% lower than the vehicle.

### 3.2. Phenotypic structure–activity relationships of **γ**-peptides affecting A**β** production

A comparative analysis of peptide structures revealed shared molecular features among the most active γ-peptides (P24, P33 and P39; Figure 2). These included (i) a high overall positive charge distributed along the sequence, arising either from bis-alkylated aminoproline residues forming quaternary ammonium groups (P33 and P39) or from primary amines on alternating side chains (P24), and (ii) an alternating pattern of residues bearing aromatic (phenethyl or indolylacetyl) and aliphatic moieties such as isopentyl or aminopropyl moieties (Figure 2). In contrast, two peptides, P7 and P16, failed to reduce Aβ release at 8 DIV, producing a modest increase in Aβ levels (approximately 15–20%) relative to the control (Figure 1A). Despite sharing similar amine density and residue patterning with P33, peptides P7 and P16 differ in the nature of their cationic functionalities. P33 has six positive charges that are quaternary ammonium groups derived from bis-alkylated aminoproline residues. In contrast, P7 and P16 contain tertiary ammonium groups resulting from monoalkylation. Additionally, P7 has acetylated residues at both the N-and C-termini, and P16 lacks the isopentyl substituents present in P33. These observations suggest that both the chemical nature of the cationic groups and the spatial presentation of the diverse side chains both contribute to the Aβ-lowering activity observed in this peptide series.

**Figure 2.**
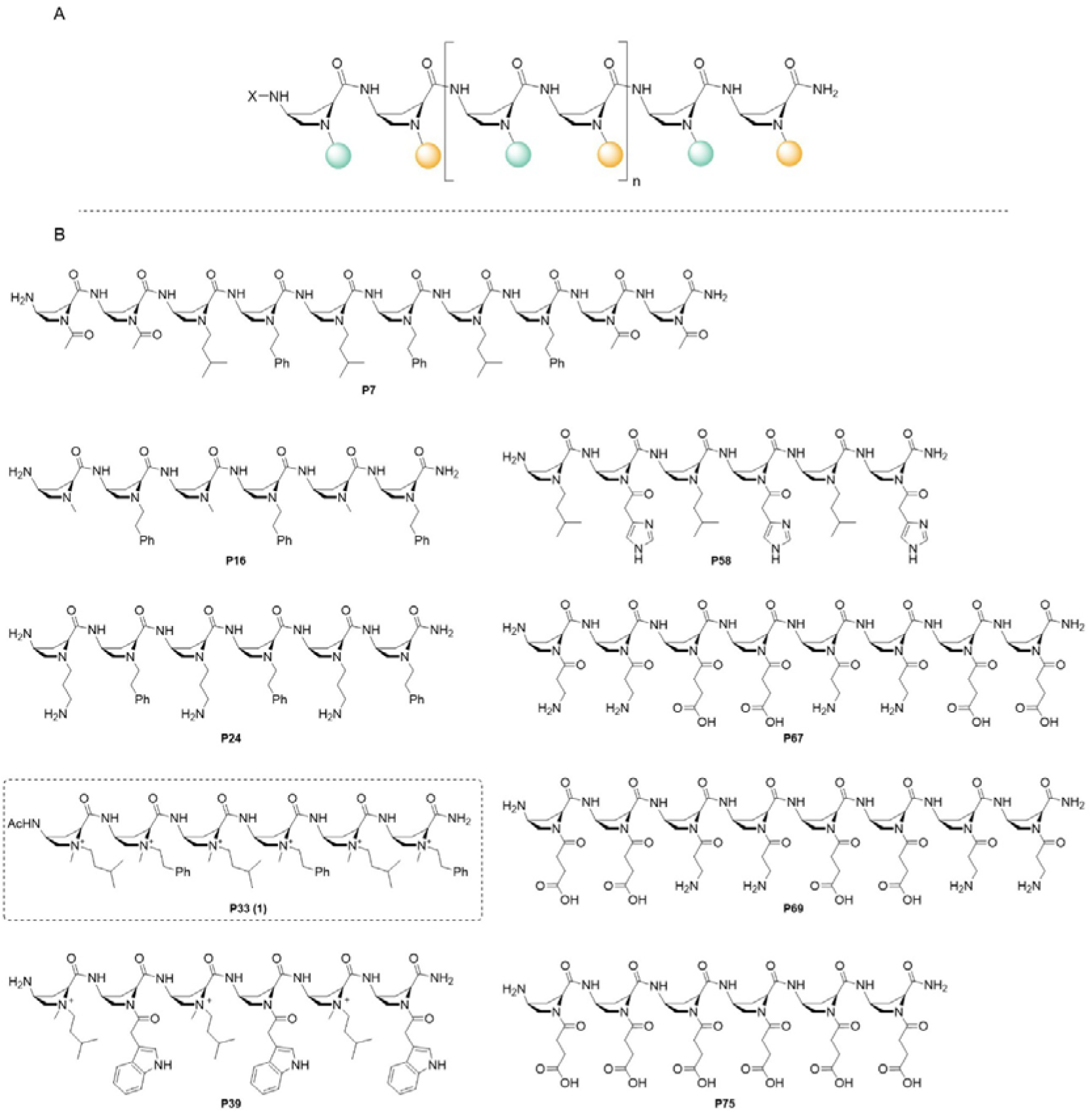
(A) General structure of the cis-γ-amino-L-proline peptides. (B) Structures of the γ-peptides that showed differential modulation of Aβ production.

The structure of these γ-peptides, which exhibited a 30–50% reduction in Aβ release and a significant increase in cytotoxicity (P58, P67, and P69), differs from that of the most active peptides in terms of (i) length and different patterns of aminopropionyl- and succinyl-derivatized aminoproline residues (P67 and P69); and (ii) the substitution of the aromatic moieties by imidazolylacetyl and the presence of three tertiary amines (P58).

Taken together, these results indicate that cis-γ-amino-L-proline-derived peptides can modulate endogenous Aβ production in primary neurons in a sequence-dependent manner. The data also suggest that γ-peptides with a high positive charge density, particularly from quaternary ammonium functionalities, and an alternating presentation of aromatic and aliphatic side chains are more likely to reduce Aβ release while preserving neuronal viability.

### 3.3. Modulation of **β**-secretase activity by selected **γ**-peptides in neuronal cultures

To further characterize the cellular effects of the most active and non-cytotoxic γ-peptides, four peptides (P7, P24, P33, and P39) were selected for analysis of β-secretase-related activity in primary neuronal cultures at 8 and 11 DIV. This selection was based on their ability to modulate Aβ release and their favorable toxicity profiles.

Cell lysates were analyzed using a fluorometric assay that detects β-secretase enzymatic activity within the cells. Among the tested peptides, P33 induced a robust and reproducible reduction in β-secretase activity, exceeding 50% inhibition at both analyzed time points (Figure 3A). In contrast, P7 did not significantly alter β-secretase activity, despite reducing Aβ levels at 11 DIV. P39 failed to reduce enzymatic activity at either time point, despite its pronounced effect on Aβ release. Peptide P24 produced a modest reduction in β-secretase activity (∼25%) only at 11 DIV.

**Figure 3.**
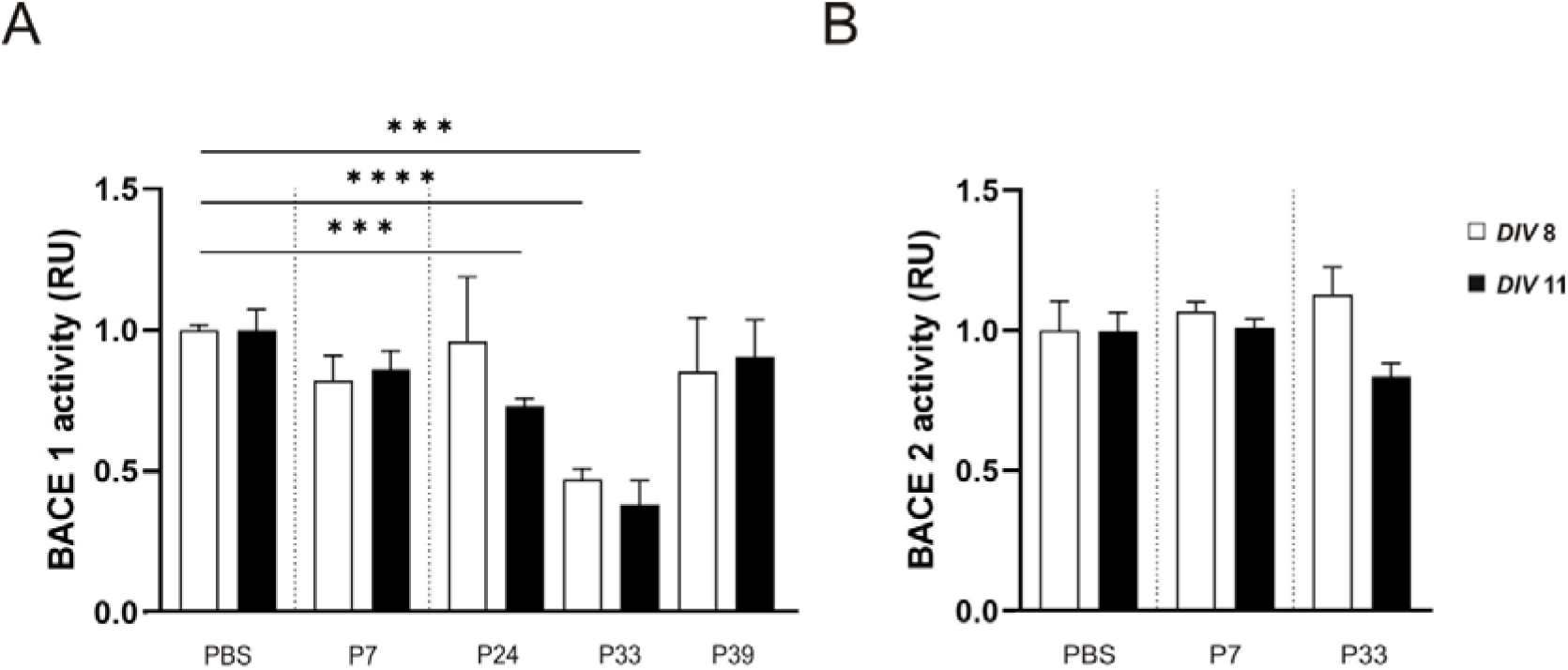
Cell-based assessment of β-secretase–associated activity following treatment with selected γ-peptides. Primary cortical cultures from CD1 mouse embryos were treated with selected γ-peptides (25 µM) or vehicle (PBS). Cells were harvested at 8 and 11 days *in vitro* (DIV), and β-secretase–associated activity was quantified using a fluorogenic substrate assay. (A) BACE1-associated activity and (B) BACE2-associated activity are shown relative to vehicle-treated controls (normalized to 1). Results are shown as the fold change (relative units, RU). Values from the vehicle group were used as the reference (normalized to 1), Data are presented as mean ± SEM. Asterisks indicate statistically significant differences (*** p < 0.001, **** p < 0.0001, Student t-test).

Given these differential effects, we further explored the specificity of P33 by assessing its impact on BACE2 activity under the same experimental conditions. Unlike its effect on β-secretase activity, P33 did not significantly alter BACE2 activity (Figure 3B). The structurally related control peptide, P7, also failed to affect BACE2 activity. Although these assays do not demonstrate direct enzyme inhibition using purified BACE1, the consistent reduction of BACE1-associated activity in neuronal cultures, coupled with the absence of comparable effects on BACE2, suggests P33’s functional selectivity within a cellular context.

Because of its consistent reduction of Aβ release, favorable cytotoxicity profile, and selective modulation of β-secretase-associated activity in neuronal cultures, P33 was selected for subsequent *in vivo* evaluation using an APP/PS1 mouse model of amyloid pathology.

### 3.4. *In vitro* BBB permeability of the γ-peptide P33

Given the central nervous system–restricted pathology of Alzheimer’s disease, we examined whether the γ-peptide **P33** exhibits the ability to traverse an *in vitro* model of the human blood–brain barrier (BBB). BBB transport was assessed using a static Transwell-based system comprising induced pluripotent stem cell (iPSC)-derived brain endothelial cells co-cultured with human pericytes [36,45,46]. Apparent permeability (Papp) values were determined as a comparative measure of transendothelial transport.

**P33** exhibited measurable BBB permeability, with a Papp value of 5.48 ± 0.59 × 10LJLJ cm/s (Table 1). This value falls within the range reported for several previously described cell-penetrating peptides and BBB shuttle peptides evaluated using comparable *in vitro* systems [38,47]. In contrast, the γ-peptide P75, which was included as a negative control due to its limited cell-penetrating properties, showed markedly reduced permeability (Papp = 0.83 ± 0.16 × 10LJLJ cm/s).

**Table 1.** Apparent permeability (Papp) of P33 and the γ-peptide negative control (P75) across the *in vitro* BBB model. Data are expressed as mean ± SEM.

| Peptide | Papp*10<sup>6</sup> |
| --- | --- |
| P33 | 5.48±0.59 |
| P75 | 0.83±0.16 |

### 3.5. Large scale synthesis of P33

At this point, larger quantities of **P33** (**1**) were needed for *in vivo* experiments. **P33** was initially synthesized alongside the full γ-peptide library using solid-phase peptide synthesis (SPPS) with anFmoc/Boc/Alloc protection strategy on a p-methylbenzhydrylamine (MBHA) resin. In this synthesis, the Fmoc group was chosen as the temporary protecting group for the γ-amino group of the 4-aminoproline (Amp) residues used to build the γ-peptide backbone. Boc and Alloc were used as semi-permanent protecting groups for the α-amino position of each Amp residue, following the substitution pattern of P33. Removal of the Boc or Alloc groups allowed introduction of the side chains via alkylation with the corresponding aldehyde derivatives. Finally, the six residues were simultaneously methylated. However, this strategy required the use of hydrofluoric acid (HF) for cleavage. This procedure became obsolete due to the hazards associated with handling HF, particularly on a large scale. Therefore, we explored resins such as Rink Amide polystyrene or ChemMatrix. These resins allow the synthesis of **P33** under mild acidolysis conditions using TFA cocktails. In this synthesis, preformed 4-aminoproline monomers that already bear the mimetic natural amino acid side chain at the α position were used. This procedure was challenging due to the insolubility of the residues, which resulted in dirty crudes and very low yields in all attempts.

Given the repeated dipeptide motif in the **P33** structure, a modular solution-phase strategy was developed. This strategy involved synthesizing a dipeptide intermediate to produce **P33**, thereby reducing the number of required coupling reactions. In the initial attempt, dialkylated monomers were prepared via reductive amination at the α-position of (2S,4S)-methyl-(tert-butoxycarbonyl)-4-((tert-butoxycarbonyl)amino)pyrrolidine-2-carboxylate, followed by tertiary amine methylation. However, the high acidity of the α-proton, due to the positively charged amino group, led to epimerization issues that could not be overcome (see Supplementary Scheme 1). To avoid this issue, the methylation step was postponed until after peptide elongation. Then, after convenient deprotection, the monoalkylated residues (**2** and **4**) were coupled using EDC/HOBt after convenient deprotection to obtain dipeptide 7 in a 78% yield (see Scheme 1).

#### Scheme 1. Synthesis of the repeated dipeptide (7)^a^

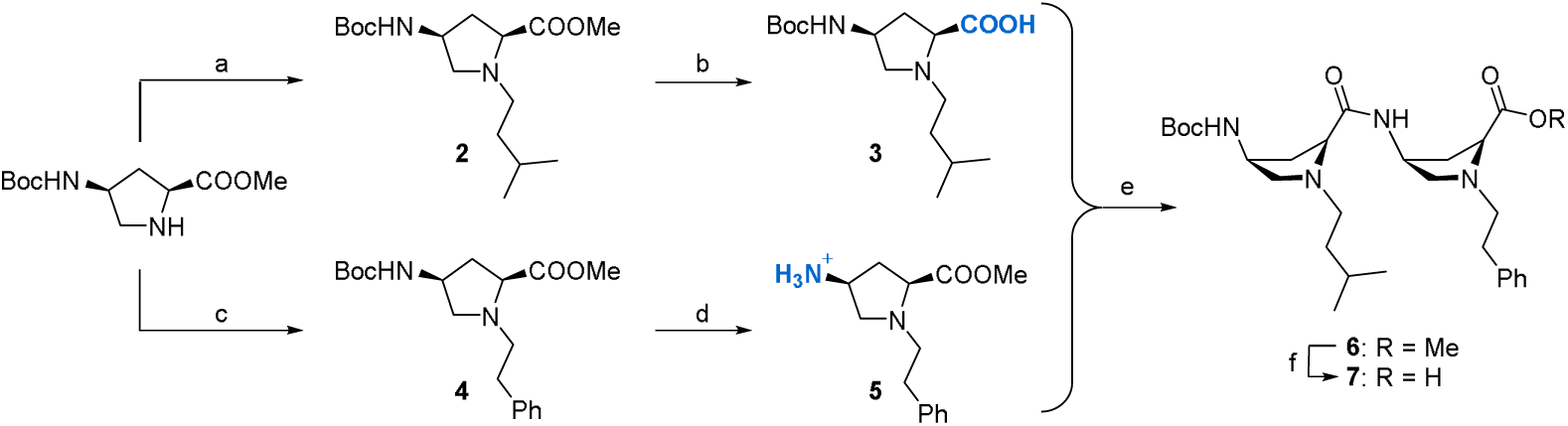

^a^Reagents and conditions: (a) isovaleraldehyde, NaBH_3_CN, DCE, rt, 16 h, 54%; (b) LiOH, THF, H_2_O, rt, 2 h 93%; (c) phenylacetaldehyde, NaBH_3_CN, DCE, rt, 16 h, 63%; (d) HCl, dioxane, rt, 1.5 h, 99%; (e) EDC·HCl, HOBt·H_2_O, DIPEA, DMF, rt, 16 h; (f) LiOH, THF, rt, 3 h, 78% (two steps).

Dipeptide **7** was converted into its carboxamide with an 82% yield using NHLJCl and HBTU to produce the C-terminal dipeptide **8** (see Scheme 2, step a) [48]. With **7** and **8** on hand, synthetic strategy A (described in Supplementary Scheme 2) was attempted. In this strategy, the intermediates were methylated after each dipeptide incorporation. However, the three methylation steps were unreasonably long, and the process was incomplete. Purification was required after each methylation step, resulting in an overall yield of 7%. To optimize the route (strategy B, Scheme 2), the methylation of the residues was postponed until the entire γ-backbone had been constructed. This time, amide dipeptide **8** was deprotected and coupled twice with dipeptide **7** using EDC/HOBt (Scheme 2, steps b–e). This strategy produced the monoalkylated backbone **12** via simple aqueous workups without further purification in a 200 mg scale with an overall yield of 60%. Finally, permethylation and deprotection/acetylation of the N-terminal group produced **P33** (**1**) with a 26% yield (Scheme 2, steps f–h).

#### Scheme 2. Synthesis of P33 from dipeptide 7, Strategy B^a^

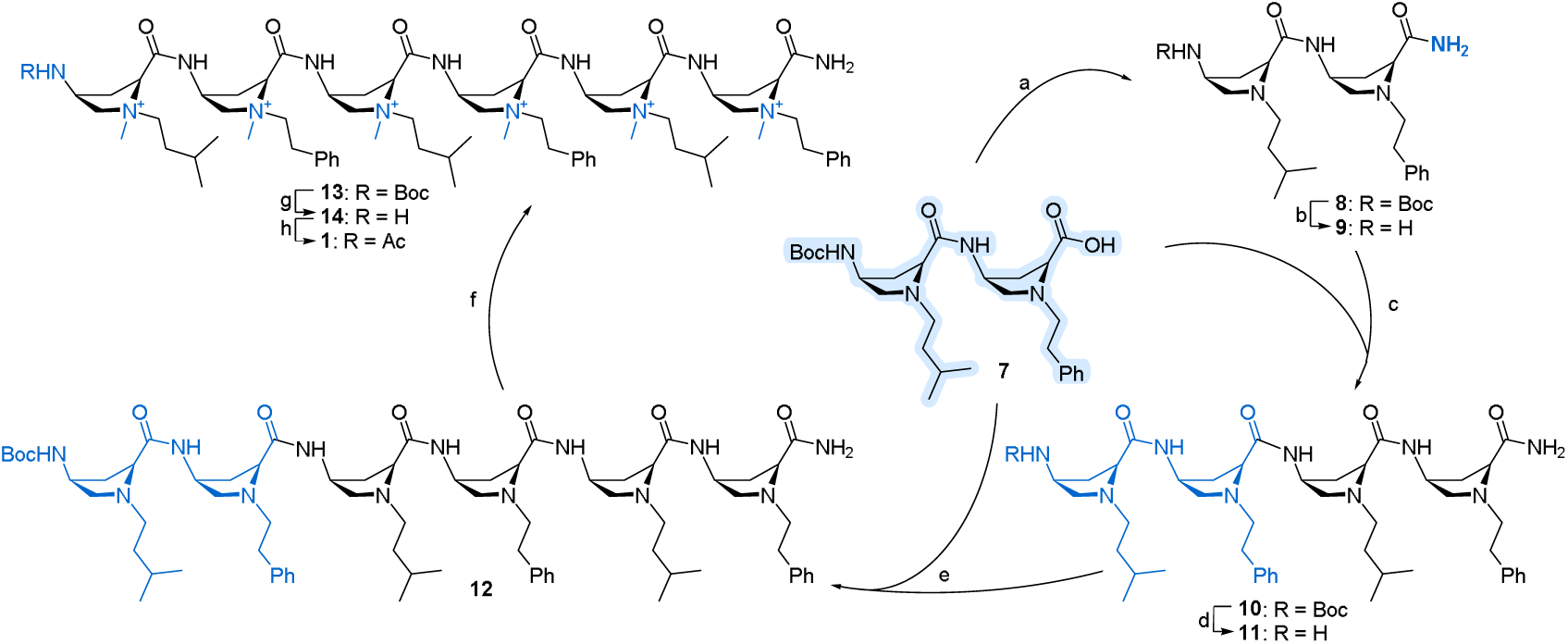

^a^Reagents and conditions: (a) NH_4_Cl, HBTU, Et_3_N, ACN, rt, 1.5 h, 82%; (b) HCl, dioxane, ACN, rt, 1 h; (c) 7, EDC·HCl, HOBt·H_2_O, DIPEA, DMF, rt, 3 h, 93% (two steps); (d) HCl, dioxane, ACN, rt, 1 h; (e) 7, EDC·HCl, HOBt·H_2_O, DIPEA, DMF, rt, 3 h, 79% (two steps); (f) MeI, NaHCO_3_, ACN, 35 LJC, 14 d, 58%; (g) HCl, dioxane, ACN, rt, 3.5 h; (h); Ac_2_O, DIPEA, ACN, rt, 1 h, 45% (two steps).

### 3.6. Acute intraperitoneal injection of P33 significantly reduced A**β**1-42 levels in the brains of APP/PS1 mice

An acute treatment study was conducted on APP/PS1 mice to evaluate the *in vivo* efficacy of P33. Seven six-month-old APP/PS1 mice received daily intraperitoneal injections of P33 for three consecutive days. Six littermates were treated with PBS as vehicle control. Six days after the first injection, the brains were collected and processed. Homogenized brain tissues were analysed using ELISA, Western blot, and RT-qPCR (Figure 4).

**Figure 4.**
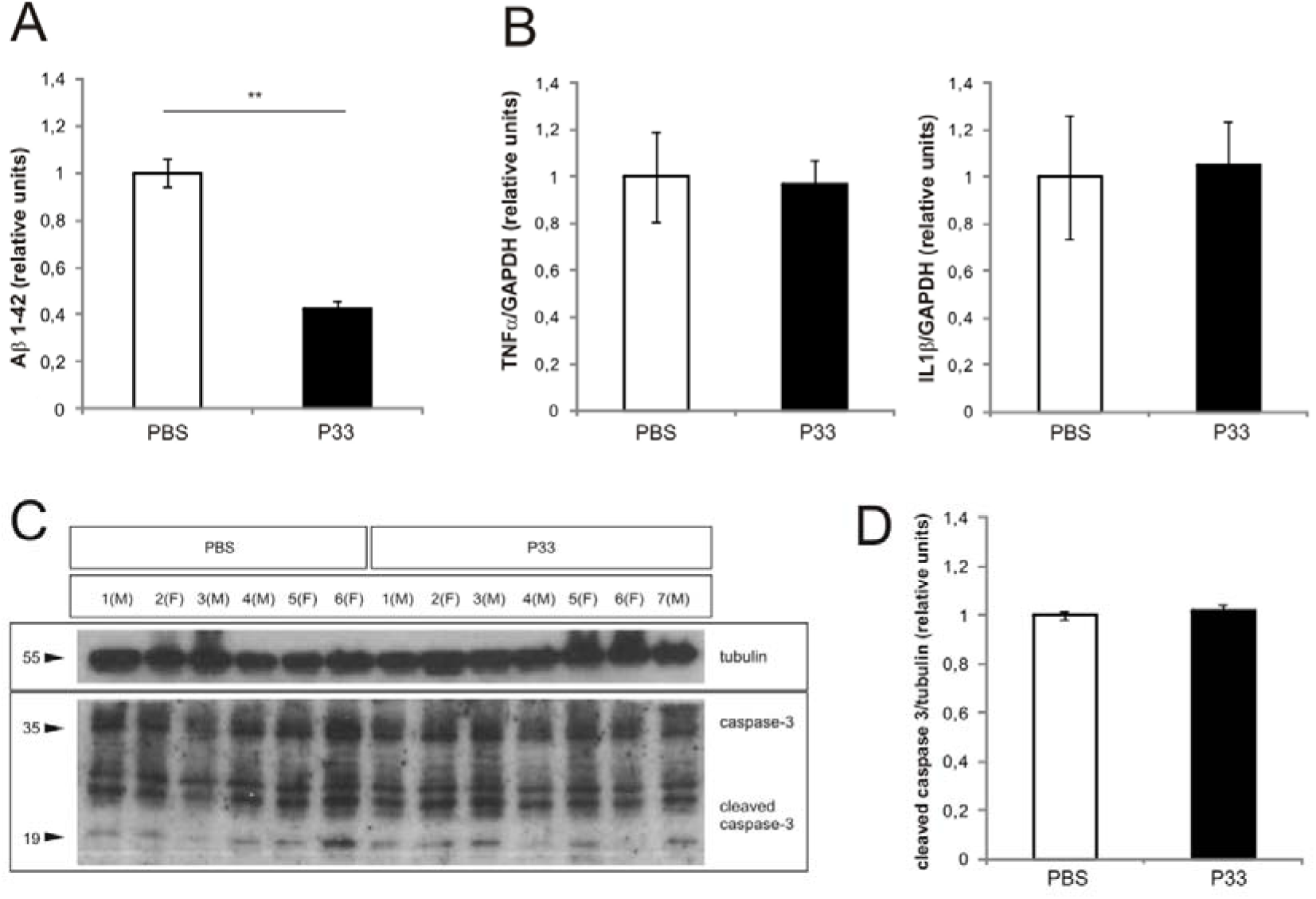
Acute *in vivo* acute treatment with P33 in APP/PS1 mice. Seven 6-month-old APP/PS1 mice received intraperitoneal injections of P33, while six APP/PS1 littermates were treated with PBS as vehicle control for three consecutive days. Brains were collected and processed 6 days after the first injection. (A) Aβ1-42 levels were quantified by ELISA in soluble brain extracts. (B) Inflammatory response determined (C) Western blots of caspase-3 and cleaved caspase-3 in soluble brain extracts. (D) Densitometric analysis of cleaved caspase-3 levels. Data are presented as mean ± SEM. Asterisks indicate statistically significant differences (** *p* < 0.01, Student *t*-test).

Quantification of Aβ burden by ELISA revealed a significant reduction in Aβ1–42 levels in the brains of P33-treated mice compared to vehicle-treated controls (−45.53%, p = 0.019; see Figure 4A). These results suggest that acute P33 administration effectively reduces cerebral Aβ1–42 levels in this Alzheimer’s disease mouse model.

To evaluate the safety of acute P33 treatment, we assessed the potential induction of neuroinflammation by measuring proinflammatory cytokine expression after administration. RT-qPCR analysis of brain mRNA showed no significant changes in tumor necrosis factor alpha (TNF-α) or interleukin-1 beta (IL-1β) expression in P33-treated animals compared to controls (Figure 4B).

Alzheimer’s disease (AD) is characterized by impairment in the cholinergic and glutamatergic systems, which are involved in processes such as memory and learning[49]. Dysfunction in the glutamatergic system can lead to increased neuronal loss through an apoptotic process, which reduces Aβ levels in the brain. Since reductions in Aβ levels could theoretically result from increased neuronal loss, we examined whether P33 treatment induces apoptosis. However, Western blot analysis of cleaved caspase-3 levels showed no significant differences between P33-treated and control animals (Figure 4C–D). These results suggest that the observed reduction in Aβ1–42 was not associated with increased cell death.

### 3.7. Chronic intraperitoneal administration of P33 reduces amyloid plaque burden and improves novel object recognition in aged APP/PS1 mice

After characterizing acute P33 administration and its safety profile, we evaluated the effects of chronic administration on amyloid burden and cognitive function in aged APP/PS1 mice. Nineteen 12-month-old APP/PS1 mice received intraperitoneal injections three times per week for eight consecutive weeks. Eleven mice received PBS as the vehicle control, and eight mice received P33 at a dose of 200 µg/kg/day. Three days after the final administration, the mice underwent behavioral testing and were then sacrificed for tissue collection. The brains were processed for biochemical and histological analyses. One hemisphere was homogenized for ELISA and Western blot analyses, and the other was immersion-fixed in 4% phosphate-buffered paraformaldehyde for immunohistochemistry. In parallel, the major peripheral organs (the lungs, kidneys, liver, and spleen) of each animal were collected for histopathological assessment.

ELISA analysis revealed a substantial decrease in cerebral Aβ1–42 levels in P33-treated mice versus vehicle-treated controls (−35.60%, p = 0.038; see Figure 5A). Consistent with these findings, immunohistochemical analysis using the 4G8 antibody showed a marked reduction in Aβ-immunoreactive plaques in both the neocortex (CTX) and hippocampus (HIP) of P33-treated mice (see Figure 5B-D).

**Figure 5.**
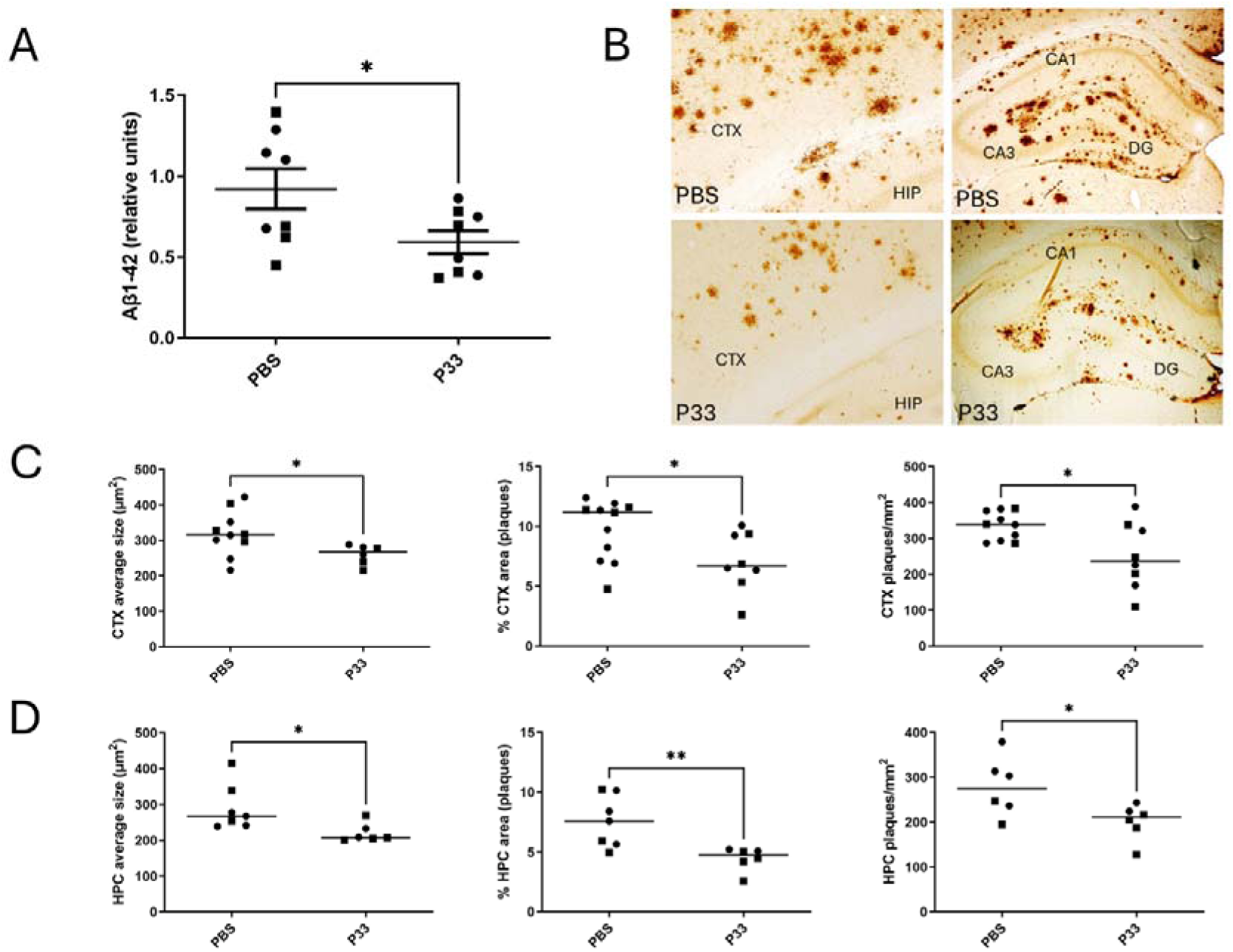
Chronic administration of P33 reduces cerebral Aβ levels and amyloid plaque burden in APP/PS1 mice. Eight 12-month-old APP/PS1 mice were treated intraperitoneally with P33 (200 µg/kg/day), while eleven age-matched APP/PS1 mice received PBS as vehicle control. Injections were administered three times per week for eight consecutive weeks. (A) Aβ1-42 levels quantified by ELISA in soluble hemibrain extracts. (B) Representative immunohistochemistry images showing 4G8-positive amyloid plaques in the CTX (left) and HIP (right) of 14-month-old mice, illustrating reduced plaque burden following P33 treatment compared to PBS control. (C) Quantification of amyloid plaques in the CTX shown in B, including average plaque size (μm²), percentage of area occupied by plaques, and plaque density (plaques/mm²). (D) Quantification of HIP amyloid plaques shown in B using the same parameters: average plaque size, plaque area percentage, and plaque density. Each dot represents one animal; shape code indicates sex (square: male; circle: female). Asterisks indicate statistically significant differences (\**p* < 0.05, \*\**p* < 0.01, Student’s *t*-test). Abbreviations in B: CTX = Neocortex, HIP = Hippocampus, CA1-3 = *Cornus ammonis* 1-3, DG = *Dentate gyrus*.

Quantitative analysis revealed a significant decrease in amyloid plaque burden after P33 treatment. In the CTX, average plaque size was reduced by approximately 18.83% (p = 0.046) and plaque number by 25.30% (p = 0.019), and plaque area fraction or plaque density (the percentage of the relative area occupied by Aβ-immunoreactive plaques) by 27.31% (p = 0.037) (Figure 5C). Similarly, in the HIP, plaque size was reduced by 24.04% (p = 0.031), plaque number by 28.04% (p = 0.032), and plaque density by 41.30% (p = 0.0076) (Figure 5D).

Importantly, the reduction in Aβ levels and senile plaque burden observed following chronic P33 administration correlated with significant improvements in memory performance in APP/PS1 mice, as assessed by the novel object recognition (NOR) test (Figure 6) [50]. P33-treated APP/PS1 mice displayed a significantly higher interest index for the novel object than PBS-treated controls, indicating enhanced recognition memory (Figure 6).

**Figure 6.**
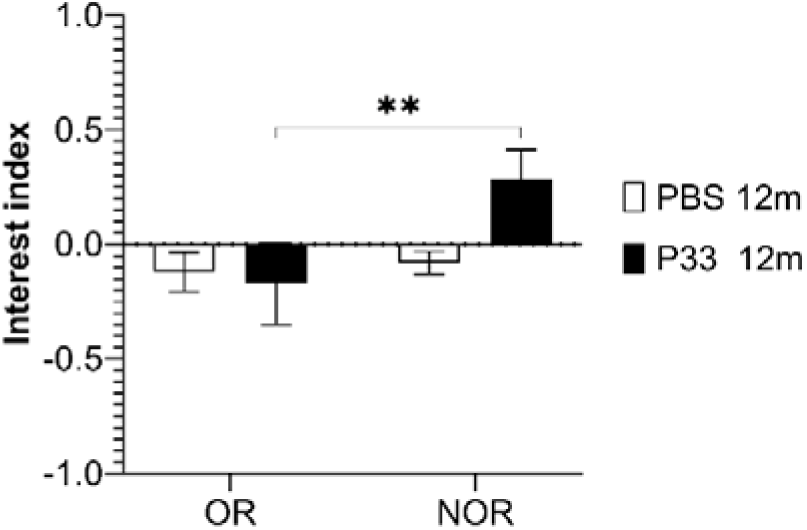
Chronic P33 treatment improves recognition memory in APP/PS1 mice. **Recognition memory of** twelve-month-old APP/PS1 mice chronically treated with P33 or PBS was evaluated using the novel object recognition (NOR) test. On day 1 (object recognition, OR), mice were exposed to two identical objects. On day 2 (novel object recognition, NOR), one familiar object was replaced with a novel object. The interest index was calculated as (touches to novel object − touches to familiar object) / total touches. P33-treated mice displayed a significantly higher interest index on day two (NOR phase) compared to PBS-treated controls, indicating improved recognition memory. Data are presented as mean ± SEM. Asterisks indicate statistically significant differences (\*\**p* < 0.01, two-way ANOVA).

To evaluate the long-term safety of P33, we performed histological analyses of major peripheral organs following chronic treatment. Hematoxylin-eosin staining of the livers, spleens, lungs, and kidneys of randomly selected animals revealed no pathological alterations or tissue lesions in P33-treated mice compared to the control group (Figure 7A).

**Figure 7.**
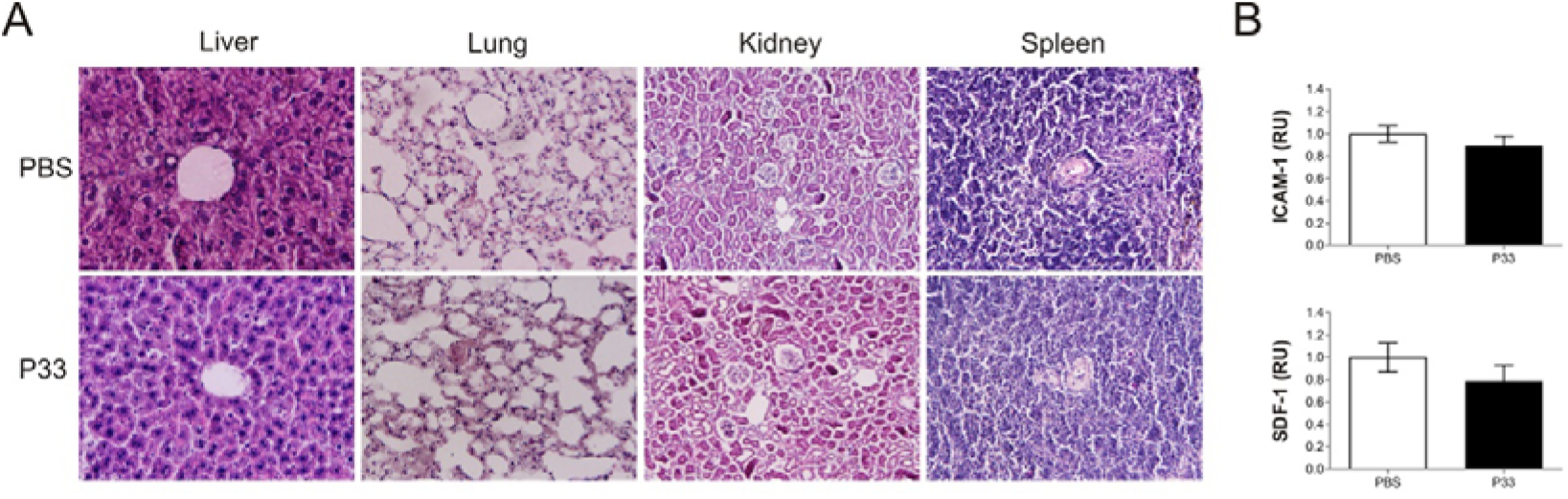
Chronic P33 administration does not induce peripheral toxicity or neuroinflammatory responses in APP/PS1 mice. (A) Representative hematoxylin-eosin staining sections of liver, lung, kidney, and spleen from APP/PS1 mice following chronic P33 or PBS administration, showing no detectable histopathological alterations. Magnification: 200× for lungs and kidney; 400× for liver and spleen. (B) Densitometric histograms of chemiluminescent signals detected on nitrocellulose membranes, representing the expression levels of intercellular adhesion molecule-1 (ICAM-1) and stromal cell-derived factor-1 (SDF-1) in brain extracts, respectively. Data are presented as mean ± SEM.

Additionally, we assessed potential treatment-induced neuroinflammation by cytokine profiling in brain extracts. Only intercellular adhesion molecule 1 (ICAM-1) and stromal cell-derived factor 1 (SDF-1) were detected, with no significant differences observed between P33- and vehicle-treated animals (Figure 7B and Supplementary Figure 3). These results indicate that chronic P33 administration does not induce detectable peripheral toxicity or neuroinflammatory responses.

## 4. Discussion and conclusions

Alzheimer’s disease is a complex, multifactorial neurodegenerative disorder. Its molecular pathology involves the interplay of amyloidogenic processing, tau dysfunction, neuroinflammation, and synaptic failure. This complexity highlights the necessity of molecular tools that can modulate disease-relevant pathways without requiring complete target inhibition or extensive systemic exposure.

Among amyloid-centered strategies, modulation of β-site amyloid precursor protein-cleaving enzyme 1 (BACE1) has been extensively explored due to its pivotal role in the initiation of the amyloidogenic processing of amyloid precursor protein (APP). Significant medicinal chemistry efforts have produced potent BACE1 small-molecule inhibitors, however their clinical development has been hindered by limited efficacy and liabilities related to the mechanism of action, including off-target effects, excessive target suppression, and interference with essential physiological substrates. These outcomes highlight the disadvantages of achieving sustained, complete inhibition of the enzyme in the central nervous system and suggest that alternative approaches to modulating the pathway would be worthwhile.

In this study, we investigated the use of cis-γ-amino-L-proline-derived peptides as modulators of amyloidogenic processing. These γ-peptides are characterized by a non-natural scaffold that exhibits high metabolic stability, tunable charge distribution, and modular side-chain functionalization. Using a phenotypic screening approach in primary neuronal cultures, we identified peptide **P33** as a lead γ-peptide capable of reducing endogenous Aβ production with minimal cytotoxicity. Comparative analysis of structurally related analogues revealed qualitative structure–activity relationships, highlighting the importance of charge density, quaternary ammonium groups, and alternating aromatic/aliphatic side-chain patterns in driving biological activity.

Notably, **P33** reduced Aβ release in neuronal cultures in parallel with a partial reduction in BACE1 activity, while showing minimal effect on BACE2 under the same conditions. Although these data do not prove direct enzymatic inhibition, they support the notion that **P33** modulates amyloidogenic processing in a manner consistent with selective interference at the level of BACE1-associated pathways. Importantly, this modulation does not rely on complete enzyme inhibition, which could be advantageous for preserving the physiological functions of BACE1 substrates involved in synaptic maintenance and neuronal homeostasis.

*In vivo* studies using the APP/PS1 mouse model further supported the biological relevance of this scaffold. The acute administration of **P33** in six-month-old mice reduced Aβ1-42 levels within days, suggesting potential efficacy. Chronic administration of **P33** in 12-month-old mice, representing an advanced stage of the disease, resulted in reduced soluble Aβ levels and amyloid plaque burden. This was accompanied by improved performance in a hippocampal-dependent memory task. Although these results should be interpreted with caution, as they do not definitively establish a pharmacokinetic–pharmacodynamic relationship, they demonstrate that γ-peptides with the appropriate structural features can reach the central nervous system and produce measurable biological effects *in vivo*. Notably, chronic administration of P33 was not associated with overt toxicity, histopathological alterations in peripheral organs, or induction of neuroinflammatory markers. These results support the biocompatibility of this scaffold.

In conclusion, our study establishes cis-γ-amino-L-proline peptides as a metabolically stable and modular scaffolds for the intracellular modulation of amyloidogenic processing. Phenotypic structure-activity relationship analysis shows the potential to fine-tune sequence, charge, and side-chain patterns to achieve selective modulation of BACE1 activity. These γ-peptides provide a versatile chemical platform that will facilitate the future design of intracellular probes and potential therapeutic leads in AD-related research.

## Abbreviations

Standard abbreviations and acronyms recommended by *Journal of Medicinal Chemistry* are used. Others include

ACN: Acetonitrile
Amp: 4-aminoproline
Braak V–VI: Braak staging (stages V–VI)
CGC: cerebellar granular cells
CPPs: Cell-penetrating peptides
CTX: neocortex
DAB: diaminobenzidine
DIPEA: *N*,*N*-Diisopropylethylamine
DIV: days *in vitro*
EDC: *N*-Ethyl-*N’*-(3-dimethylaminopropyl)carbodiimide
ELISA: Enzyme-Linked immunosorbent assay
HBTU: N-[(1H-benzotriazol-1-yl)-(dimethylamino)methylene]-N-methylmethanaminium hexafluorophosphate N-oxide
HE: hematoxylin-eosin
HIP: hippocampus
HOBt: 1-Hydroxybenzotriazole
ICAM-1: Intercellular cell adhesion molecule-1
ICD-10: International Classification of Diseases 10th Revision
IL1β: Interleukin-1β
ipen: isopentyl
MCI: Mild cognitive deficit
NINCDS-ADRDA: National Institute of Neurological and Communicative Disorders and Stroke-Alzheimer’s Disease and Related Disorders Association
NFTs: Neurofibrillary tangles
NOR: novel object recognition
OR: object recognition test
PFA: Paraformaldehyde
phet: phenethyl
RT-qPCR: Reverse transcription quantitative polymerase chain reaction
SDS-PAGE: Sodium dodecyl sulfate–polyacrylamide gel electrophoresis
SEM: Standard error of the mean
SDF-1: Stromal cell-derived factor-1
TAT: Transactivator of transcription
TNFα: Tumor necrosis factor α.

## Acknowledgments

We would like to thank all the members of the J.A.D.R. and M.R. laboratories for their valuable comments on the study. Additionally, we are grateful to M. Segura-Feliu and J. L. Jiménez for their technical support. We also thank Tom Yohannan for editorial advice.

## Author contributions: CRediT

Dayaneth Jácome: Investigation, Methodology, Formal analysis. Marina Pérez: Investigation, Methodology, Writing. Inés Martínez-Soria: Investigation, Methodology, Formal analysis. Laia Lidón: Investigation, Methodology, Formal analysis. Cristina Vergara: Investigation, Methodology, Formal analysis. Daniel Carbajo: Investigation, Methodology, Formal analysis. Ximena Pulido: Investigation, Methodology, Formal analysis. Macarena Sánchez-Navarro: Investigation, Methodology, Formal analysis. Ernest Giralt: Conceptualization, Writing - review & editing. Fernando Albericio: Conceptualization, Writing - review & editing. Miriam Royo: Conceptualization, Funding acquisition, Writing - original draft, Writing - review & editing. Rosalina Gavín: Conceptualization, Formal analysis, Funding acquisition, Investigation, Writing - original draft, Writing - review & editing. José Antonio Del Río: Conceptualization, Writing - original draft, Writing - review & editing.

## Data availability statement

The data that support the findings of this study are available on request from the corresponding authors.

## Supplementary material

**Supplementary Figure 1.**
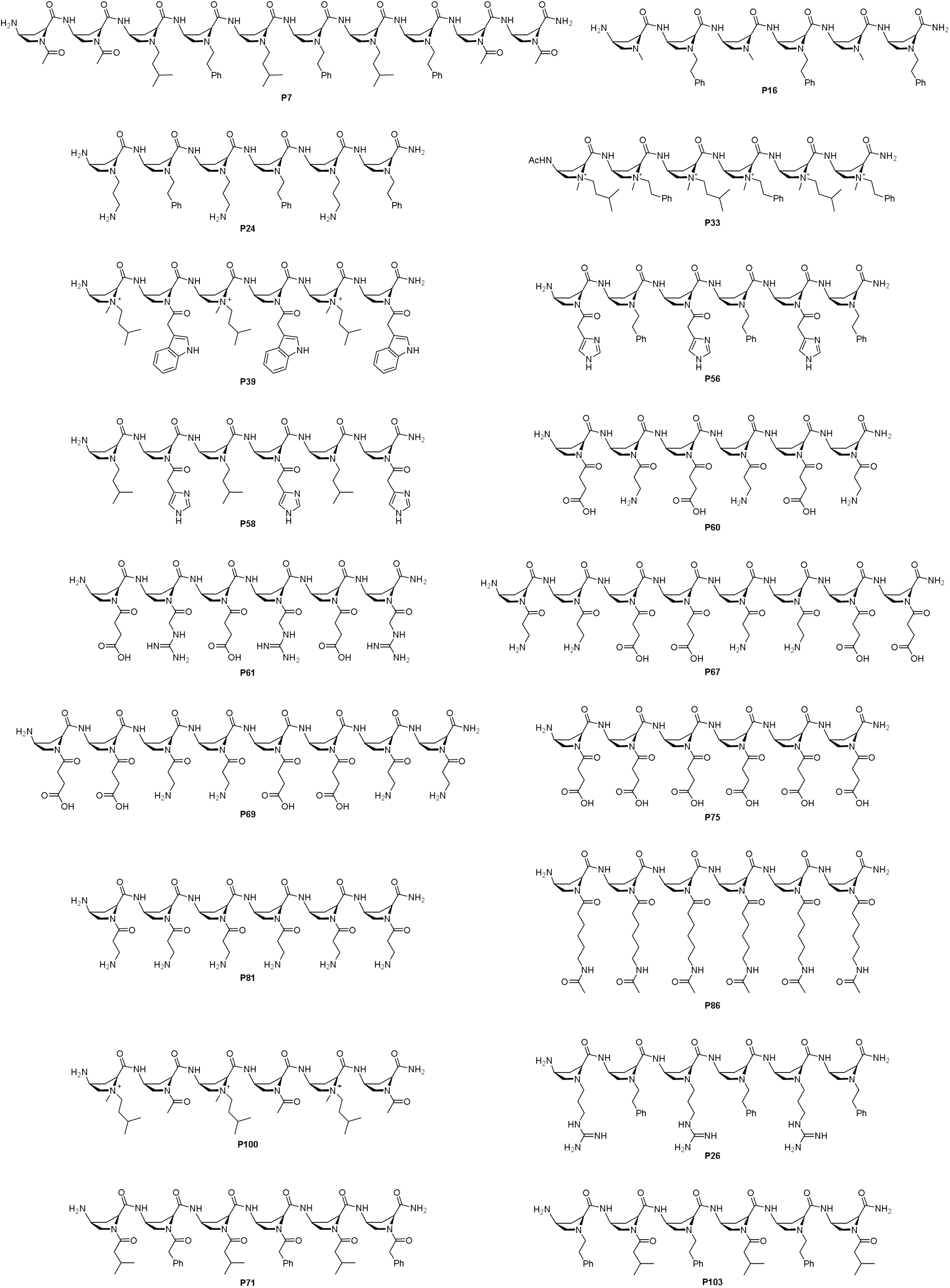
Chemical structure of the eighteen γ-peptides selected by their low cytotoxicity in neurons.

**Supplementary Figure 2.**
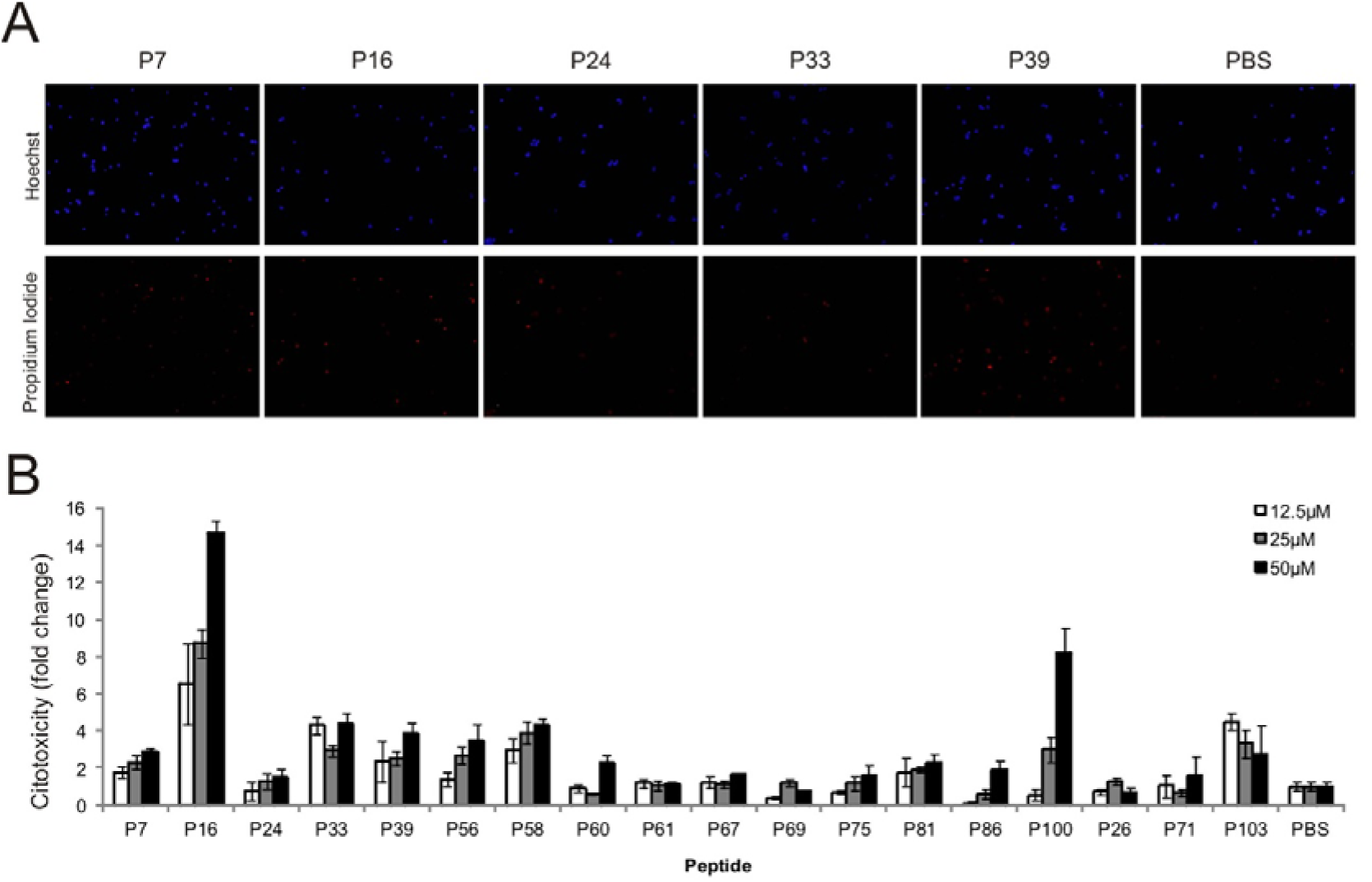
Analysis of the cytotoxicity derived from peptide treatment in neuronal primary cultures. Three different concentrations of the selected γ-peptides - 12.5 μM, 25 μM, and 50 μM - along with vehicle (PBS) were applied to primary cortical neuron cultures for two days following 10 days in vitro (DIV). Cytotoxicity was assessed by propidium iodide uptake (n = 3 independent experiments per γ-peptide and concentration). (A) Representative photomicrographs showing Hoechst and propidium iodide staining of primary cultures one day after treatment with selected γ-peptides at 25 μM, illustrating the number of viable cells under each condition. A dose-dependent trend was observed in most cases, with higher concentrations inducing greater cytotoxicity. (B) Quantitative analysis of cytotoxicity across all peptides at the three tested concentrations. PBS-treated cultures were used as the reference (unit). Results are presented as fold change relative to PBS, showing minimal differences at the lower doses.

**Supplementary Figure 3.**
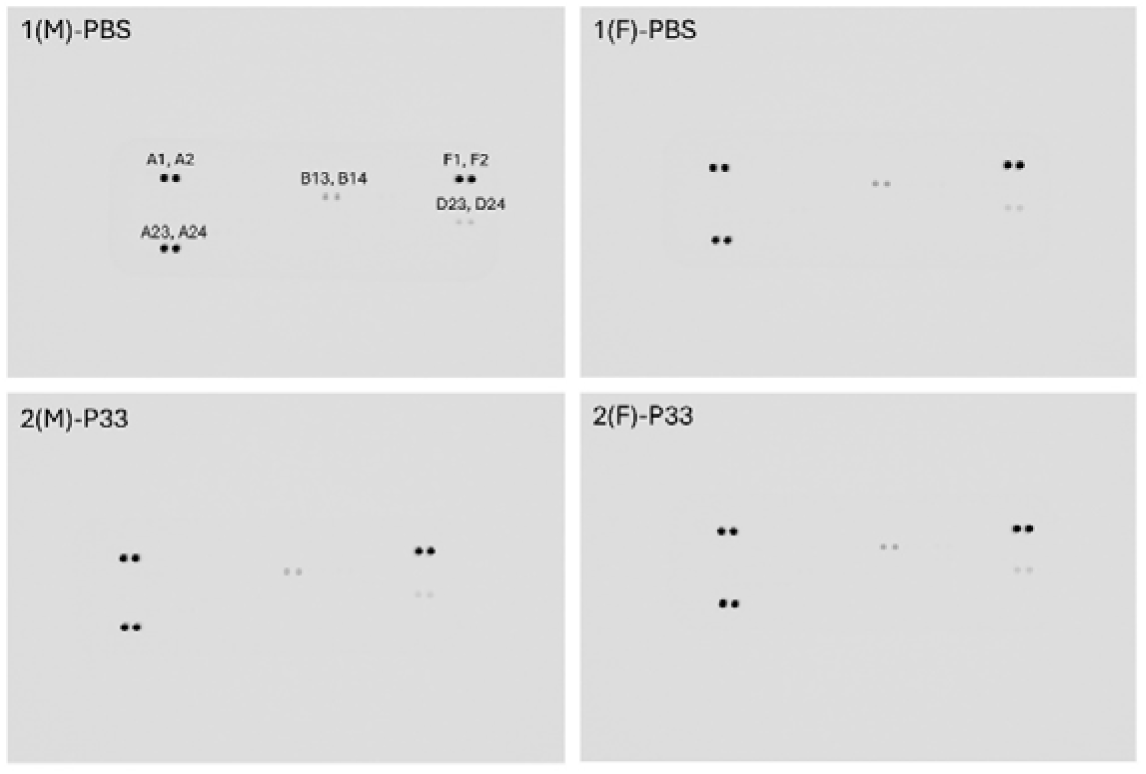
Nitrocellulose membranes after immunochemical cytokine detection. Each membrane shows the cytokine profile of brain protein lysates from one animal. Relative coordinates are shown in membrane from 1(M)-PBS mouse: A1, A2: Reference spot, A23, A24: Reference spot, F1, F2: Reference spot, B13, B14: ICAM-1, D23, D24: SDF-1

**Supplementary Scheme 1.**
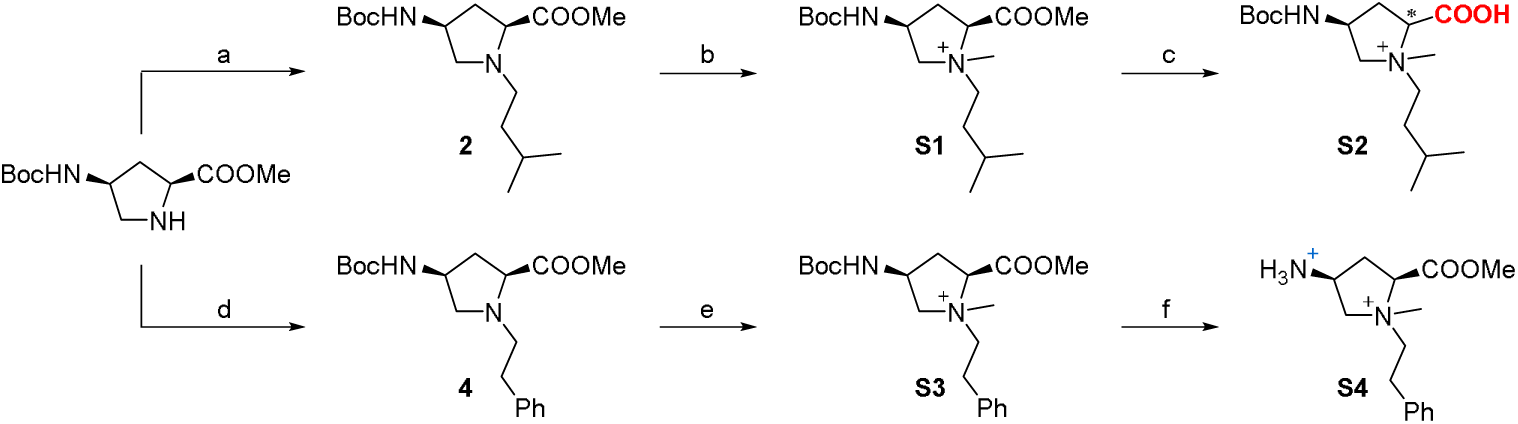
Attempt to synthesize the fully functionalized monomers of P33. ^a^Reagents and conditions: (a) isovaleraldehyde, NaBH_3_CN, DCE, rt, 16 h, 54%; (b) MeI, ACN, rt, 16 h (c) all conditions lead to the epimerization of the α-carbon; (d) phenylacetaldehyde, NaBH_3_CN, DCE, rt, 16 h, 63%; (e) MeI, ACN, rt, 16 h, 77%; (f) HCl, dioxane, rt, 1 h, 99%.

**Supplementary Scheme 2.**
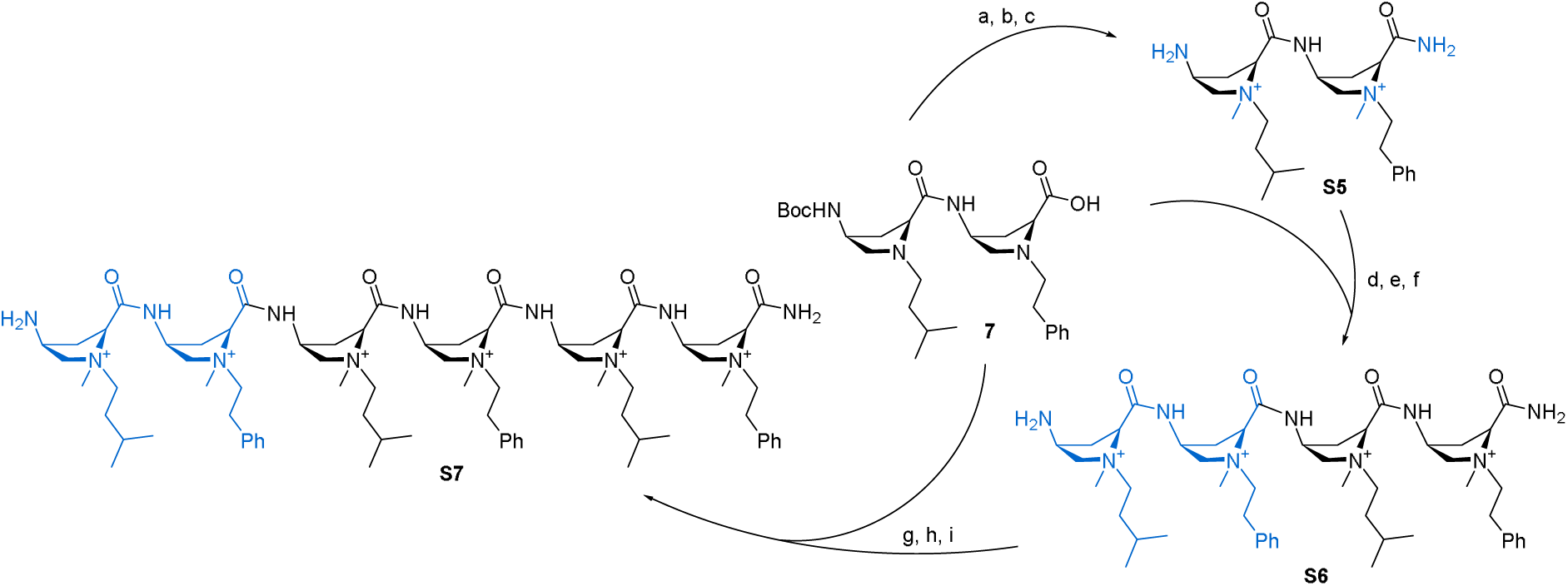
Synthesis of P33 unacetylated precursor from dipeptide 7, Strategy A. ^a^Reagents and conditions: (a) NH_4_Cl, HBTU, Et_3_N, ACN, rt, 1.5 h, 82%; (b) MeI, ACN, 30 LJC, 6 d, 53%; (c) HCl, dioxane, ACN, rt, 1 h, 99%; (d) **7**, EDC·HCl, HOBt·H_2_O, DIPEA, DMF, rt, 4 h, 68%; (e) MeI, ACN, 35 LJC, 7 d, 60%; (f) HCl, dioxane, ACN, rt, 45 min, 99%; (g) **7**, EDC·HCl, HOBt·H_2_O, DIPEA, DMF, rt, 2 h; (h) MeI, ACN, 35 LJC, 15 d, 39% (two steps); (i) HCl, dioxane, ACN, rt, 1 h, 99%.

---

**NMR Spectra of synthesis intermediates and P33**

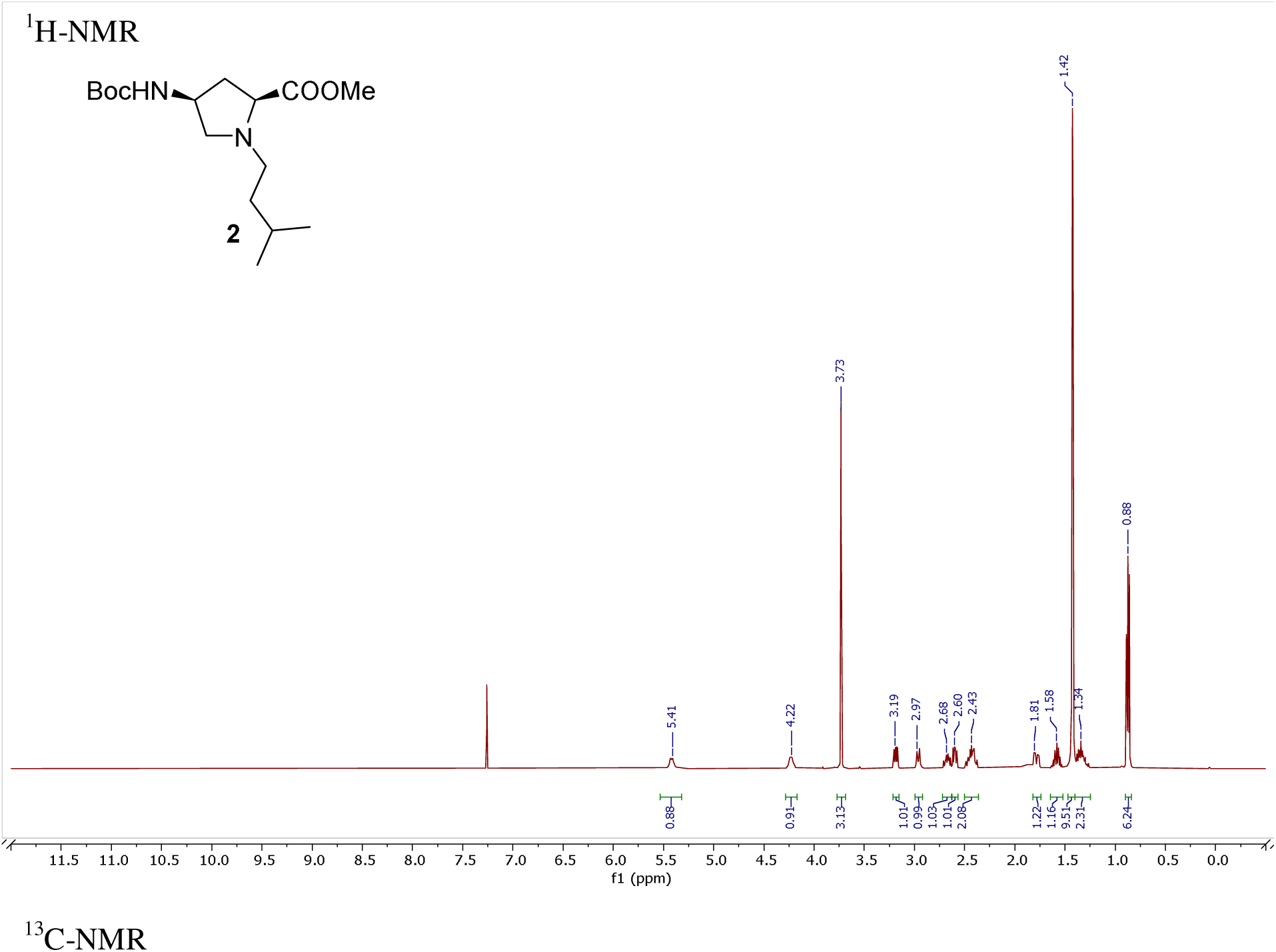

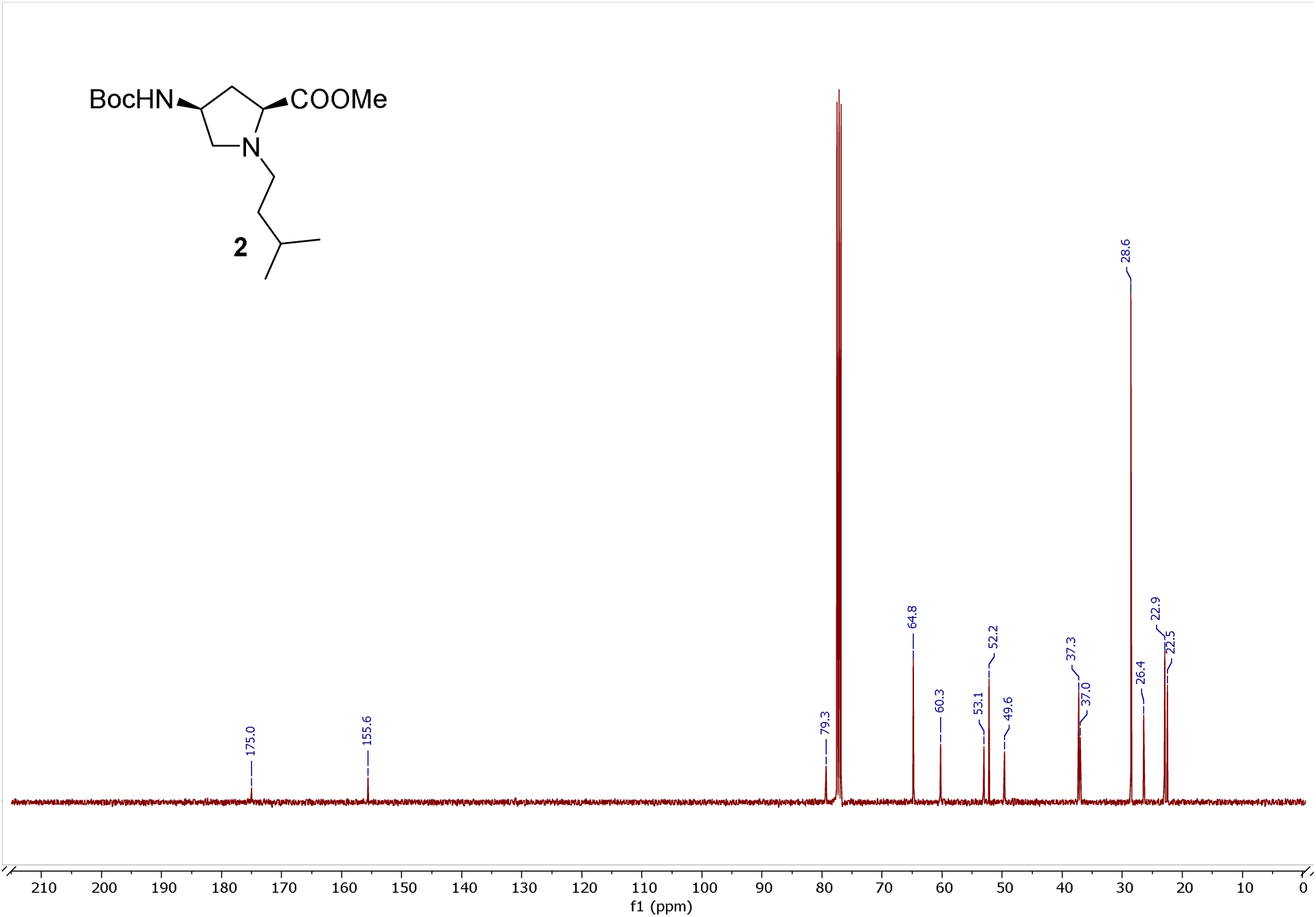

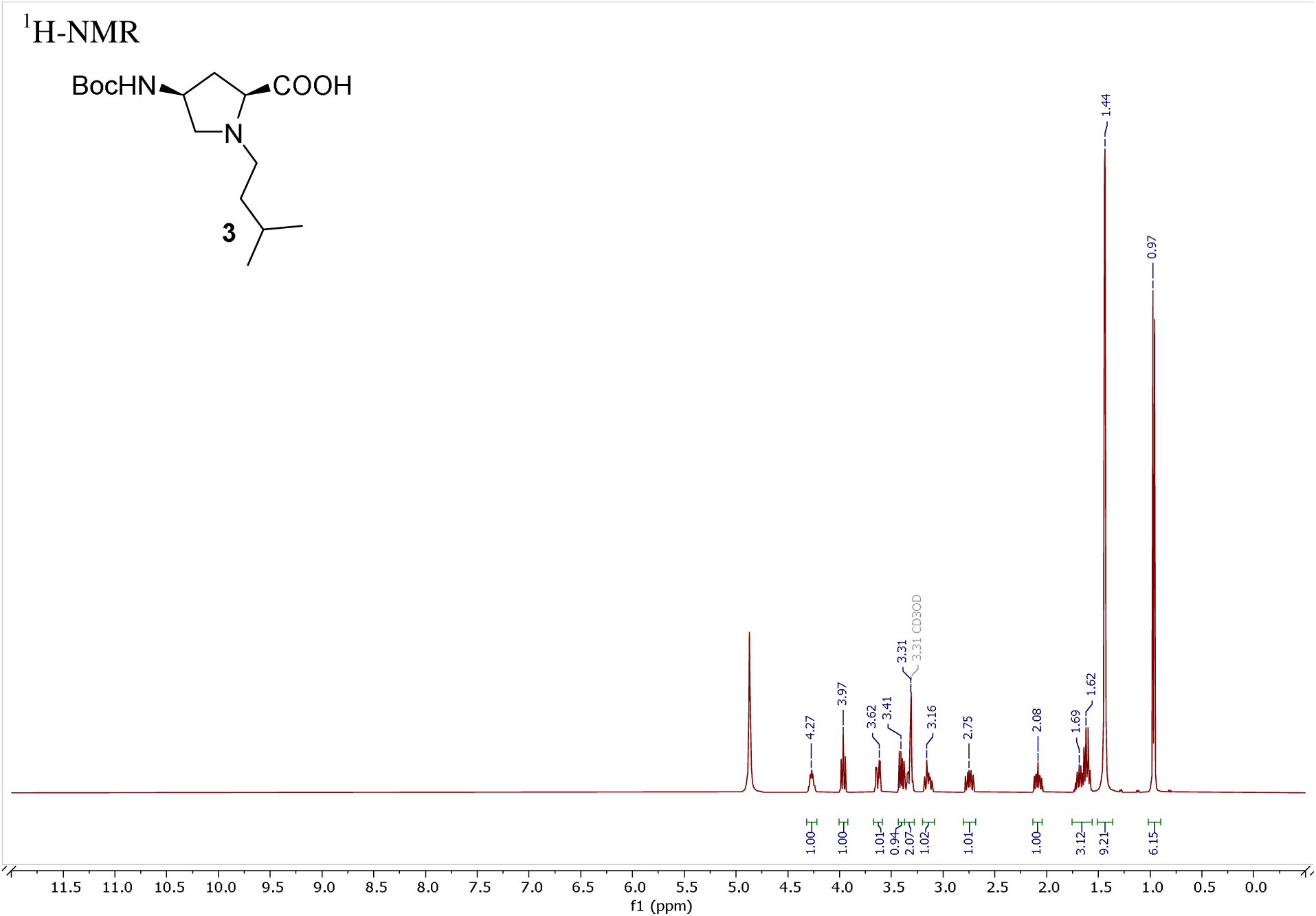

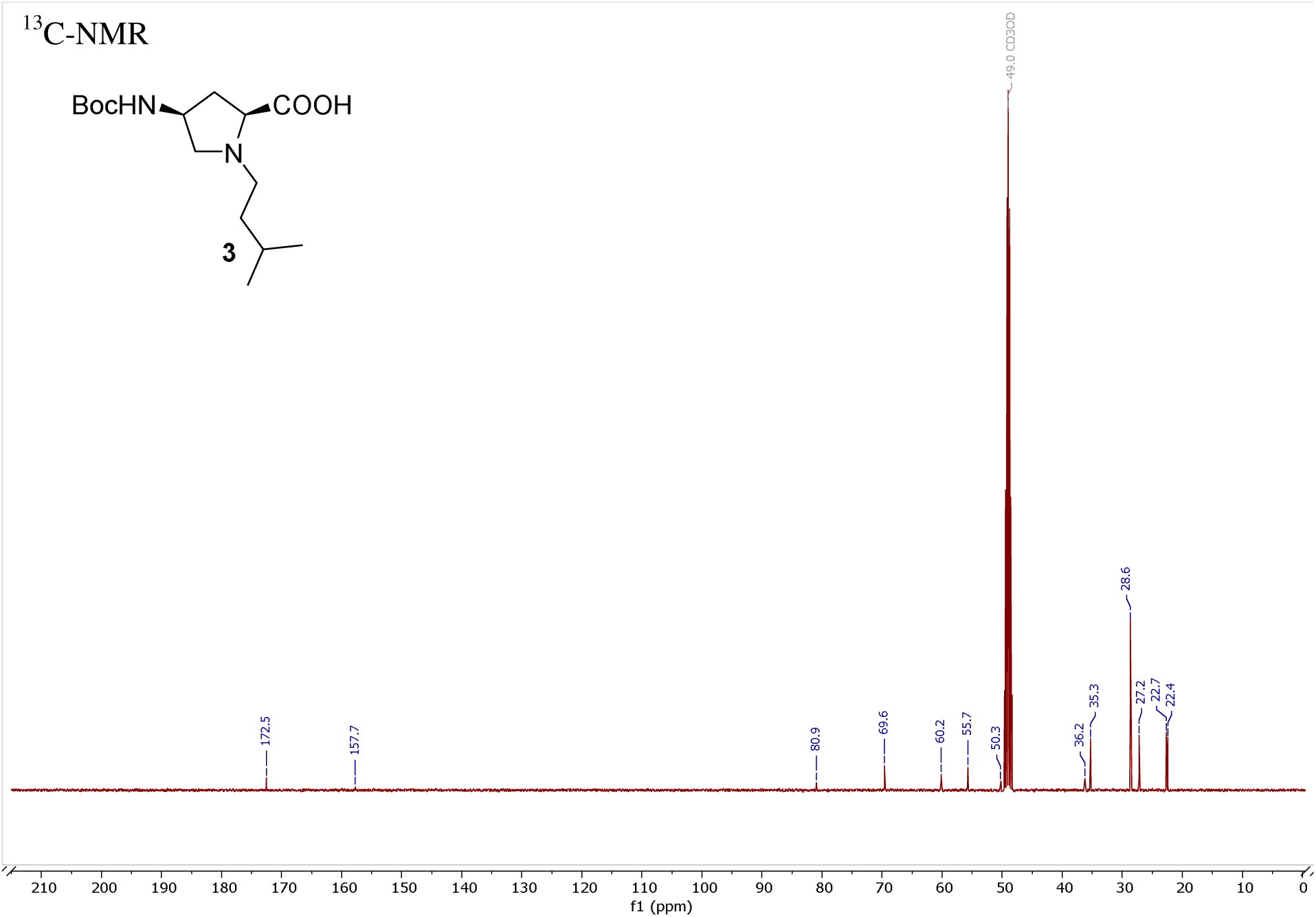

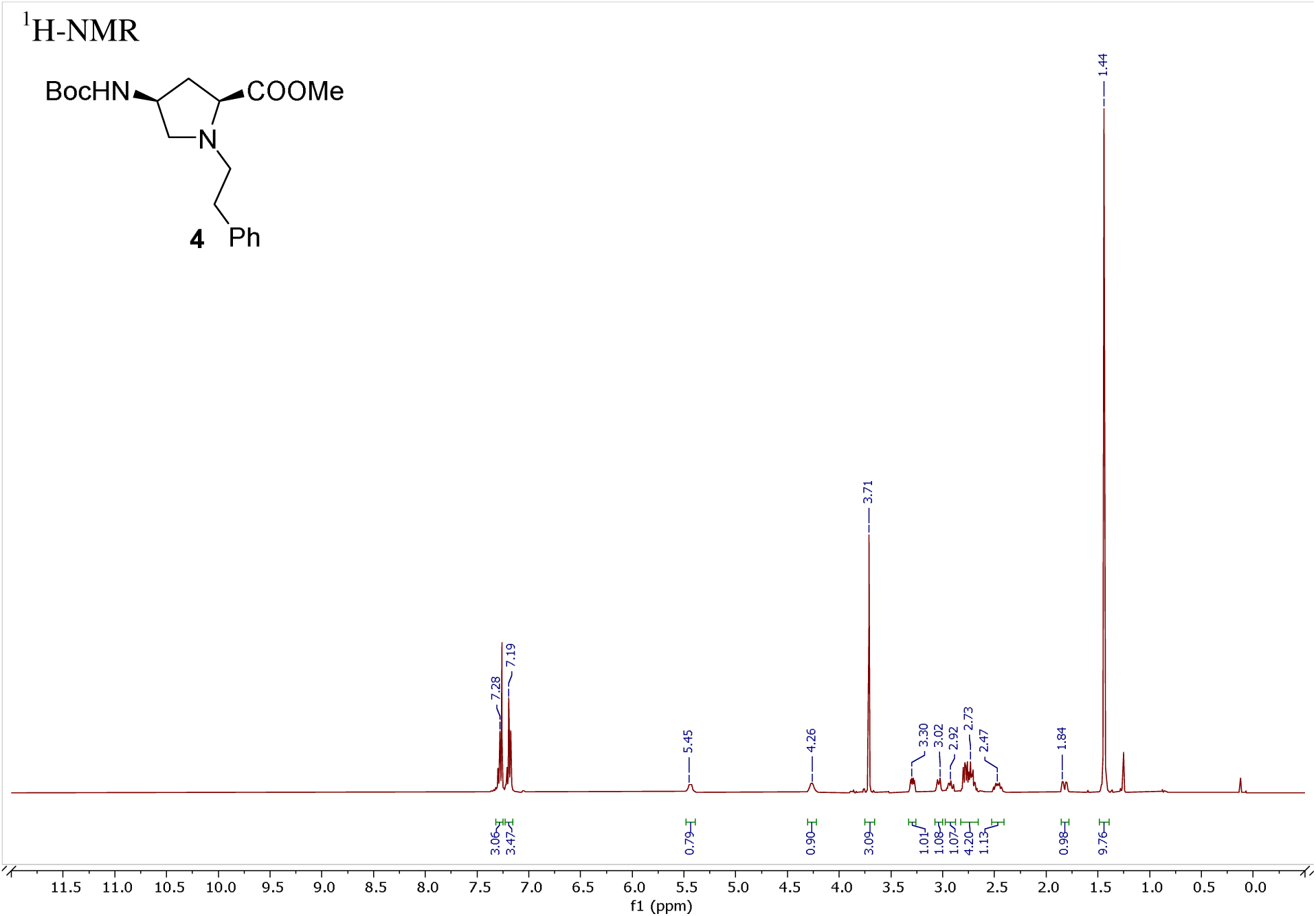

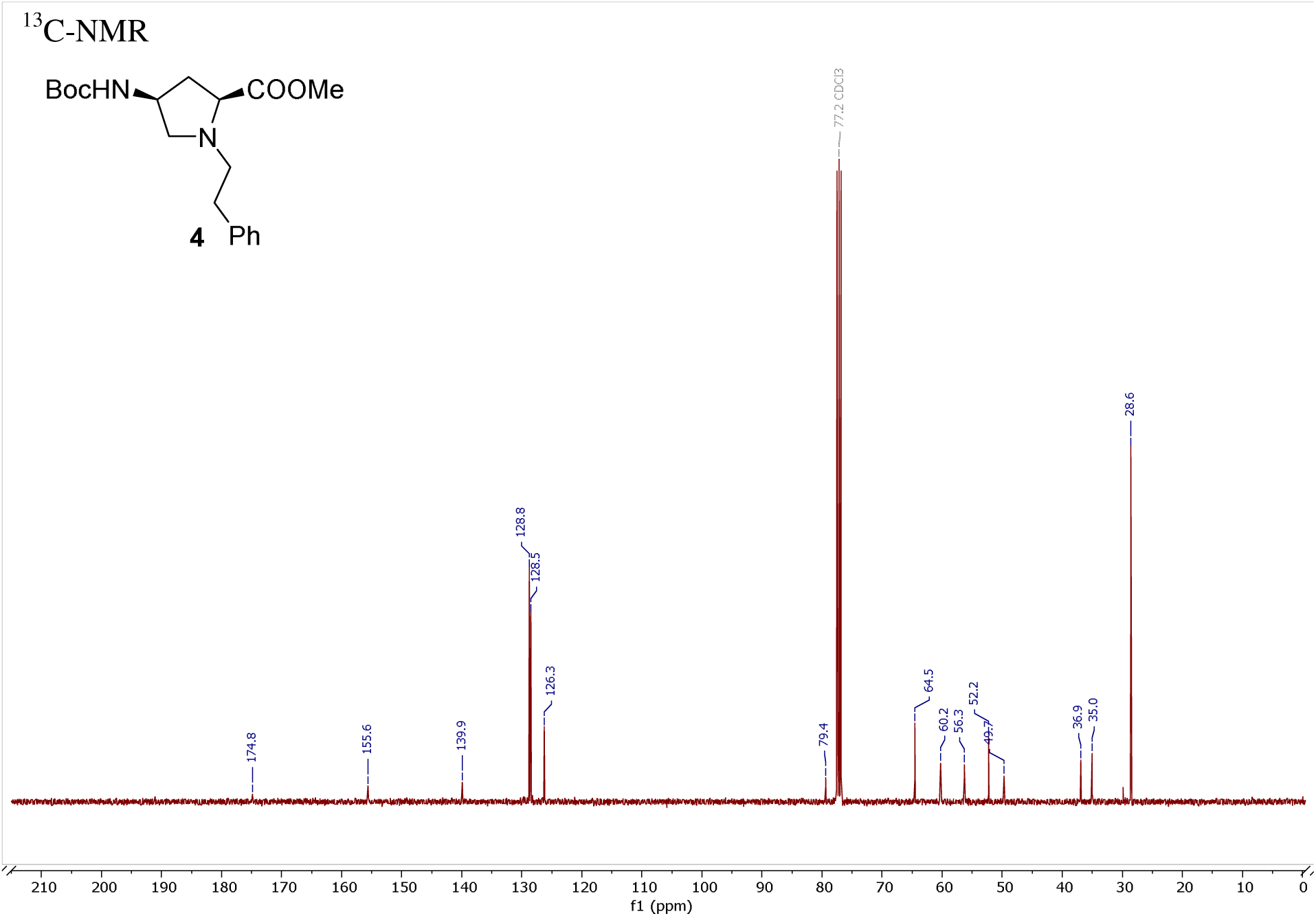

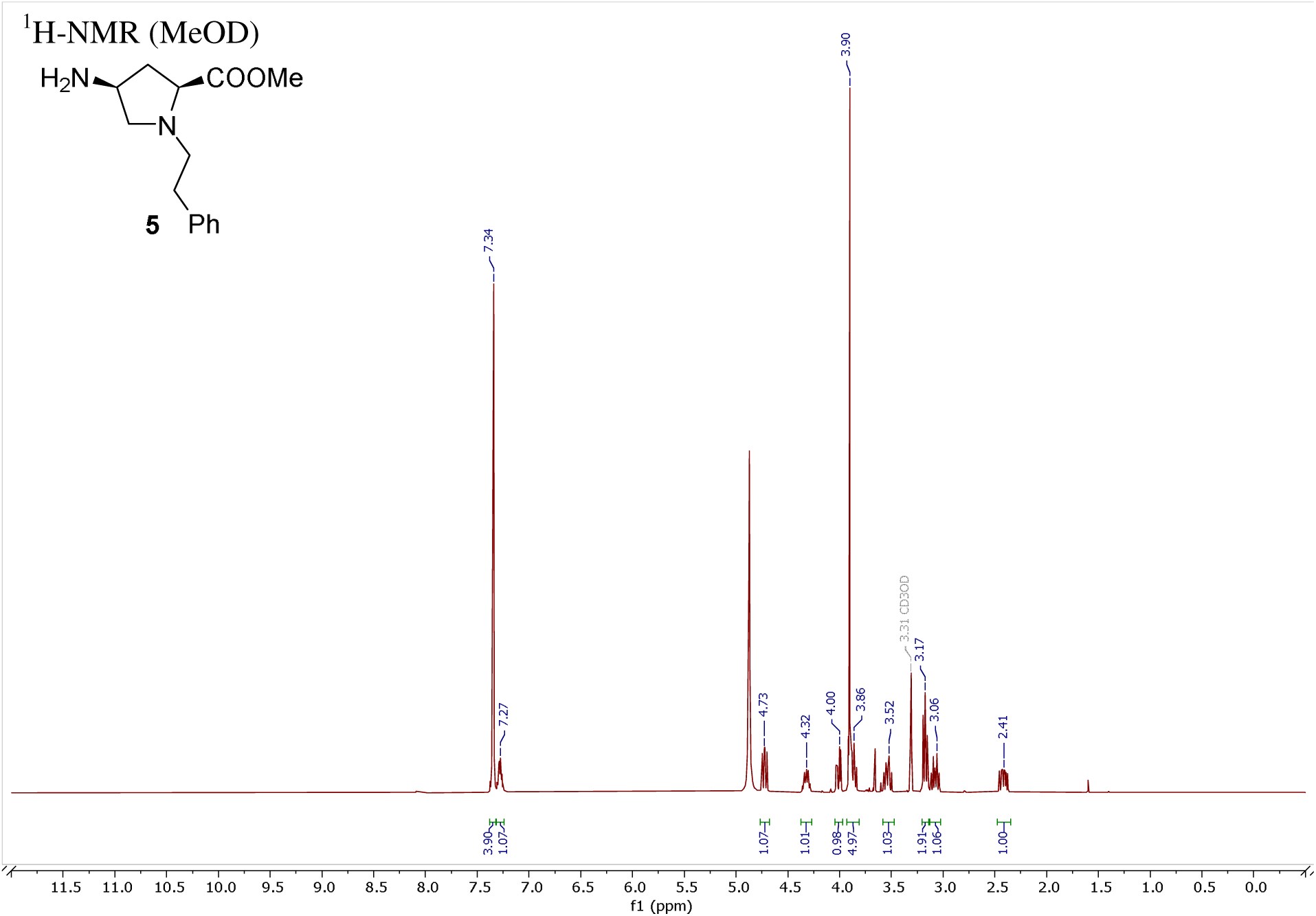

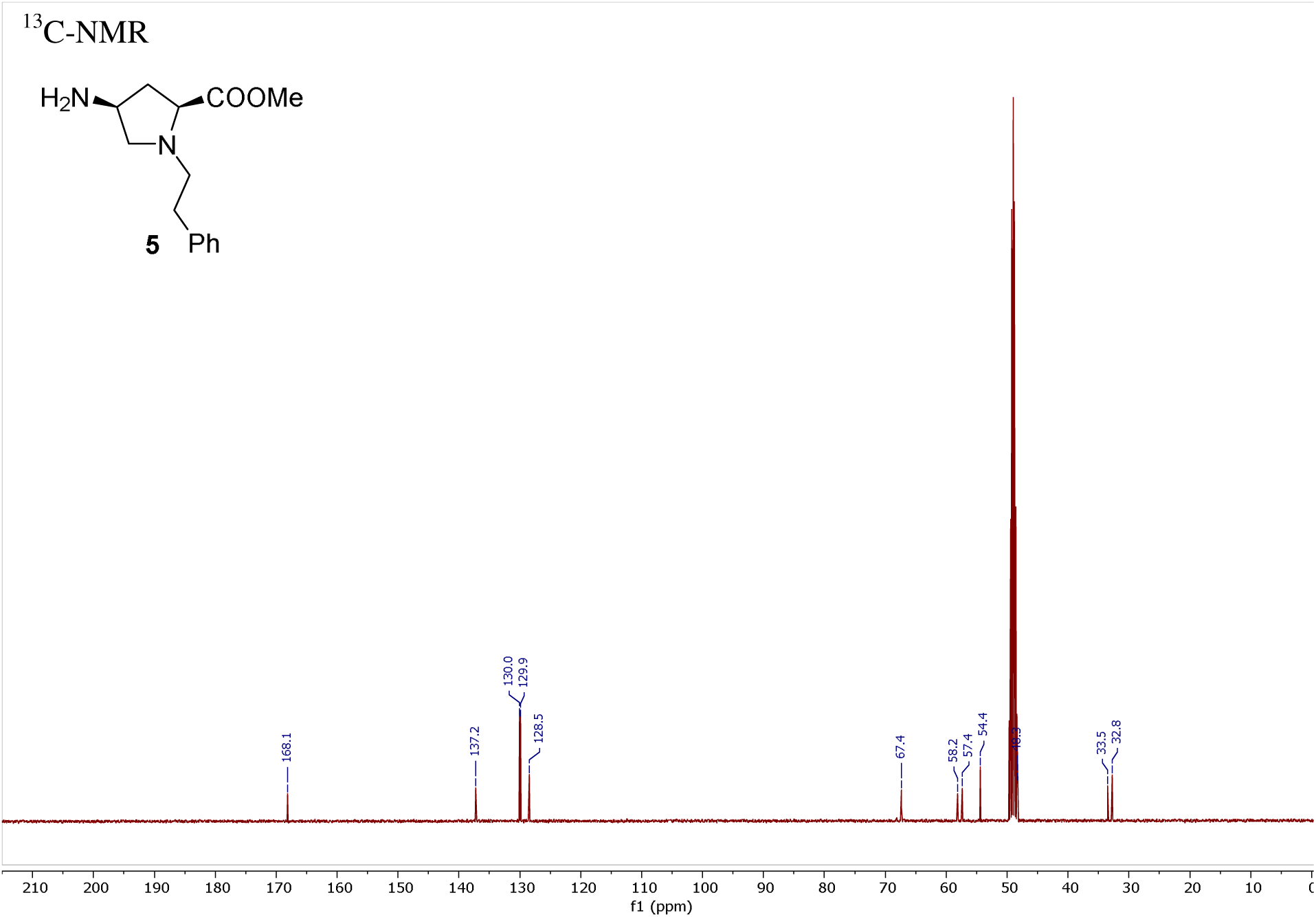

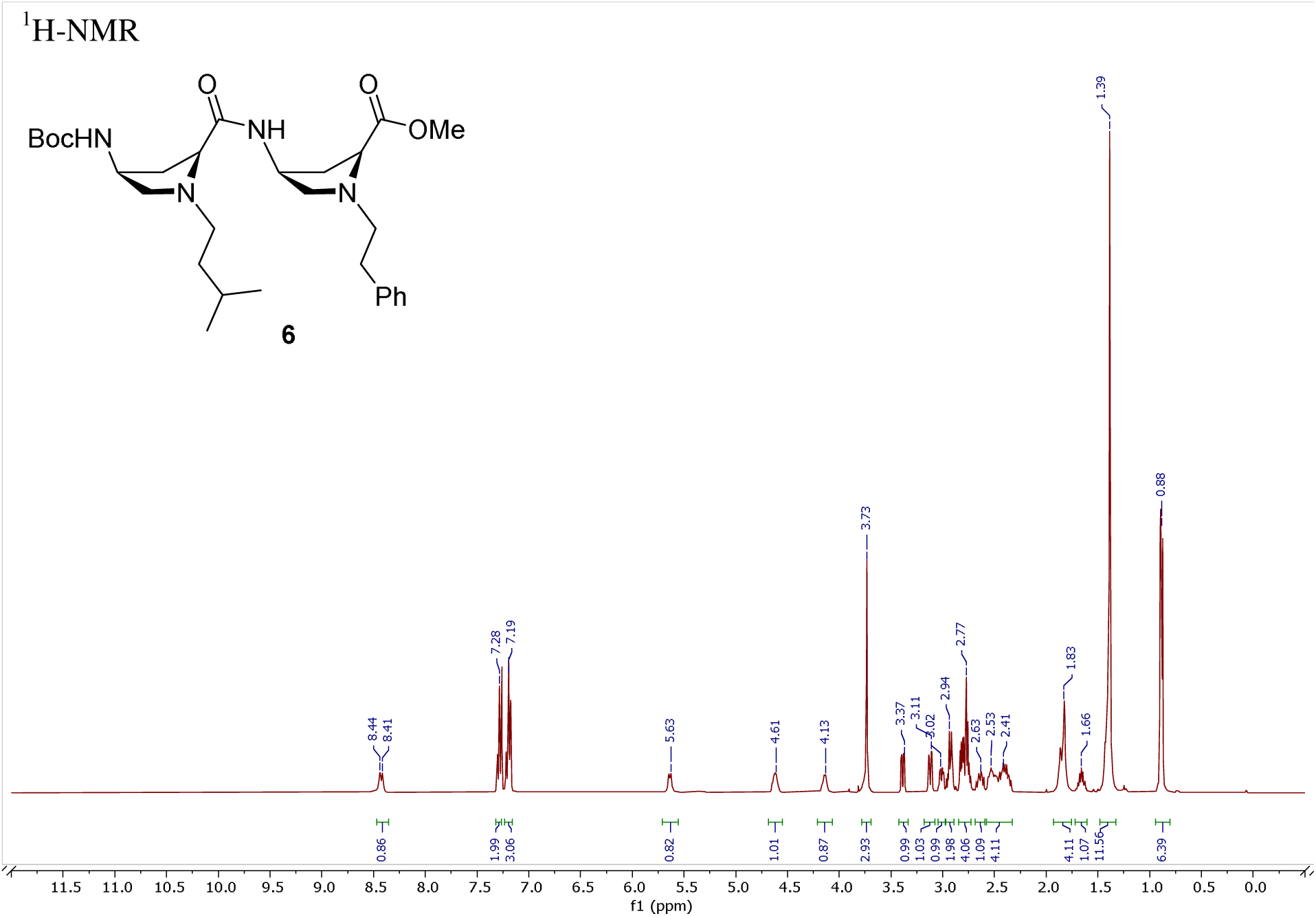

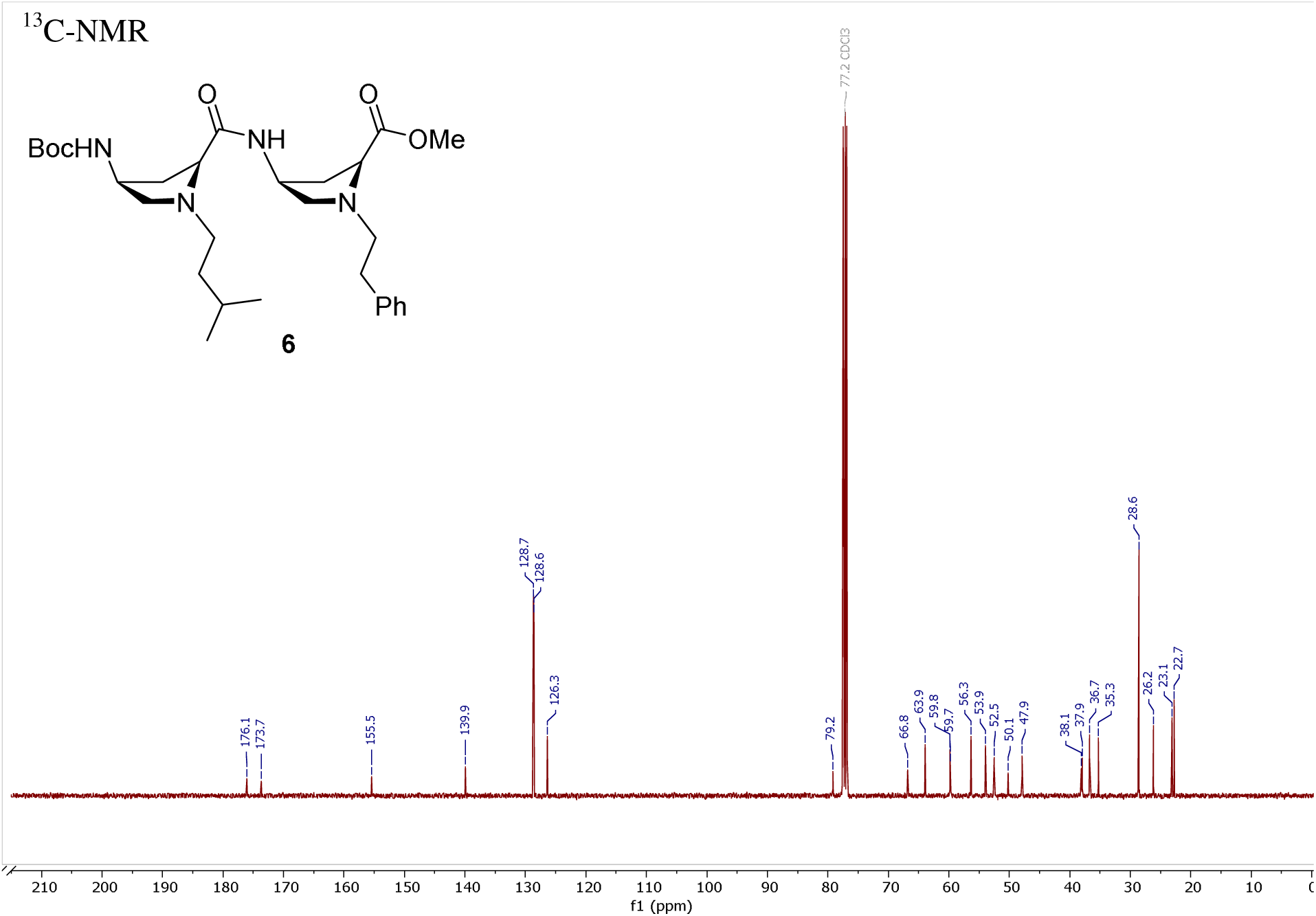

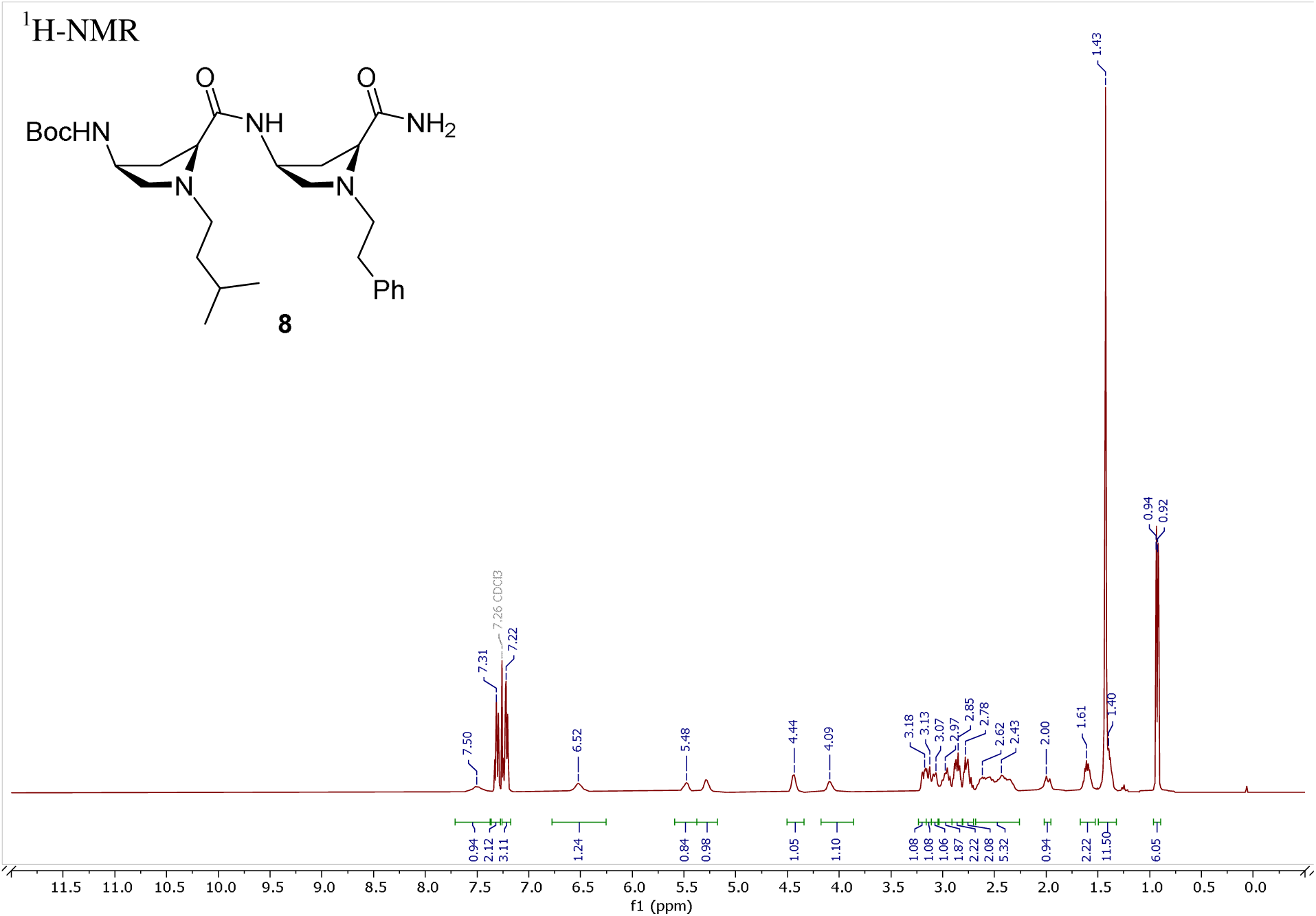

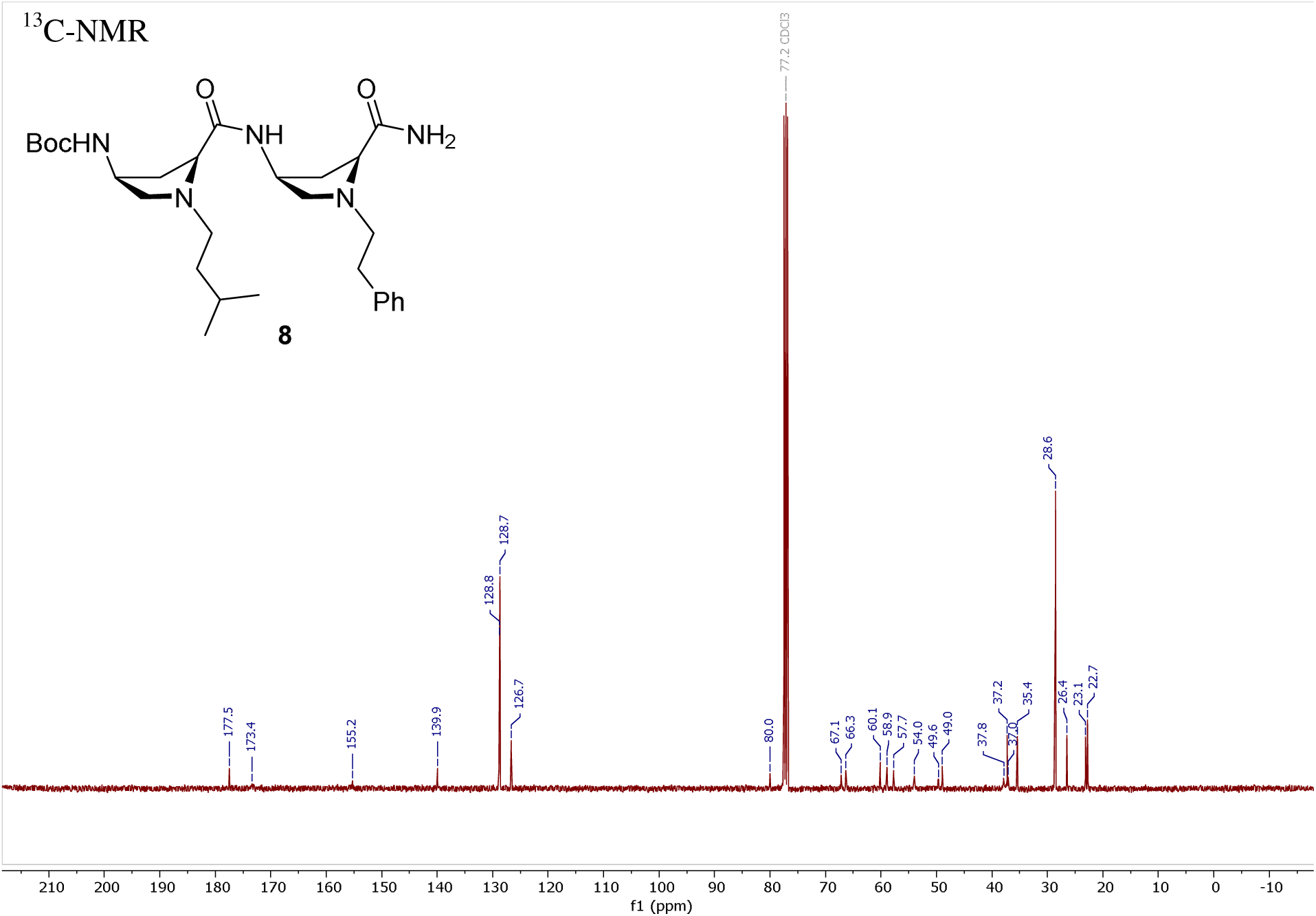

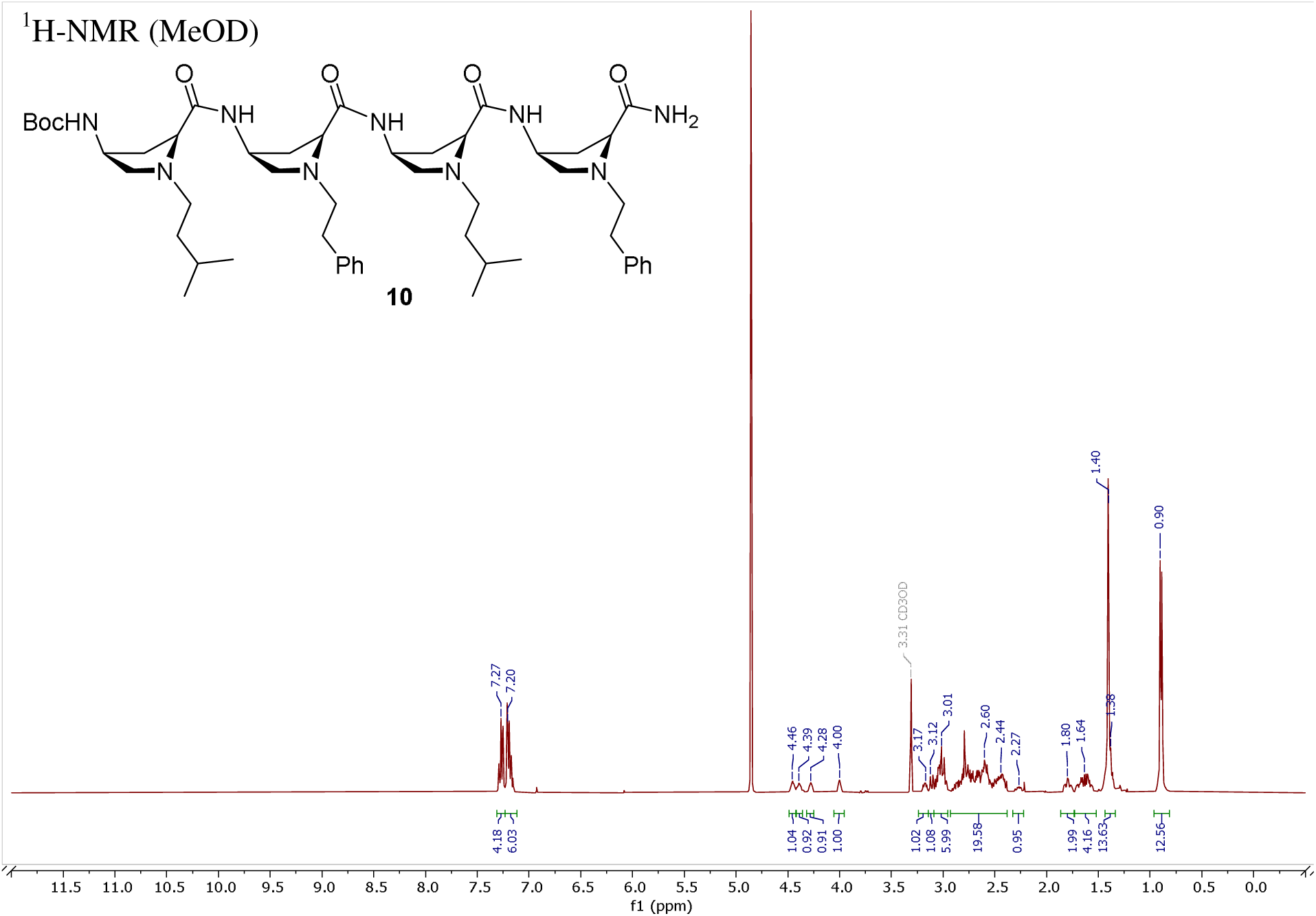

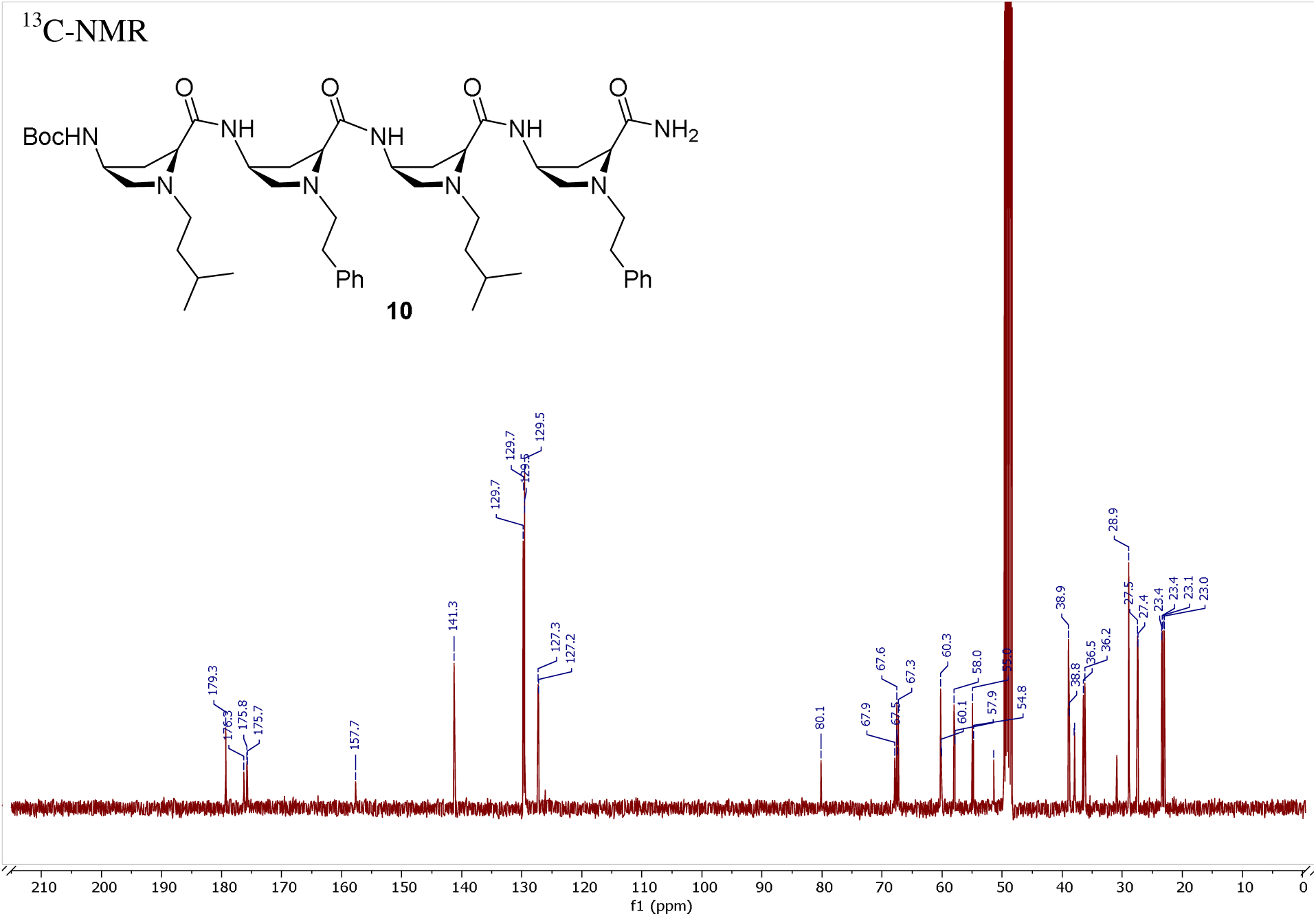

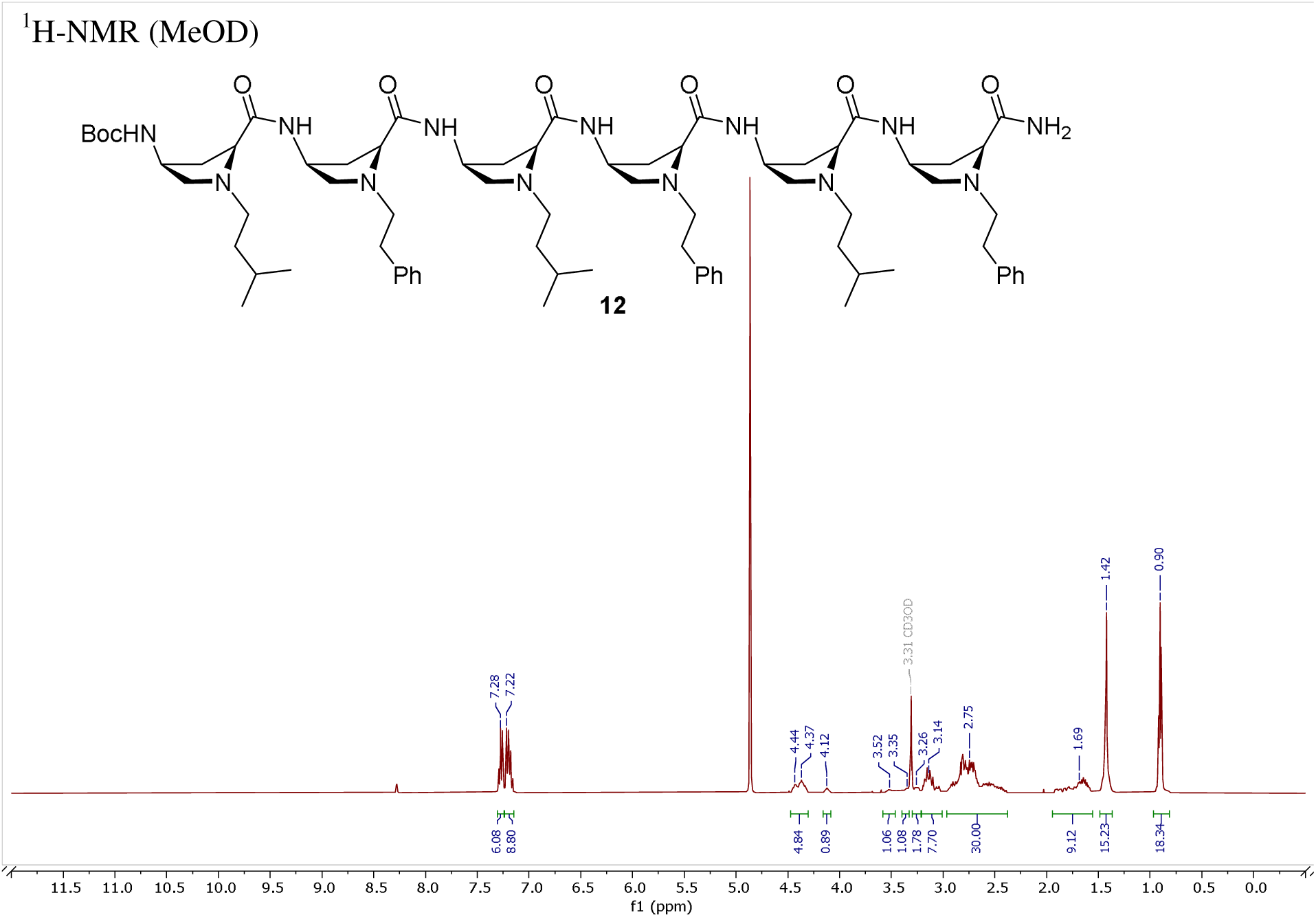

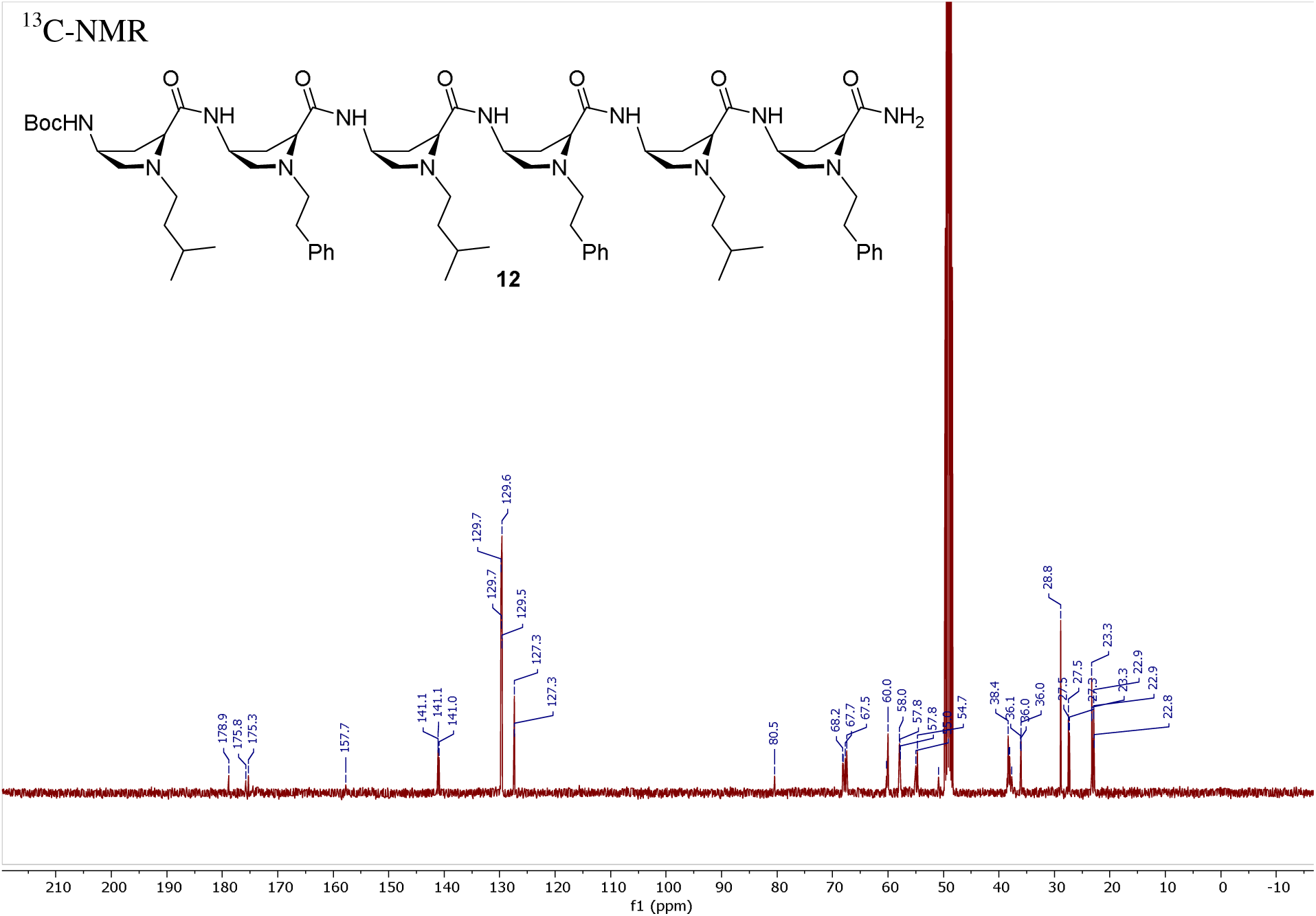

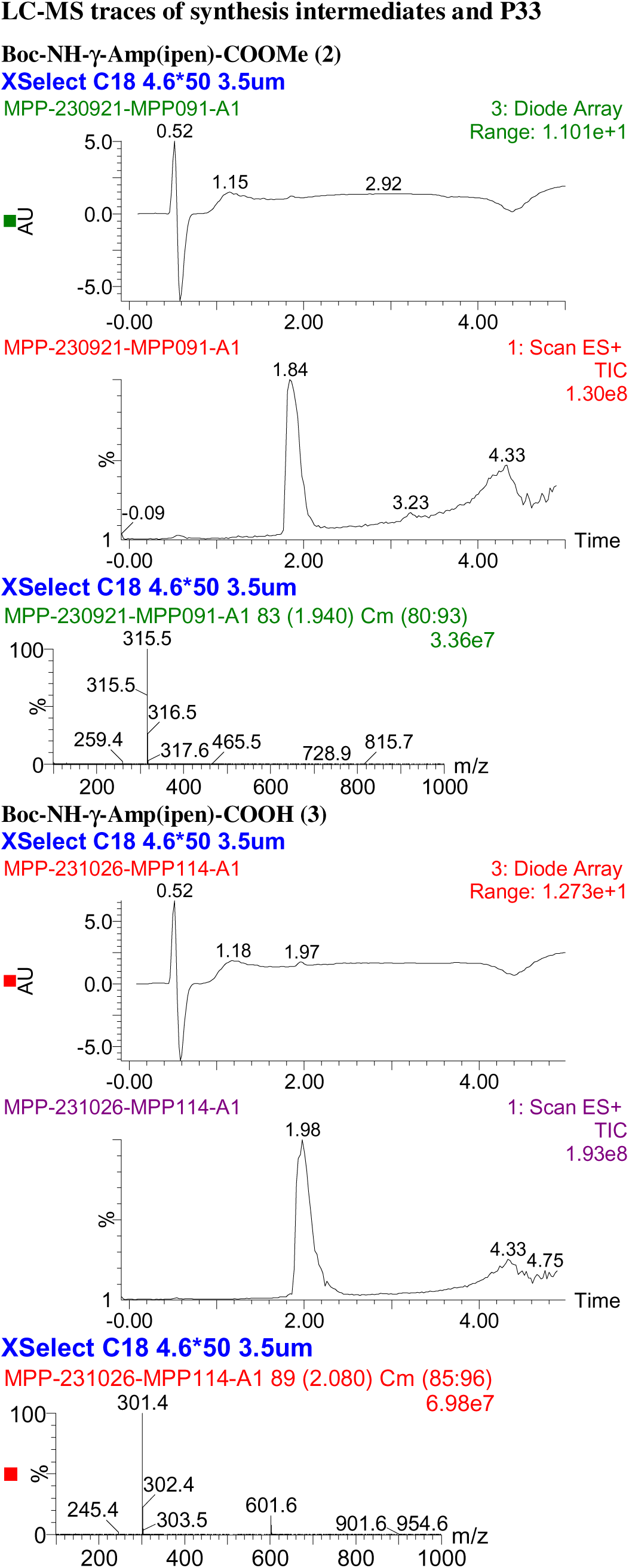

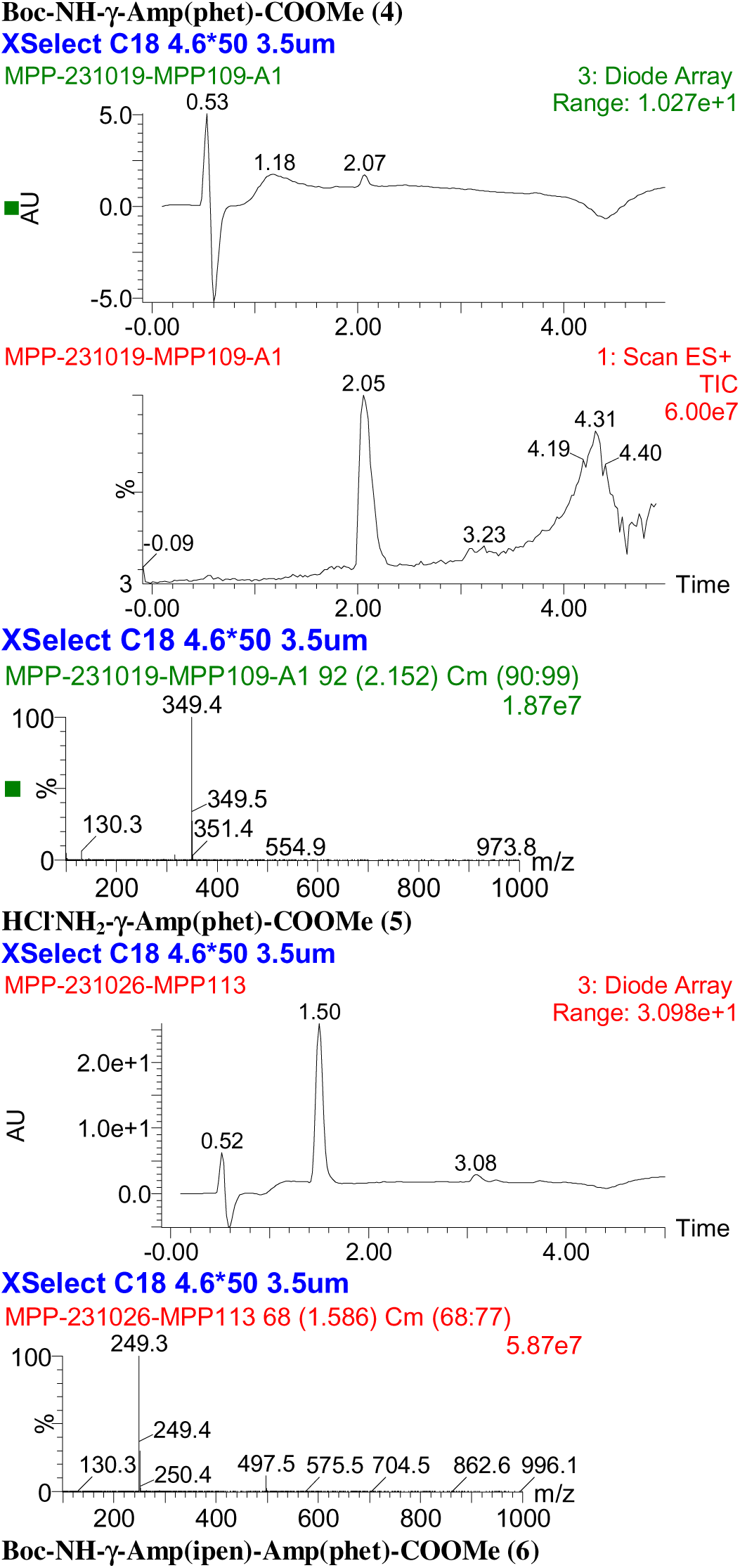

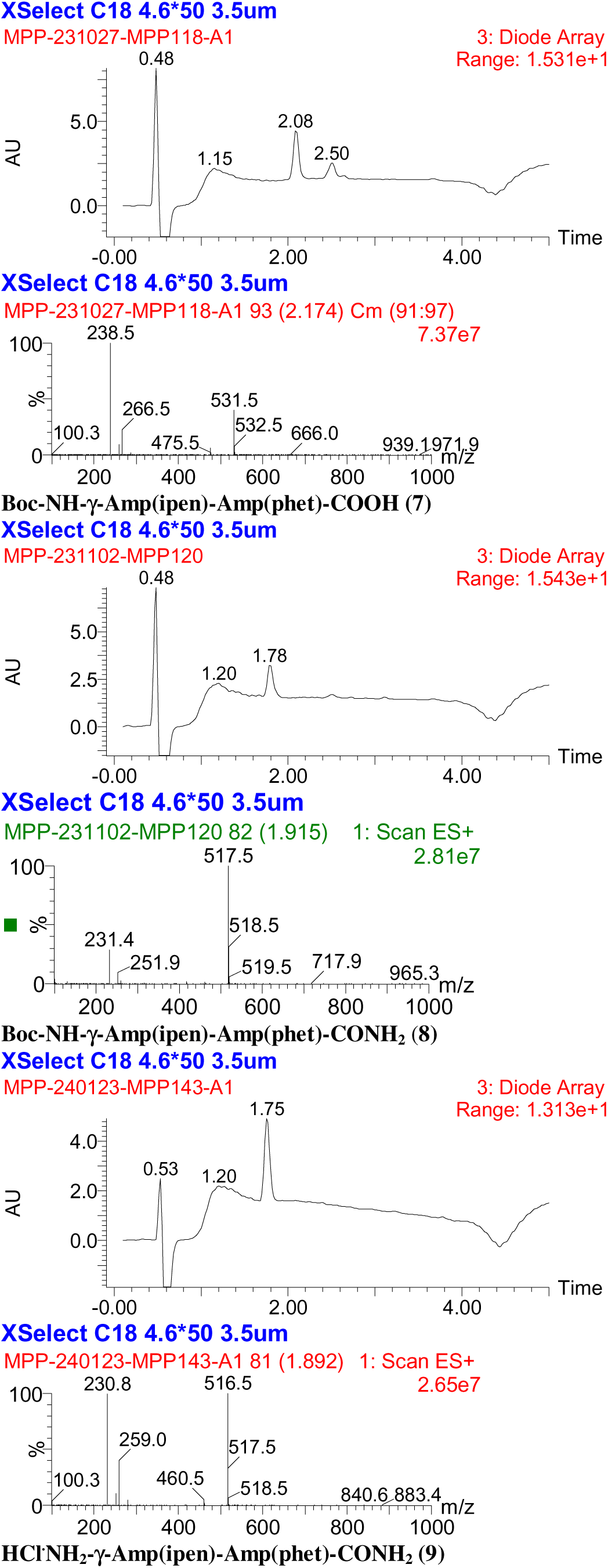

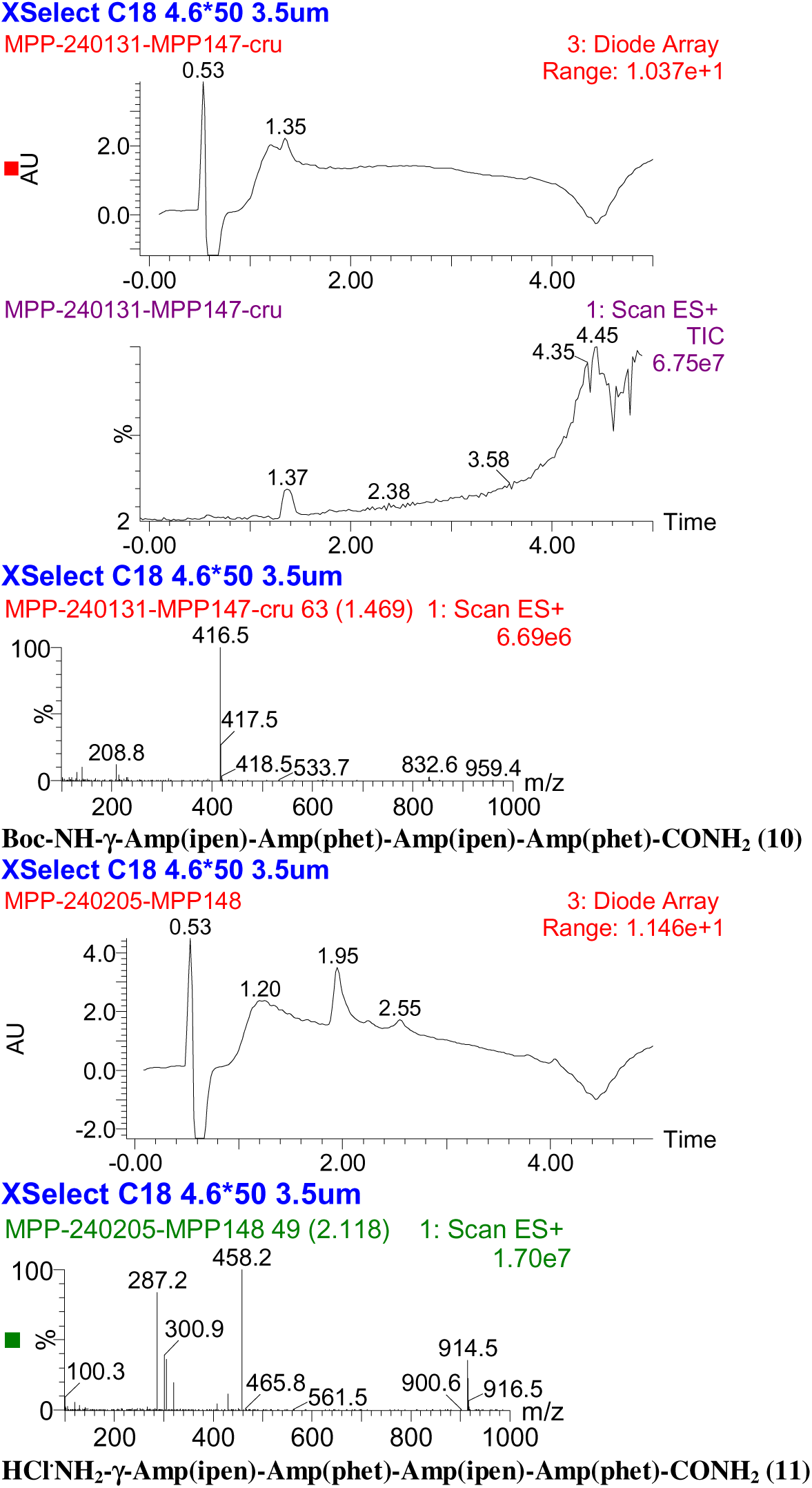

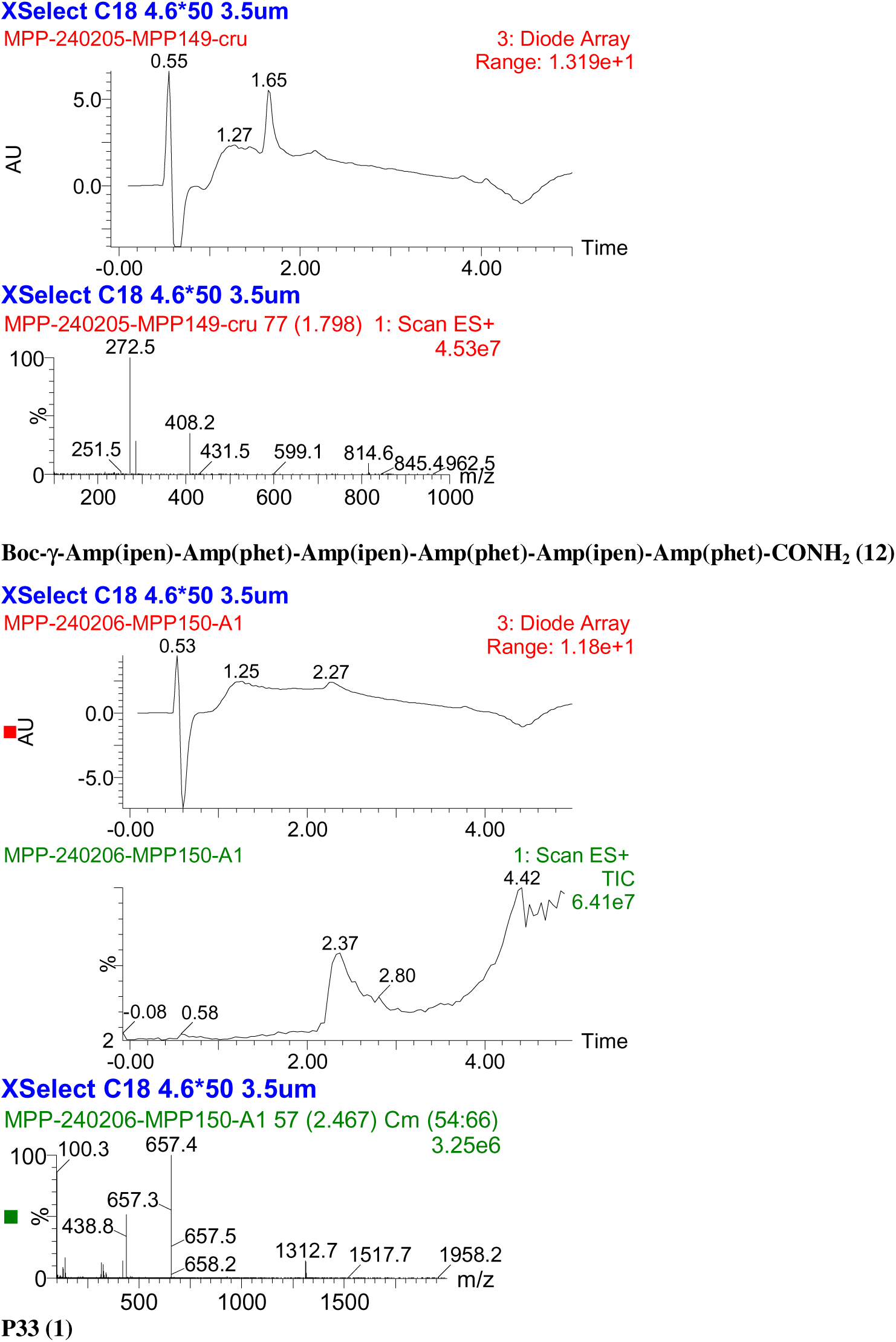

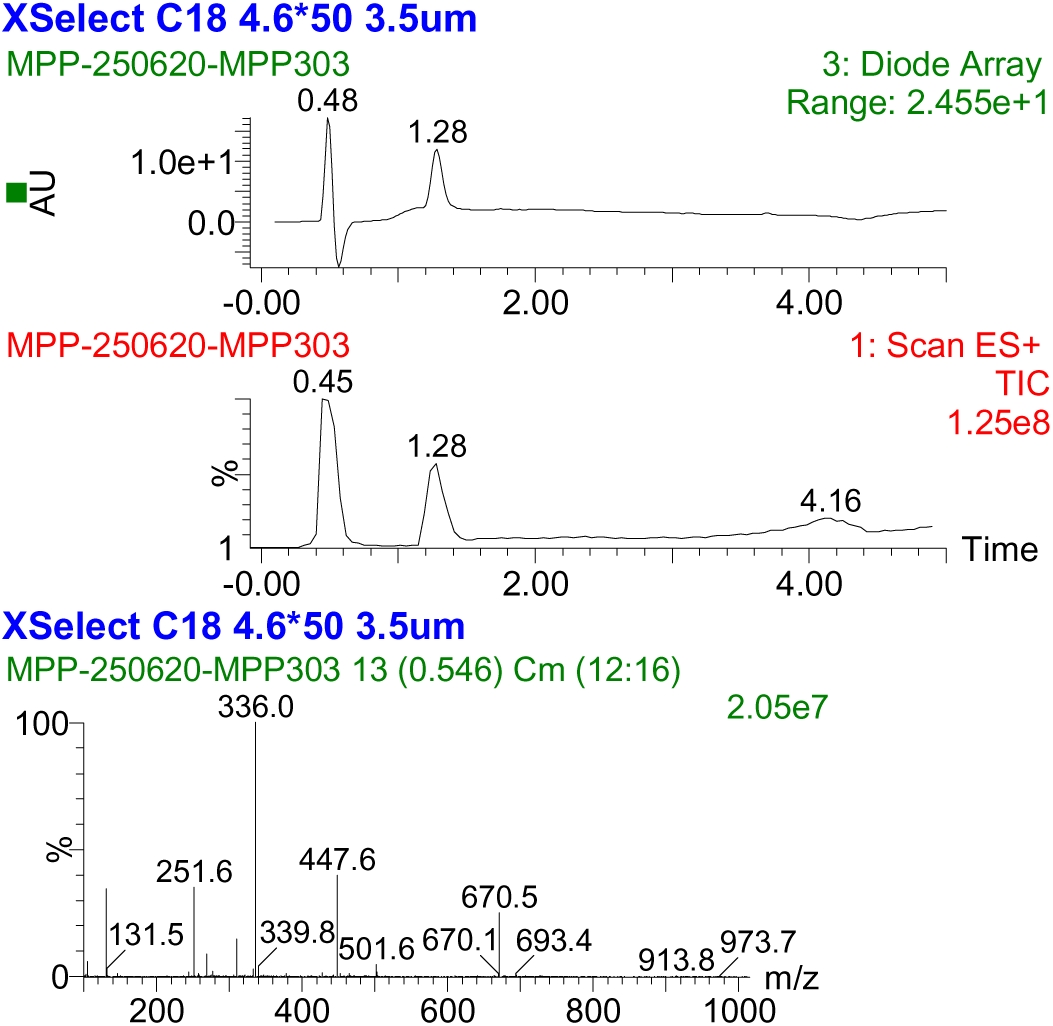

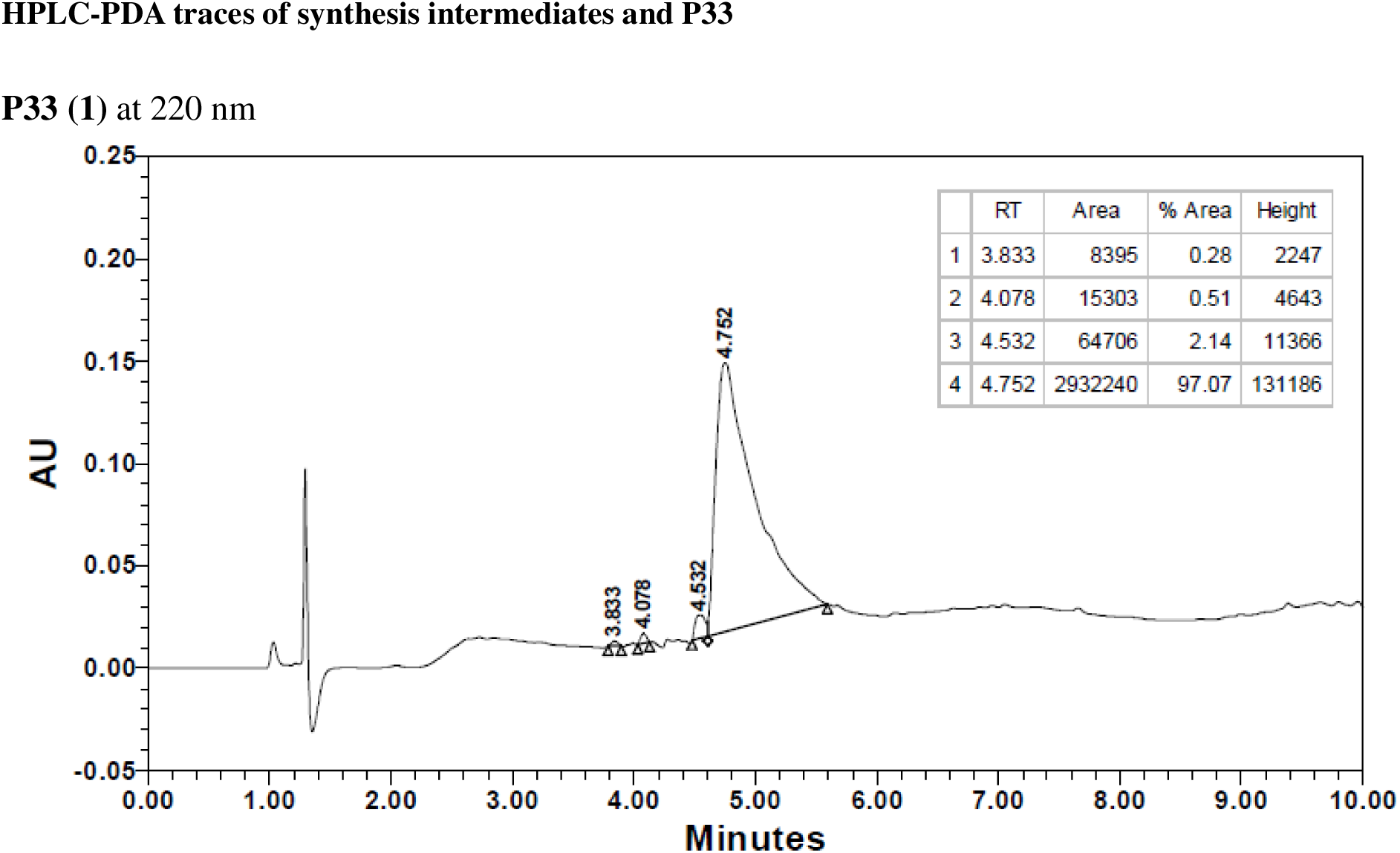

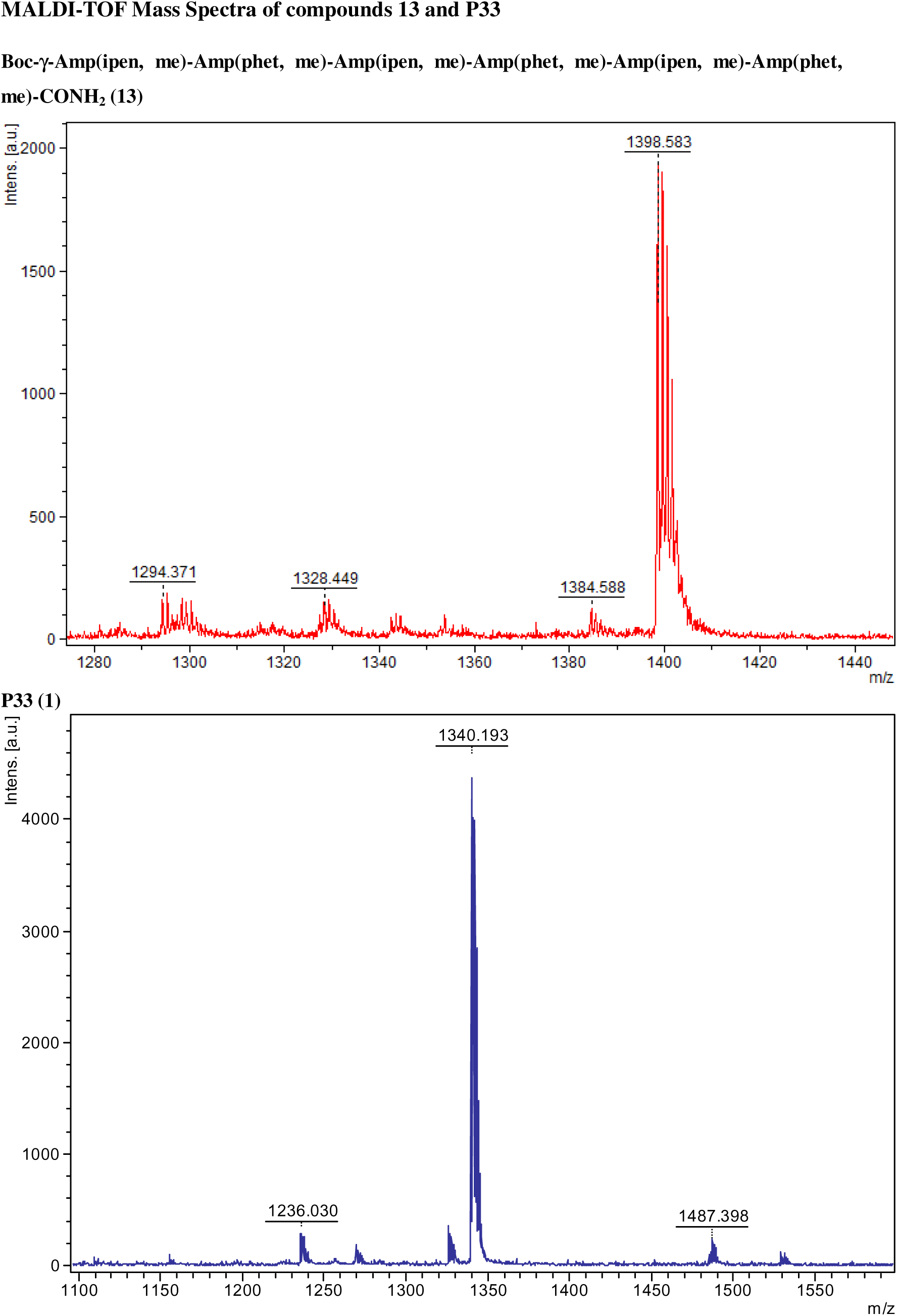

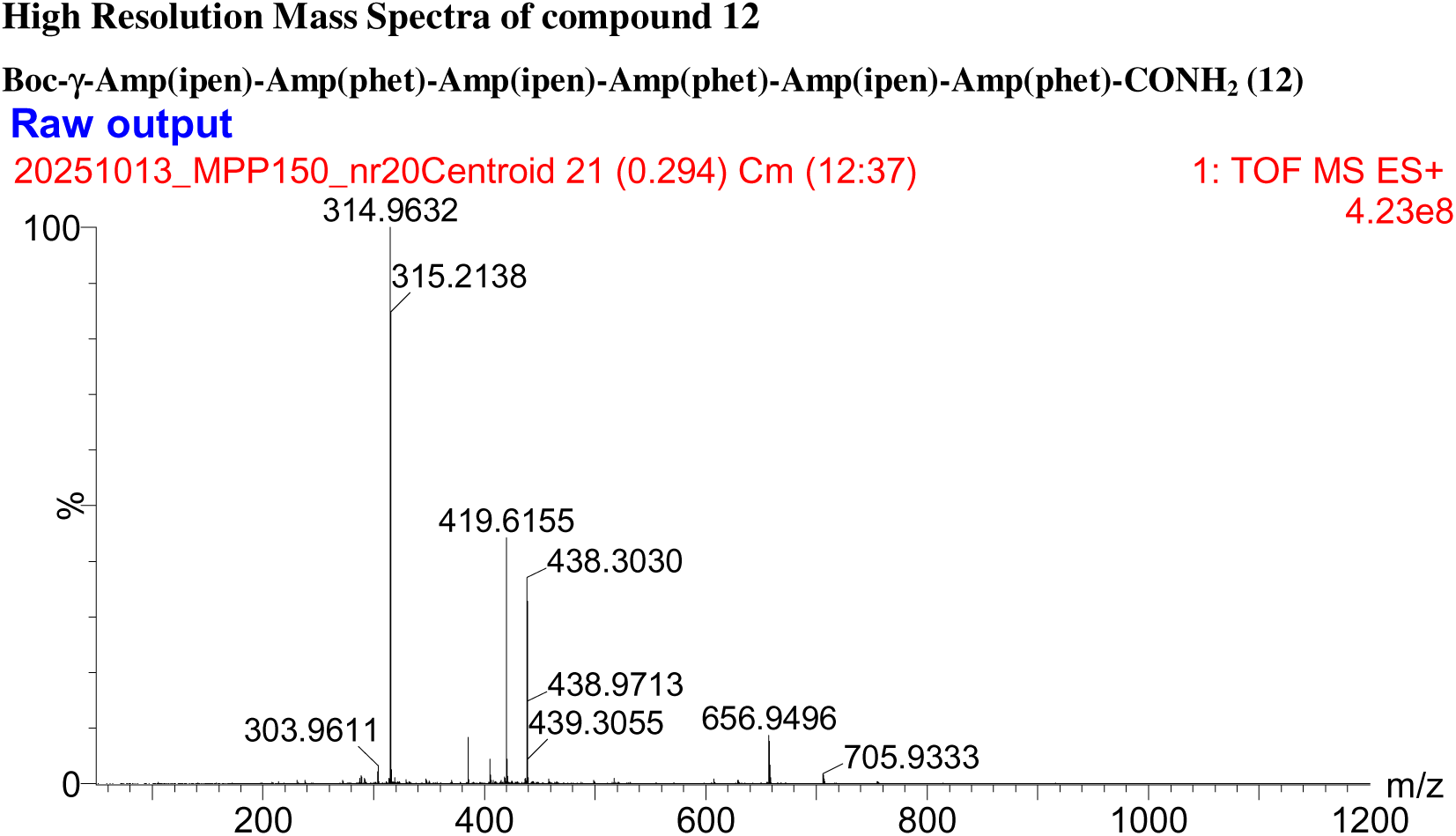

## References

[1] P. Hermann, I. Zerr, Rapidly progressive dementias - aetiologies, diagnosis and management, Nat Rev Neurol 18 (2022) 363–376. 10.1038/s41582-022-00659-0.

[2] A. Schmidt, J. Pahnke, Efficient near-infrared in vivo imaging of amyoid-beta deposits in Alzheimer’s disease mouse models, J Alzheimers Dis 30 (2012) 651–664. 10.3233/JAD-2012-112168.

[3] C. Schmidt, M. Wolff, M. Weitz, T. Bartlau, C. Korth, I. Zerr, Rapidly progressive Alzheimer disease, Arch Neurol 68 (2011) 1124–1130. 10.1001/archneurol.2011.189.

[4] H. Braak, K. Del Tredici, Alzheimer’s pathogenesis: is there neuron-to-neuron propagation?, Acta Neuropathol 121 (2011) 589–595. 10.1007/s00401-011-0825-z.

[5] H. Braak, K. Del Tredici, U. Rub, R.A. de Vos, E.N. Jansen Steur, E. Braak, Staging of brain pathology related to sporadic Parkinson’s disease, Neurobiol. Aging 24 (2003) 197–211. 10.1016/s0197-4580(02)00065-9.

[6] P.T. Nelson, I. Alafuzoff, E.H. Bigio, C. Bouras, H. Braak, N.J. Cairns, R.J. Castellani, B.J. Crain, P. Davies, K. Del Tredici, C. Duyckaerts, M.P. Frosch, V. Haroutunian, P.R. Hof, C.M. Hulette, B.T. Hyman, T. Iwatsubo, K.A. Jellinger, G.A. Jicha, E. Kovari, W.A. Kukull, J.B. Leverenz, S. Love, I.R. Mackenzie, D.M. Mann, E. Masliah, A.C. McKee, T.J. Montine, J.C. Morris, J.A. Schneider, J.A. Sonnen, D.R. Thal, J.Q. Trojanowski, J.C. Troncoso, T. Wisniewski, R.L. Woltjer, T.G. Beach, Correlation of Alzheimer disease neuropathologic changes with cognitive status: a review of the literature, J. Neuropathol. Exp. Neurol. 71 (2012) 362–381. 10.1097/NEN.0b013e31825018f7.

[7] E. Karran, M. Mercken, B. De Strooper, The amyloid cascade hypothesis for Alzheimer’s disease: an appraisal for the development of therapeutics, Nat Rev Drug Discov 10 (2011) 698–712. 10.1038/nrd3505 nrd3505 [pii].

[8] J.L. Cummings, D.P. Goldman, N.R. Simmons-Stern, E. Ponton, The costs of developing treatments for Alzheimer’s disease: A retrospective exploration, Alzheimers Dement 18 (2022) 469–477. 10.1002/alz.12450.

[9] R.A. Sperling, C.R. Jack Jr., P.S. Aisen, Testing the right target and right drug at the right stage, Sci Transl Med 3 (2011) 111cm33. 10.1126/scitranslmed.3002609.

[10] R.A. Sperling, P.S. Aisen, L.A. Beckett, D.A. Bennett, S. Craft, A.M. Fagan, T. Iwatsubo, C.R. Jack Jr., J. Kaye, T.J. Montine, D.C. Park, E.M. Reiman, C.C. Rowe, E. Siemers, Y. Stern, K. Yaffe, M.C. Carrillo, B. Thies, M. Morrison-Bogorad, M. V Wagster, C.H. Phelps, Toward defining the preclinical stages of Alzheimer’s disease: recommendations from the National Institute on Aging-Alzheimer’s Association workgroups on diagnostic guidelines for Alzheimer’s disease, Alzheimers Dement 7 (2011) 280–292. 10.1016/j.jalz.2011.03.003.

[11] N. Khartabil, A. Awaness, Targeting Amyloid Pathology in Early Alzheimer’s: The Promise of Donanemab-Azbt, Pharmacy (Basel) 13 (2025). 10.3390/pharmacy13010023.

[12] N.C. Fox, C. Belder, C. Ballard, H.C. Kales, C. Mummery, P. Caramelli, O. Ciccarelli, K.S. Frederiksen, T. Gomez-Isla, Z. Ismail, C. Paquet, R.C. Petersen, R. Perneczky, L. Robinson, O. Sayin, G.B. Frisoni, Treatment for Alzheimer’s disease, The Lancet 406 (2025) 1408–1423. 10.1016/S0140-6736(25)01329-7.

[13] M.R. Farlow, J.R. Brosch, Immunotherapy for Alzheimer’s disease, Neurol Clin 31 (2013) 869–878. 10.1016/j.ncl.2013.03.012.

[14] R.J. Bateman, Y. Li, E.M. McDade, J.J. Llibre-Guerra, D.B. Clifford, A. Atri, S.L. Mills, A.M. Santacruz, G. Wang, C. Supnet, T.L.S. Benzinger, B.A. Gordon, L. Ibanez, G. Klein, M. Baudler, R.S. Doody, P. Delmar, G.A. Kerchner, T. Bittner, J. Wojtowicz, A. Bonni, P. Fontoura, C. Hofmann, L. Kulic, J. Hassenstab, A.J. Aschenbrenner, R.J. Perrin, C. Cruchaga, A.E. Renton, C. Xiong, A.A. Goate, J.C. Morris, D.M. Holtzman, B.J. Snider, C. Mummery, W.S. Brooks, D. Wallon, S.B. Berman, E. Roberson, C.L. Masters, D.R. Galasko, S. Jayadev, R. Sanchez-Valle, J. Pariente, J. Kinsella, C.H. van Dyck, S. Gauthier, G.-Y.R. Hsiung, M. Masellis, B. Dubois, L.S. Honig, C.R. Jack, A. Daniels, D. Aguillón, R. Allegri, J. Chhatwal, G. Day, N.C. Fox, E. Huey, T. Ikeuchi, M. Jucker, J.-H. Lee, A.I. Levey, J. Levin, F. Lopera, J. Roh, P. Rosa-Neto, P.R. Schofield, Safety and efficacy of long-term gantenerumab treatment in dominantly inherited Alzheimer’s disease: an open-label extension of the phase 2/3 multicentre, randomised, double-blind, placebo-controlled platform DIAN-TU trial, Lancet Neurol. 24 (2025) 316–330. 10.1016/S1474-4422(25)00024-9.

[15] X. Xing, X. Zhang, K. Wang, Z. Wang, Y. Feng, X. Li, Y. Hua, L. Zhang, X. Dong, Post-marketing safety concerns with lecanemab: a pharmacovigilance study based on the FDA Adverse Event Reporting System database, Alzheimers Res Ther 17 (2025) 15. 10.1186/s13195-024-01669-4.

[16] J. Hardy, D.J. Selkoe, The Amyloid Hypothesis of Alzheimer’s Disease: Progress and Problems on the Road to Therapeutics, Science (1979). 297 (2002) 353–356. 10.1126/science.1072994.

[17] P.C. May, B.A. Willis, S.L. Lowe, R.A. Dean, S.A. Monk, P.J. Cocke, J.E. Audia, L.N. Boggs, A.R. Borders, R.A. Brier, D.O. Calligaro, T.A. Day, L. Ereshefsky, J.A. Erickson, H. Gevorkyan, C.R. Gonzales, D.E. James, S.S. Jhee, S.F. Komjathy, L. Li, T.D. Lindstrom, B.M. Mathes, F. Martényi, S.M. Sheehan, S.L. Stout, D.E. Timm, G.M. Vaught, B.M. Watson, L.L. Winneroski, Z. Yang, D.J. Mergott, The Potent BACE1 Inhibitor LY2886721 Elicits Robust Central Aβ Pharmacodynamic Responses in Mice, Dogs, and Humans, The Journal of Neuroscience 35 (2015) 1199–1210. 10.1523/JNEUROSCI.4129-14.2015.

[18] M. Ohno, E.A. Sametsky, L.H. Younkin, H. Oakley, S.G. Younkin, M. Citron, R. Vassar, J.F. Disterhoft, BACE1 Deficiency Rescues Memory Deficits and Cholinergic Dysfunction in a Mouse Model of Alzheimer’s Disease, Neuron 41 (2004) 27–33. 10.1016/S0896-6273(03)00810-9.

[19] M. Timmers, J.R. Streffer, A. Russu, Y. Tominaga, H. Shimizu, A. Shiraishi, K. Tatikola, P. Smekens, A. Börjesson-Hanson, N. Andreasen, J. Matias-Guiu, M. Baquero, M. Boada, I. Tesseur, L. Tritsmans, L. Van Nueten, S. Engelborghs, Pharmacodynamics of atabecestat (JNJ-54861911), an oral BACE1 inhibitor in patients with early Alzheimer’s disease: randomized, double-blind, placebo-controlled study, Alzheimers Res. Ther. 10 (2018) 85. 10.1186/s13195-018-0415-6.

[20] D.K. Lahiri, B. Maloney, J.M. Long, N.H. Greig, Lessons from a BACE1 inhibitor trial: OffLJsite but not off base, Alzheimer’s & Dementia 10 (2014). 10.1016/j.jalz.2013.11.004.

[21] A. Murray, A. Muñiz-García, I. Alić, D. Nižetić, It’s good to know what to BACE the specificity of your inhibitors on, Journal of Clinical Investigation 134 (2024). 10.1172/JCI183677.

[22] E.A. Watkins, R. Vassar, BACE Inhibitor Clinical Trials for Alzheimer’s Disease, Journal of Alzheimer’s Disease 101 (2024) S41–S52. 10.3233/JAD-231258.

[23] M. Lindgren, M. Hällbrink, A. Prochiantz, U. Langel, Cell-penetrating peptides., Trends Pharmacol. Sci. 21 (2000) 99–103. 10.1016/s0165-6147(00)01447-4.

[24] P. Lundberg, U. Langel, A brief introduction to cell-penetrating peptides., J. Mol. Recognit. 16 (2003) 227–33. 10.1002/jmr.630.

[25] S. Zhang, Y. Xu, B. Wang, W. Qiao, D. Liu, Z. Li, Cationic compounds used in lipoplexes and polyplexes for gene delivery., J. Control. Release 100 (2004) 165–80. 10.1016/j.jconrel.2004.08.019.

[26] B.L. Davidson, X.O. Breakefield, Viral vectors for gene delivery to the nervous system., Nat. Rev. Neurosci. 4 (2003) 353–64. 10.1038/nrn1104.

[27] F. Milletti, Cell-penetrating peptides: classes, origin, and current landscape., Drug Discov. Today 17 (2012) 850–60. 10.1016/j.drudis.2012.03.002.

[28] J. Farrera-Sinfreu, E. Giralt, S. Castel, F. Albericio, M. Royo, Cell-penetrating cis-gamma-amino-l-proline-derived peptides., J. Am. Chem. Soc. 127 (2005) 9459–68. 10.1021/ja051648k.

[29] H. Brooks, B. Lebleu, E. Vivès, Tat peptide-mediated cellular delivery: back to basics., Adv. Drug Deliv. Rev. 57 (2005) 559–77. 10.1016/j.addr.2004.12.001.

[30] S. Cavalli, D. Carbajo, M. Acosta, S. Lope-Piedrafita, A.P. Candiota, C. Arús, M. Royo, F. Albericio, Efficient γ-amino-proline-derived cell penetrating peptide–superparamagnetic iron oxide nanoparticle conjugates via aniline-catalyzed oxime chemistry as bimodal imaging nanoagents, Chemical Communications 48 (2012) 5322. 10.1039/c2cc17937g.

[31] S.P. Egusquiaguirre, C. Manguán-García, L. Pintado-Berninches, L. Iarriccio, D. Carbajo, F. Albericio, M. Royo, J.L. Pedraz, R.M. Hernández, R. Perona, M. Igartua, Development of surface modified biodegradable polymeric nanoparticles to deliver GSE24.2 peptide to cells: A promising approach for the treatment of defective telomerase disorders, European Journal of Pharmaceutics and Biopharmaceutics 91 (2015) 91–102. 10.1016/j.ejpb.2015.01.028.

[32] J. Fernández-Carneado, M.J. Kogan, S. Castel, E. Giralt, Potential peptide carriers: amphipathic proline-rich peptides derived from the N-terminal domain of gamma-zein., Angew. Chem. Int. Ed Engl. 43 (2004) 1811–4. 10.1002/anie.200352540.

[33] M. Rizzuti, M. Nizzardo, C. Zanetta, A. Ramirez, S. Corti, Therapeutic applications of the cell-penetrating HIV-1 Tat peptide., Drug Discov. Today 20 (2015) 76–85. 10.1016/j.drudis.2014.09.017.

[34] J.L. Jankowsky, H.H. Slunt, T. Ratovitski, N.A. Jenkins, N.G. Copeland, D.R. Borchelt, Co-expression of multiple transgenes in mouse CNS: a comparison of strategies, Biomol Eng 17 (2001) 157–165. S1389034401000673 [pii].

[35] C. Vergara, L. Ordonez-Gutierrez, F. Wandosell, I. Ferrer, J.A. del Rio, R. Gavin, Role of PrP(C) Expression in Tau Protein Levels and Phosphorylation in Alzheimer’s Disease Evolution, Mol Neurobiol 51 (2015) 1206–1220. 10.1007/s12035-014-8793-7.

[36] R. Cecchelli, S. Aday, E. Sevin, C. Almeida, M. Culot, L. Dehouck, C. Coisne, B. Engelhardt, M.-P. Dehouck, L. Ferreira, A Stable and Reproducible Human Blood-Brain Barrier Model Derived from Hematopoietic Stem Cells, PLoS One 9 (2014) e99733. 10.1371/journal.pone.0099733.

[37] M. Sánchez-Navarro, E. Giralt, M. Teixidó, Blood–brain barrier peptide shuttles, Curr. Opin. Chem. Biol. 38 (2017) 134–140. 10.1016/j.cbpa.2017.04.019.

[38] B. Oller-Salvia, M. Sánchez-Navarro, E. Giralt, M. Teixidó, Blood–brain barrier shuttle peptides: an emerging paradigm for brain delivery, Chem. Soc. Rev. 45 (2016) 4690–4707. 10.1039/C6CS00076B.

[39] V. Selvaraj, M.M. Soundarapandian, O. Chechneva, A.J. Williams, M.K. Sidorov, A.M. Soulika, D.E. Pleasure, W. Deng, PARP-1 deficiency increases the severity of disease in a mouse model of multiple sclerosis., J. Biol. Chem. 284 (2009) 26070–84. 10.1074/jbc.M109.013474.

[40] K.J. Livak, T.D. Schmittgen, Analysis of relative gene expression data using real-time quantitative PCR and the 2(-Delta Delta C(T)) Method, Methods 25 (2001) 402–408. 10.1006/meth.2001.1262S1046-2023(01)91262-9 [pii].

[41] P. Carulla, A. Bribian, A. Rangel, R. Gavin, I. Ferrer, C. Caelles, J.A. Del Rio, F. Llorens, Neuroprotective role of PrPC against kainate-induced epileptic seizures and cell death depends on the modulation of JNK3 activation by GluR6/7-PSD-95 binding, Mol Biol Cell 22 (2011) 3041–3054. 10.1091/mbc.E11-04-0321 mbc.E11-04-0321 [pii].

[42] E. Bello-Arroyo, H. Roque, A. Marcos, J. Orihuel, A. Higuera-Matas, M. Desco, V.R. Caiolfa, E. Ambrosio, E. Lara-Pezzi, M.V. Gómez-Gaviro, MouBeAT: A New and Open Toolbox for Guided Analysis of Behavioral Tests in Mice, Front. Behav. Neurosci. 12 (2018). 10.3389/fnbeh.2018.00201.

[43] O. Seira, R. Gavin, V. Gil, F. Llorens, A. Rangel, E. Soriano, J.A. del Rio, Neurites regrowth of cortical neurons by GSK3beta inhibition independently of Nogo receptor 1, J Neurochem 113 (2010) 1644–1658. 10.1111/j.1471-4159.2010.06726.x JNC6726 [pii].

[44] D.J. Selkoe, Alzheimer’s disease results from the cerebral accumulation and cytotoxicity of amyloid beta-protein, J Alzheimers Dis 3 (2001) 75–80. http://www.ncbi.nlm.nih.gov/pubmed/12214075.

[45] R. Behrendt, P. White, J. Offer, Advances in Fmoc solidLJphase peptide synthesis, Journal of Peptide Science 22 (2016) 4–27. 10.1002/psc.2836.

[46] R. Cecchelli, V. Berezowski, S. Lundquist, M. Culot, M. Renftel, M.-P. Dehouck, L. Fenart, Modelling of the blood–brain barrier in drug discovery and development, Nat. Rev. Drug Discov. 6 (2007) 650–661. 10.1038/nrd2368.

[47] B. OllerLJSalvia, M. Teixidó, E. Giralt, From venoms to BBB shuttles: Synthesis and blood–brain barrier transport assessment of apamin and a nontoxic analog, Peptide Science 100 (2013) 675–686. 10.1002/bip.22257.

[48] D.S. Mortensen, S.M. Perrin-Ninkovic, R. Harris, B.G.S. Lee, G. Shevlin, M. Hickman, G. Khambatta, R.R. Bisonette, K.E. Fultz, S. Sankar, Discovery and SAR exploration of a novel series of imidazo[4,5-b]pyrazin-2-ones as potent and selective mTOR kinase inhibitors, Bioorg. Med. Chem. Lett. 21 (2011) 6793–6799. 10.1016/j.bmcl.2011.09.035.

[49] C.G. Parsons, W. Danysz, A. Dekundy, I. Pulte, Memantine and Cholinesterase Inhibitors: Complementary Mechanisms in the Treatment of Alzheimer’s Disease, Neurotox. Res. 24 (2013) 358–369. 10.1007/s12640-013-9398-z.

[50] B. Grayson, M. Leger, C. Piercy, L. Adamson, M. Harte, J.C. Neill, Assessment of disease-related cognitive impairments using the novel object recognition (NOR) task in rodents, Behavioural Brain Research 285 (2015) 176–193. 10.1016/j.bbr.2014.10.025.

